# Dynamic HSF2 regulation drives breast cancer progression by steering the balance between proliferation and invasion

**DOI:** 10.1101/2024.06.24.600354

**Authors:** Jenny C. Pessa, Oona Paavolainen, Mikael C. Puustinen, Hendrik S. E. Hästbacka, Alejandro J. Da Silva, Sandra Pihlström, Silvia Gramolelli, Pia Boström, Pauliina Hartiala, Emilia Peuhu, Jenny Joutsen, Lea Sistonen

## Abstract

Phenotypic plasticity is a hallmark of breast carcinogenesis that facilitates the acquisition of invasive properties *via* epithelial-mesenchymal transition (EMT). Transforming growth factor-beta (TGF-β), a key EMT-inducing cytokine, drives pro-metastatic gene programs through downstream transcription factors. Among mammalian stress-protective transcription factors, heat shock factor 2 (HSF2) has been associated with cancer progression, but the mechanisms regulating HSF2 expression and activity are unknown. Here, we demonstrate that TGF-β stimulation downregulates HSF2 to enable activation of an invasive phenotype. Remarkably, ectopic expression of HSF2 in breast cancer cells inhibited TGF-β-mediated effects on gene expression and cellular properties. By using both *in vitro* cell models and *in vivo* zebrafish xenografts, we found that a temporal downregulation of HSF2 is a prerequisite for EMT activation, while ectopically sustained HSF2 promotes rapid cell proliferation and survival. Analyses of human patient tissues corroborated our results by showing that HSF2 is dynamically regulated during breast cancer progression. Specifically, HSF2 expression and nuclear co-localization with the proliferation marker Ki67 are dramatically increased already in ductal carcinoma *in situ*. Altogether, our findings expand the pathological landscape of HSF2, demonstrating that dynamic regulation of HSF2 is an inherent property of malignant progression and characterizes the distinct stages of breast cancer.

## INTRODUCTION

Breast cancer is a heterogeneous disease that follows a complex progression path where each stage is defined by unique cellular behaviors and dynamic molecular events. The pre-invasive state of breast cancer, ductal carcinoma *in situ* (DCIS), is characterized by abnormal proliferation of epithelial cells within the breast ducts that are confined from the stroma by myoepithelial cells and an intact basement membrane (Wilson et al., 2022). Although DCIS has a good prognosis, it is also considered as the non-obligate precursor of invasive ductal carcinoma (IDC). Yet the drivers of the invasive transition remain poorly characterized, obscuring the reliable identification of high-risk lesions (Wilson et al., 2022; Wang et al., 2024). In IDC, neoplastic epithelial cells have breached through the basement membrane and evaded the ductal confinement into the surrounding stromal tissue (Lo et al., 2017; Wilson et al., 2022; Wang et al., 2024). The transition from DCIS to IDC is initiated by a subset of transformed cells that unlock the restricted capacity of phenotypic plasticity. The cell plasticity program epithelial-mesenchymal transition (EMT) plays a crucial role during this stage, facilitating the transition of epithelial cells into a more motile mesenchymal phenotype (Dongre & Weinberg, 2019). The gradual changes in cellular characteristics during EMT involve the loss of cell-cell adhesions, disruption of the apico-basal polarity, and the gain of mesenchymal markers, which allow cancer cells to detach from the primary tumor and invade to adjacent tissues (Dongre & Weinberg, 2019). At the early stages of IDC, the invasive cancer cells form microlesions around the ducts that can continue to grow into the stroma and may lead to metastasis. Throughout the different stages of breast cancer progression, dysregulated cell proliferation, driven by genetic alterations and disrupted signaling pathways, fuels tumor expansion. Understanding the interplay between cell proliferation, EMT, and invasion is fundamental for elucidating the mechanisms driving breast cancer progression.

Many signals in the microenvironment, including cytokines and components of the extracellular matrix (ECM), can trigger EMT. Transforming growth factor-beta (TGF-β) has emerged as a pivotal EMT-inducing cytokine, encompassing widespread functions, which guide development, immune modulation, cell fate decisions, and stem cell functions (Dongre & Weinberg, 2019; Lee & Massagué, 2022). Due to its multifaceted roles, disturbances to the TGF-β pathway are strongly associated with human diseases, including cancer (Massagué, 2012; Derynck et al., 2021). Under physiological conditions, TGF-β acts as a tumor suppressor by inhibiting the cell cycle and cell proliferation through regulation of cyclin-dependent kinase activity, and by inducing cell differentiation programs and apoptosis (Principe et al., 2014; Chen et al., 2020). However, mutations in various components of the TGF-β signaling pathway impair its growth inhibitory function, leading to uncontrollable proliferation of both cancer cells and stromal cells. During cancer progression, TGF-β signaling promotes ECM remodeling and cell migration through EMT induction, which enables cancer cell invasion and metastasis (Massagué, 2012; Derynck et al., 2021). In breast cancer, the aberrant activation of EMT orchestrated by TGF-β provides a mechanism underlying the acquisition of aggressive phenotypes and disease dissemination. Activation of TGF-β signaling in breast myoepithelial cells has been implicated in promoting the invasive progression of DCIS to IDC (Lo et al., 2017). The signaling effects of TGF-β are mediated through transmembrane serine/threonine kinase receptors type I and type II, which can activate either a canonical signaling pathway, consisting of SMAD family proteins, or a non-canonical pathway, comprised of other intermediators (Massagué 2012; Lee & Massagué, 2022). Irrespective of the activated pathway, the cellular response to TGF-β is transmitted by specific downstream transcription factors and gene expression programs.

Heat shock factors (HSFs) are the main transcriptional regulators of the evolutionarily conserved heat shock response (Gomez-Pastor et al., 2018; Joutsen & Sistonen, 2019; Pessa et al., 2024). Among the mammalian HSFs, HSF1 is the master stress-responsive transcription factor that can act either independently or synergistically with HSF2 (Kmiecik & Mayer, 2022; Roos-Mattjus & Sistonen, 2022). In addition to their fundamental roles in cell stress, HSFs are involved in multiple physiological and pathological processes, ranging from cell cycle regulation and differentiation to neurodevelopmental diseases and cancer (Riva et al., 2012; Li et al., 2016; Puustinen & Sistonen, 2020; Sala & Morimoto, 2022; Mayer, 2024). Specifically, HSF1 is exploited during tumorigenesis to induce proliferation, survival, and metastasis through cancer-specific transcriptional programs (Mendillo et al., 2012; Scherz-Shouval et al., 2014; Smith et al., 2022). In contrast to HSF1, the role of HSF2 in tumorigenesis is still poorly understood. We have previously identified that HSF2 acts as a tumor suppressor in prostate cancer, where its decreased expression correlates with EMT-associated cell invasion (Björk et al., 2016). Other cancer models have revealed that HSF2 can either promote or suppress cancer cell growth (Mustafa et al., 2010; Zhong et al., 2016; Meng et al., 2017; Yang et al., 2019; Smith et al., 2022). At the cellular level, maintenance of HSF2 expression is required for cell-cell adhesion contacts (Joutsen et al., 2020; de Thonel et al., 2022). To date, no HSF2-specific gene program supporting tumorigenesis has been identified, and the signaling pathways regulating HSF2 expression and its activity during malignant transformation remain to be uncovered. Moreover, since previous research has either employed cell line models or focused on advanced stages of breast cancer, the role of HSF2 in the initial phases of breast cancer progression, including DCIS and its early transitioning to IDC, is not known.

In this study, we explored the dynamic regulation of HSF2 and its functional consequences in breast cancer. To characterize the function of HSF2 in early malignant transformation, we treated human breast epithelial cells with TGF-β to mimic the induction of EMT. TGF-β dramatically downregulated HSF2 levels and activated target genes crucial for the acquisition of pro-metastatic behavior. Intriguingly, ectopic expression of HSF2 disrupted the TGF-β-mediated gene programs by upregulating the expression of cell cycle regulators and downregulating ECM and cell-matrix adhesion-related genes. Accordingly, cells with forced expression of HSF2 displayed induced cell proliferation and reduced migration both *in vitro* and *in vivo*. In healthy human breast tissue, HSF2 was expressed in the cytoplasm of breast epithelial cells, whereas its levels, nuclear expression, and co-localization with Ki67 increased remarkably in the pre-invasive DCIS and in IDC. Our results show that dynamic HSF2 expression steers the balance between cell proliferation and invasion during breast cancer progression. Altogether, this study expands the pathological landscape of HSF2, demonstrating that dynamic regulation of HSF2 levels is an inherent property of the distinct stages of breast cancer.

## RESULTS

### HSF2 is downregulated in response to TGF-β-induced EMT

We have previously shown in cell-based models that downregulation of HSF2 coincides with EMT and destabilization of cell-cell adhesion contacts (Björk et al., 2016; Joutsen et al., 2020; de Thonel et al., 2022). Here, we hypothesized that HSF2 is responsive to EMT-inducing signals and addressed the dynamics and functional consequences of HSF2 regulation during EMT activation in human breast epithelial cells. Transformed (HS578T and MDA-MB-231) and non-transformed (MCF10A) human breast epithelial cells were treated with an EMT-inducing supplement containing recombinant human transforming growth factor-beta 1 (TGF-β_1_, hereafter TGF-β), Wnt family member 5A, antibodies against human E-cadherin, secreted frizzled-related protein 1, and Dickkopf-related protein 1. A 24-h treatment with the EMT supplement (Peinado et al., 2003; Medici et al., 2006) significantly reduced the levels of HSF2 protein in all studied cell lines (Fig. 1A, Fig. S1A). By using a selective small molecule inhibitor of the TGF-β type I receptor, SB431542 (Inman et al., 2002), we demonstrated that the reduction of HSF2 is a specific downstream event of the TGF-β type I receptor activation (Fig. 1A, Fig. S1A).

**Figure 1.**
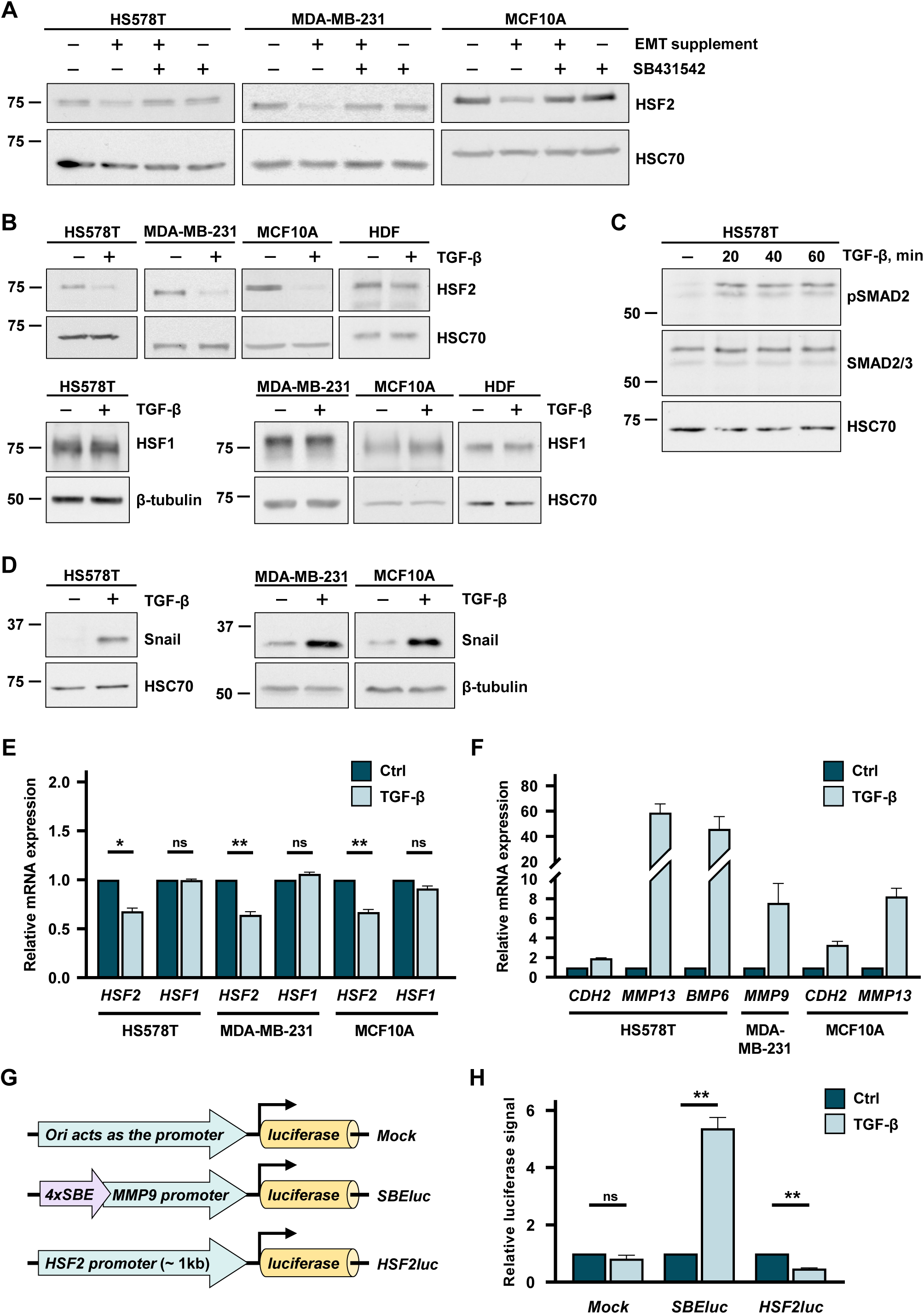
HSF2 is downregulated in response to TGF-β signaling. **(A)** Immunoblot analysis of HSF2 protein levels in human breast epithelial cell lines HS578T, MDA-MB-231 and MCF10A treated with StemXVivo EMT Inducing Media Supplement (EMT supplement) and/or 10 µM ALK5 receptor inhibitor (SB431542) for 24 h. Control cells were treated with assay medium. HSC70 was used as a loading control. **(B)** Immunoblot analysis of HSF2 and HSF1 protein levels in HS578T, MDA-MB-231 and MCF10A cells as well as human dermal fibroblasts (HDF) treated with 10 ng/ml TGF-β_1_ or assay medium for 24 h. HSC70 and β-tubulin were used as a loading control. **(C)** Immunoblot analysis of phosphorylated SMAD2 (pSMAD2) and total SMAD2/3 protein levels in HS578T cells. Cells were treated with assay medium only (-) or with 10 ng/ml TGF-β_1_ for 20, 40 or 60 min. HSC70 was used as a loading control. **(D)** Immunoblot analysis of Snail protein levels in HS578T, MDA-MB-231 and MCF10A cells treated with 10 ng/ml TGF-β_1_ or assay medium for 24 h. HSC70 or β-tubulin were used as loading controls. **(E)** qRT-PCR analysis of *HSF2* and *HSF1* mRNA expression in HS578T, MDA-MB-231 and MCF10A cells treated with 10 ng/ml TGF-β_1_ or assay medium (Ctrl) for 24 h. Relative mRNA quantities were normalized to the housekeeping gene 18S. The mRNA expression is shown relative to the Ctrl sample. Results were plotted as mean ± SEM, * = p-value ≤ 0.05, ** = p-value ≤ 0.01, ns = not significant. All data represents three biological replicates. **(F)** qRT-PCR analysis of the mRNA expression of selected genes associated with the TGF-β signaling pathway: *CDH2*, *MMP13*, and *BMP6* in HS578T cells; *MMP9* in MDA-MB-231 cells; and *CDH2* and *MMP13* in MCF10A cells treated with 10 ng/ml TGF-β_1_ or assay medium (Ctrl) for 24 h. Relative mRNA quantities were normalized to the housekeeping gene 18S. The mRNA expression is shown relative to the Ctrl sample. All data represents three biological replicates. **(G)** A schematic illustration of plasmids used for luciferase reporter assay. The negative control (*Mock*) is an empty plasmid with origin of replication (Ori) driving the basal transcription of the luciferase gene (Dobin et al., 2013). The positive control, *SBEluc,* contains four SMAD-binding elements (SBEs) upstream of the *MMP9* promoter. The *HSF2luc* plasmid contains 1 kb of the *HSF2* promoter. **(H)** Analysis of the luciferase reporter gene activity in HS578T cells co-transfected with plasmids encoding β-galactosidase and *Mock*, *SBEluc*, or *HSF2luc.* Transfected cells were treated with 10 ng/ml TGF-β_1_ or assay medium (Ctrl) for 24 h. The luciferase signal was normalized to β-galactosidase activity and is shown relative to the Ctrl sample. Results were plotted as mean ± SEM, ** = p-value ≤ 0.01, ns = not significant. All data represents three biological replicates.

Upon EMT, tumor cells lose their epithelial characteristics and acquire a mesenchymal phenotype. To investigate whether the ability of TGF-β to downregulate HSF2 is a specific feature of the epithelial phenotype, we treated mesenchymal human dermal fibroblasts (HDFs) with TGF-β. Intriguingly, HSF2 was not affected by TGF-β in HDFs, but a robust downregulation was observed in human breast epithelial cells (Fig. 1B, Fig. S1B), indicating that the TGF-β-induced downregulation of HSF2 is a unique for cells of epithelial origin. In sharp contrast to HSF2, the protein levels of HSF1 remained unchanged in all examined cell lines (Fig. 1B, Fig. S1C). By treating MDA-MB-231 HSF1 knock-out cells (HSF1-KO) (Smith et al., 2022) with TGF-β, we confirmed that HSF2 is downregulated in response to the treatment irrespective of HSF1 (Fig. S2A). The efficacy of TGF-β treatment was confirmed by analyzing the phosphorylation status of SMAD2 (Feng & Derynck, 2005) and the protein levels of the EMT marker Snail (Peinado et al., 2003). Already a 20-min treatment caused a prominent phosphorylation of SMAD2 (Fig. 1C), and Snail was substantially upregulated in response to a 24-h treatment (Fig. 1D; Fig. S1C), showing that the selected TGF-β treatment was sufficient to activate the signaling pathway.

We next addressed whether the TGF-β-mediated downregulation of HSF2 is a consequence of its reduced gene expression. The qRT-PCR analysis revealed that *HSF2* mRNA was significantly decreased in TGF-β-treated breast epithelial cells, but *HSF1* mRNA expression remained stable (Fig. 1E). Several known TGF-β target genes, such as *CDH2*, *MMP13*, *BMP6*, and *MMP9*, were measured to ensure that the cells responded to the treatment, and in line with previous studies (Peinado et al., 2003; Jordà et al., 2005; Moreno-Bueno et al., 2006; Xu et al., 2009; Kim et al., 2020), their expression was induced by TGF-β (Fig. 1F). To investigate if TGF-β decreased HSF2 expression by repressing the HSF2 promoter, we generated a plasmid containing the luciferase reporter gene, driven by 1 kb of the *HSF2* promoter (*HSF2luc*, Fig. 1G). An empty plasmid, with the origin of replication driving the basal transcription of the luciferase gene (Dobin et al., 2013), was used as a negative control (*Mock*, Fig. 1G), and a chimeric construct containing 1 kb of the *MMP9* promoter with four SMAD-binding elements was used as a positive control (*SBEluc*, Fig. 1G). HS578T cells were transfected with β-galactosidase together with either *Mock*, *SBEluc*, or *HSF2luc* and treated with TGF-β for 24 h. As expected, the luciferase signal from *Mock*-transfected cells remained unchanged, and the cells expressing *SBEluc* displayed a significant induction in response to the TGF-β treatment (Fig. 1H). Remarkably, in cells transfected with *HSF2luc*, the luciferase activity declined after exposure to TGF-β (Fig. 1H), demonstrating that the TGF-β-mediated downregulation of HSF2 occurs at the transcriptional level. Since TGF-β can regulate the *trans*-activating capacity of HSF1 (Sasaki et al., 2002), and HSF1 can directly bind to the promoter of HSF2 and regulate its transcriptional activity (Santopolo et al., 2021), we determined if HSF1 participates in the TGF-β-mediated downregulation of HSF2 at the promoter level. HS578T cells were co-transfected with scrambled siRNAs (Scr) or siRNAs targeting HSF1 (siHSF1), β-galactosidase and *HSF2luc* plasmids. The efficacy of HSF1 silencing was confirmed by immunoblotting (Fig. S2B, right), and the luciferase activity was highly similar in the presence and absence of HSF1 (Fig. S2B, left). Together this data shows that TGF-β signaling specifically downregulates HSF2.

### Forced HSF2 expression disrupts TGF-β-regulated gene programs

Downregulation of HSF2 upon TGF**-**β treatment suggests that HSF2 is one of the downstream effector transcription factors of the TGF**-**β signaling pathway. We examined the TGF-β-induced gene programs that depend on HSF2 downregulation and whether forced expression of HSF2 would disrupt this gene expression signature. HS578T cells were transiently transfected with plasmids encoding either GFP (Mock^T^) or exogenous HSF2 (HSF2oe^T^). The transient expression of HSF2 was sufficient to override the downregulating effect of TGF-β on HSF2 (Fig. 2A). The global gene expression profiles of Mock^T^ and HSF2oe^T^ cells were analyzed by RNA-seq in the presence and absence of TGF-β stimulation. In Mock^T^ cells, the treatment resulted in upregulation of well-known TGF-β target genes, *IL11*, *SNAI1*, *WNT1*, and *CDH2* (N-cadherin) (Fig. S3), confirming the functionality of our treatment. Next, we addressed the specific role of HSF2 in the TGF-β signaling pathway by identifying the TGF-β-responsive genes that are dependent on the loss of HSF2 (Fig. 2B). A set of 131 genes lost their ability to respond to TGF-β in the presence of HSF2 (Fig. 2C, Fig. S4). These genes were divided into four different categories based on their expression pattern (Fig. 2D, Data S1): Group I, genes upregulated by both TGF-β and exogenous HSF2; Group II, genes upregulated by TGF-β but not in the presence of exogenous HSF2; Group III, genes downregulated by both TGF-β and exogenous HSF2; Group IV, genes downregulated by TGF-β but not in the presence of exogenous HSF2.

**Figure 2.**
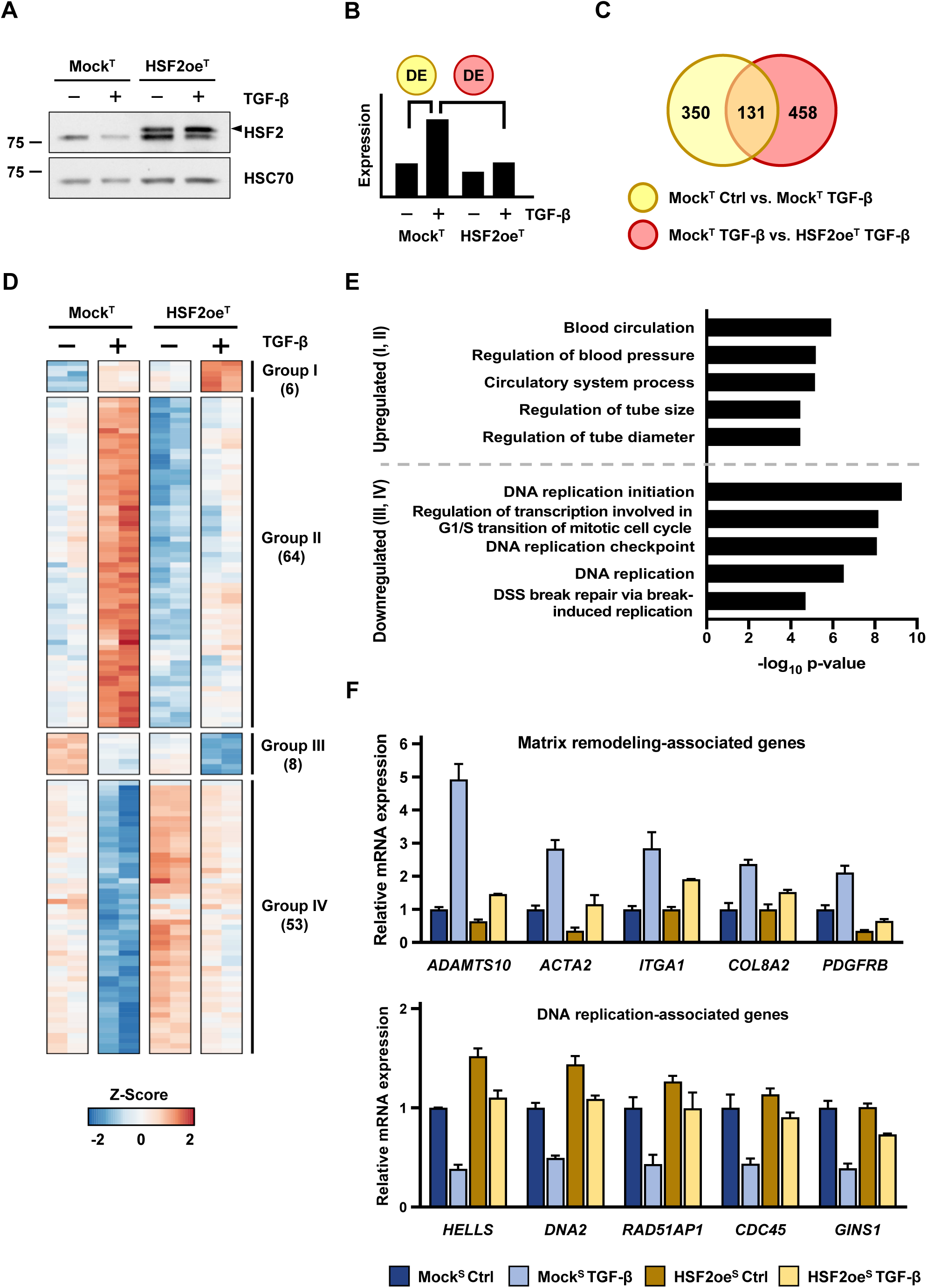
Forced HSF2 expression disrupts the regulation of a subset of TGF-β target genes. **(A)** Immunoblot analysis of HSF2 protein levels in HS578T cells transiently transfected with plasmids encoding GFP (Mock^T^) or HSF2 (HSF2oe^T^). Cells were treated with 10 ng/ml TGF-β_1_ or assay medium for 24 h. Arrowhead (◄) denotes exogenous HSF2. HSC70 was used as a loading control. **(B)** Schematic representation of the pairwise comparisons performed on the RNA-seq data used to identify differentially expressed (DE) genes that respond to TGF-β in cells expressing exogenous HSF2. **(C)** Venn diagram of DE genes. The overlapping region represents a subset of 131 TGF-β target genes, whose expression was impaired by HSF2 overexpression. **(D)** Heat map of the 131 DE genes identified in (C) and categorized into four groups. Group I: genes upregulated by both TGF-β and exogenous HSF2, Group II: genes upregulated by TGF-β but impaired response in the presence of exogenous HSF2, Group III: genes downregulated by both TGF-β and exogenous HSF2, and Group IV: genes downregulated by TGF-β and impaired response in the presence of exogenous HSF2. The number of genes in each group is indicated in brackets. The Z-score was calculated based on the log₂CPM values for each gene. The double columns for each condition represent the two biological replicates from the RNA-seq. **(E)** GO term analysis of the 131 DE genes identified in (C) and categorized in (D). The five most significant biological processes are shown. The analysis was performed with topGO. **(F)** Relative mRNA expression of a subset of the 131 DE genes associated with matrix remodeling and DNA replication. The relative difference in CPM between conditions was normalized to the CPM of the Mock^T^ Ctrl for each gene. The data is presented as mean values ± SEM from two biological replicates.

The assigned groups were subjected to gene ontology (GO) term analysis, which showed that Group I and II genes were associated with vascular functions and ECM remodeling (Fig. 2E, upper panel), and Group III and IV with DNA replication and cell cycle regulation (Fig. 2E, lower panel). As examples of Group I and II genes, *ADAMTS10*, *ACTA2*, *ITGA1*, *COL8A2*, and *PDGFRB*, were markedly increased in control cells treated with TGF-β, and their upregulation was abolished in cells expressing exogenous HSF2 (Fig. 2F, upper panel). Interestingly, ADAM metallopeptidase with thrombospondin type 1 motif 10 (ADAMTS10) is a regulator of ECM composition (Kutz et al., 2011; Cain et al., 2016), and actin alpha 2 (ACTA2) affects the cellular ability to form cell-cell and cell-matrix adhesions (Shinde et al., 2017; Massett et al., 2020). Extensive remodeling of the ECM and cell adhesion contacts are hallmarks of EMT-induced invasion and required for cancer progression. In addition, the transmembrane receptor integrin subunit alpha 1 (ITGA1) is strongly connected to TGF-β-induced EMT and metastasis (Gharibi et al., 2017). These results demonstrate that ectopic HSF2 specifically interfered with the expression of genes characteristic for TGF-β-induced EMT.

As examples of Group III and IV genes, *HELLS*, *DNA2*, *RAD51AP1*, *CDC45*, and *GINS1*, were decreased in TGF-β-treated control cells and the downregulation was inhibited by overexpression of HSF2 (Fig. 2F, lower panel). Cell division cycle 45 (CDC45) and proliferating nuclear antigen (PCNA) proteins are critical for DNA replication initiation and progression, respectively (Strzalka & Ziemienowicz, 2011; Petojevic et al., 2015). It has previously been shown that TGF-β inhibits the loading of CDC45 and PCNA onto chromatin, thereby preventing the activation of minichromosome maintenance complex (MCM), a DNA helicase vital for DNA replication (Mukherjee et al., 2010). The TGF-β-mediated inhibition disturbs the assembly of both pre-replication complexes (Pre-RCs) and pre-initiation complexes (Pre-IRs), which leads to cell cycle arrest at G1/S (Lei, 2005). Taken together, our gene program analyses provide evidence that the TGF-β-mediated downregulation of HSF2 is required to activate gene programs that promote EMT and repress cell proliferation.

### Ectopic HSF2 interferes with the expression of ECM and cell matrix receptor proteins

TGF-β is well known for its multifaceted roles in the regulation of ECM composition and cell adhesion (Xu et al., 2009). Thus, we expanded our gene expression analyses to include more components of the ECM and cell adhesion receptors, which revealed that a comprehensive selection of the genes was unable to respond to TGF-β in the presence of exogenous HSF2 (Fig. 3A). To investigate if the observed changes in mRNA expression were also detected on protein level, we performed immunoblotting of selected targets. Due to slower changes in the protein levels, we extended the TGF-β treatment from 24 h to 72 h. As transiently expressed HSF2 declined already at 72 h (Fig. S5A), stable HS578T cell lines expressing either GFP (Mock^S^) or HSF2 (HSF2oe^S^) were generated (Fig. S5B). Mock^S^ cells maintained their responsiveness to TGF-β stimulation and HSF2oe^S^ cells displayed sustained expression of HSF2 upon a 72-h TGF-β treatment (Fig. S5C). Two ECM proteins, i.e. collagen type III alpha chain 1 (COL3A1) and matrix metalloproteinase 2 (MMP2), were markedly increased in Mock^S^ cells treated with TGF-β for 72 h, but only low expression was observed in HSF2oe^S^ cells (Fig. 3B; Fig. S6A). Both proteins are direct TGF-β targets, and their elevated expression promotes tumorigenesis *via* enhanced ECM remodeling in a variety of cancers (Kuivaniemi & Tromp, 2019; Quintero-Fabián et al., 2019). Similarly, the integrin subunit alpha 1 (ITGA1) protein was dramatically upregulated in Mock^S^ cells but remained unchanged in HSF2oe^S^ cells (Fig. 3B; Fig. S6A). In contrast, the amount of cell-cell adhesion receptor, E-cadherin (CDH1), a classical epithelial state marker (Dongre & Weinberg, 2019), was induced in HSF2oe^S^ cells in comparison to Mock^S^ cells (Fig. 3B; Fig. S6A). Moreover, immunofluorescent staining highlighted the diminished expression of COL3A1 and integrin subunit beta 1 (ITGB1) in HSF2oe^S^ cells (Fig. 3C).

**Figure 3.**
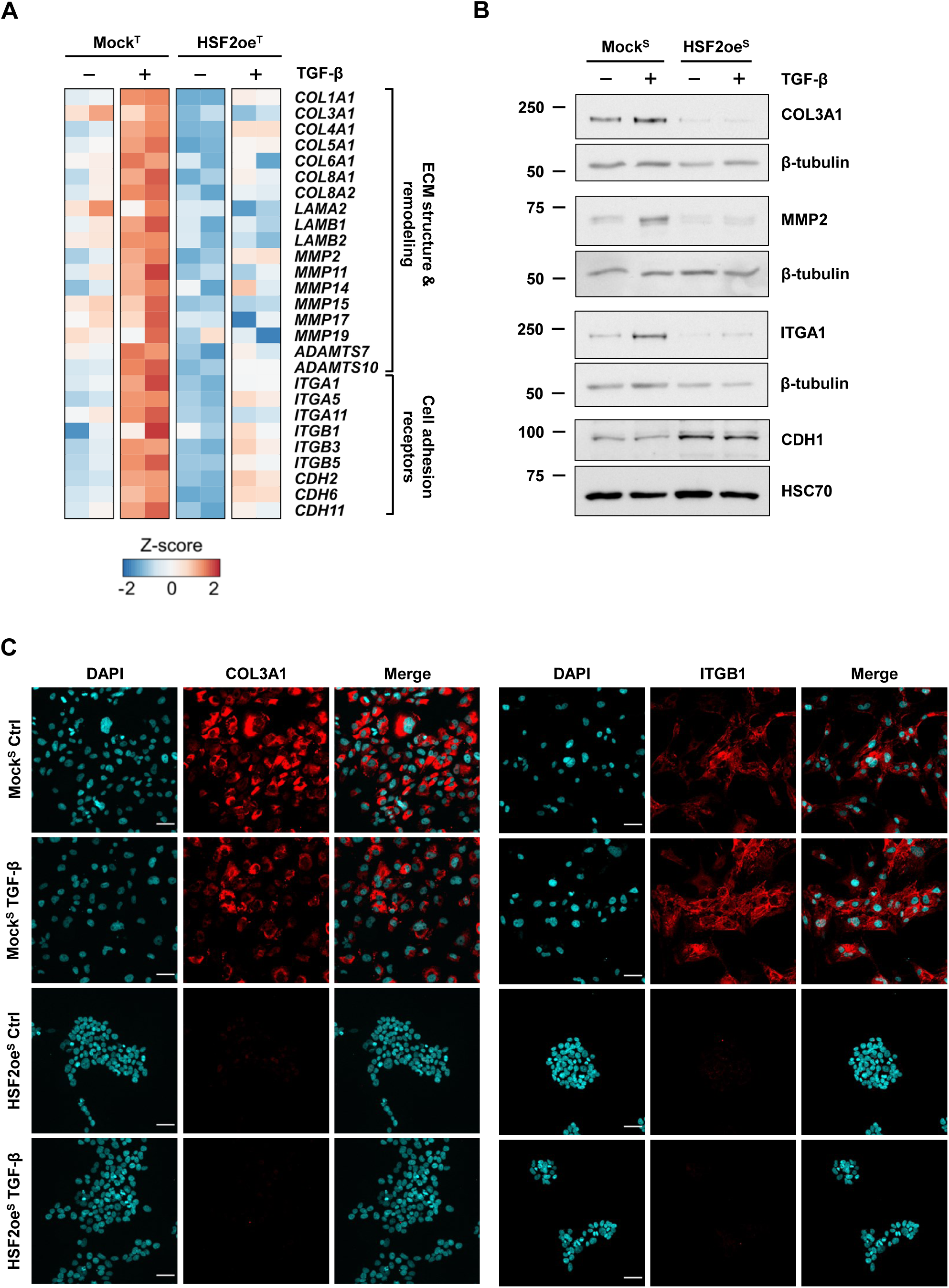
ECM and cell-matrix receptor proteins are downregulated by HSF2. **(A)** Heat map illustrating the expression changes of genes associated with ECM structure and remodeling as well as cell adhesion receptors in HS578T cells transiently transfected with plasmids encoding GFP (Mock^T^) or HSF2 (HSF2oe^T^). Cells were treated with 10 ng/ml TGF-β_1_ or assay medium for 24 h. The Z-score was calculated based on the log₂CPM values for each gene. The double columns for each condition represent the two biological replicates from the RNA-seq. **(B)** Immunoblot analysis of COL3A1, MMP2, ITGA1, and CDH1 protein levels in stable cell lines, Mock^S^ and HSF2oe^S^. Cells were treated with 10 ng/ml TGF-β_1_ or assay medium for 72 h. β-tubulin and HSC70 were used as loading controls. **(C)** Immunofluorescent staining of COL3A1 and ITGB1 in Mock^S^ and HSF2oe^S^ cells ± 10 ng/ml TGF-β_1_ for 24 h. DAPI was used as a nuclear marker. Scale bar 50 µm. Data represents two biological replicates.

Integrins mediate transmembrane crosstalk between the ECM and the cytoskeleton, modulating the linkage based on intracellular and extracellular cues (Kanchanawong & Calderwood, 2023). To investigate how the cytoskeleton was affected by altered expression of ECM and cell adhesion regulators (Fig. 2F; Fig. 3A,B), we analyzed filamentous actin (F-actin), which is critical for integrin-mediated cell adhesions (Svitkina, 2018), and the intermediate filament vimentin, which regulates migratory and mechanical properties of the cell (Ridge et al., 2022). Immunofluorescence staining demonstrated a decreased expression of F-actin and a dramatically altered subcellular localization of vimentin in HSF2oe^S^ cells (Fig. 4A), suggesting that HSF2 modulates organization of the cytoskeleton through suppression of specific ECM and cell-matrix adhesion proteins.

**Figure 4.**
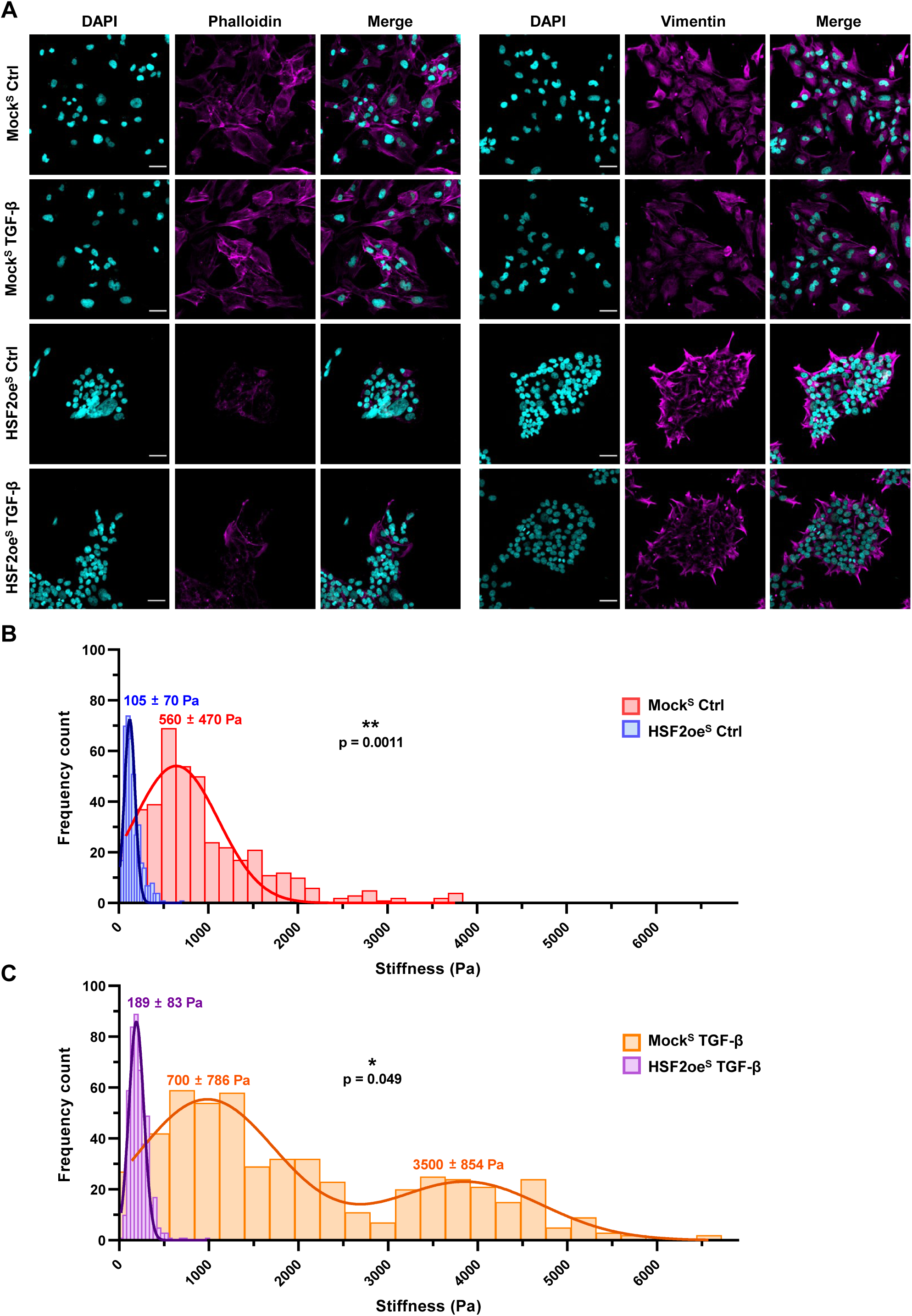
Cellular stiffness is decreased by forced expression of HSF2. **(A)** Immunofluorescent staining of F-actin (Phalloidin) and vimentin in Mock^S^ and HSF2oe^S^ cells ± 10 ng/ml TGF-β_1_ for 24 h. DAPI was used as a nuclear marker. Scale bar 50 µm. Data represents two biological replicates. **(B)** Atomic force microscopy (AFM) indentation analysis of Mock^S^ Ctrl and HSF2oe^S^ Ctrl cell elasticity. Data represents three biological replicates with 414 indentation curves for Mock^S^ (bin size 160 Pa) and 400 for HSF2oe^S^ (bin size 30 Pa). Data is fitted on the same scale to enable comparison between samples. Number of bins = 24. Peak values ± SD are shown. The fitted curves represent a Gaussian distribution. Nested two-tailed t-test, ** = p-value ≤ 0.01. Data represents three biological replicates. **(C)** AFM indentation analysis of Mock^S^ and HSF2oe^S^ cells treated with 10 ng/ml TGF-β_1_. Data represents three biological replicates with 524 indentation curves for Mock^S^ TGF-β (bin size 280 Pa) and 443 for HSF2oe^S^ TGF-β (bin size 42 Pa). Data is fitted on the same scale to enable comparison between samples. Number of bins = 24. Peak values ± SD are shown. The fitted curves represent a Gaussian distribution or a sum of two Gaussians. Nested two-tailed t-test, * = p-value ≤ 0.05. Data represents three biological replicates.

Reorganization of the cytoskeleton is directly coupled to cell mechanics (Deville & Cordes, 2019). We analyzed cellular elasticity with atomic force microscopy (AFM) indentation measurements (Fig. S7A), predicting a Hertz model of impact (Fig. S7B,C). Intriguingly, HSF2oe^S^ cells displayed significantly lower stiffness (peak value 105 Pa) in comparison to Mock^S^ cells (peak value 560 Pa) (Fig. 4B). In contrast, the TGF-β treatment increased the stiffness of Mock^S^ cells (peak values 700 Pa and 3500 Pa). Interestingly, only a minor change was observed in HSF2oe^S^ cells (peak value 189 Pa) (Fig. 4C). Although transformed cells are typically considered softer than non-transformed cells, TGF-β stimulation was recently shown to induce cytoskeletal remodeling that mediated elevation of cell stiffness and invasiveness in non-small-cell lung carcinoma (Gladilin et al., 2019). This emphasizes the cell-type specificity in how cellular elasticity is regulated during metastatic transformation. Altogether, we conclude that by counteracting the TGF-β-induced activation of ECM and cell-matrix adhesion-associated genes, and by inducing cytoskeletal changes, HSF2 modulates the mechanical properties of human breast cancer cells.

### Exogenous HSF2 counteracts TGF-β-induced cell migration and stabilizes cell-cell adhesions

Since alterations in cellular elasticity can affect the migratory properties of cancer cells (Kashani & Packirisamy, 2020), we explored the functional impact of HSF2 on cell migration and invasion. First, *in vitro* vasculogenic mimicry assay was used to study the ability of Mock^S^ and HSF2oe^S^ cells to modulate the ECM and reorganize into tube-like structures in 3D, in the presence and absence of TGF-β (Fig. 5A; Fig. S8A). TGF-β treatment augmented cellular network formation only in Mock^S^ cells (Fig. 5A). The resulting networks were more complex and interconnected than in untreated cells, as indicated by several parameters (Fig. S8B), including increased mean mesh size, number of segments and branching interval (Fig. 5B). The impaired network formation in HSF2oe^S^ cells (Fig. 5A,B), and specifically the difference in mean mesh size and number of segments between TGF-β-treated Mock^S^ and HSF2oe^S^ cells, establish that HSF2 attenuates the cellular ability to modify ECM in response to TGF-β signaling.

**Figure 5.**
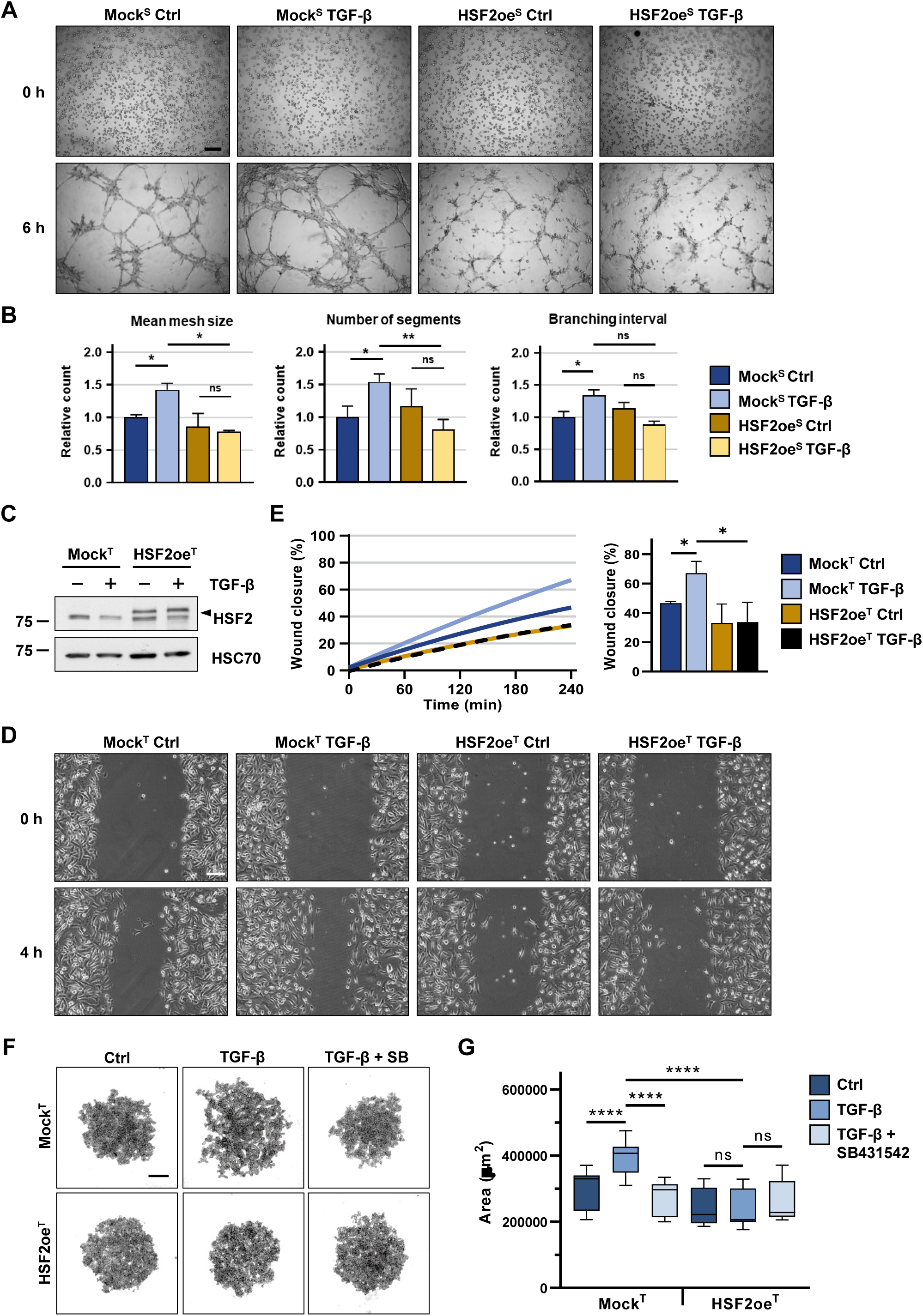
Ectopically expressed HSF2 counteracts TGF-β-induced cell migration and stabilizes cell-cell adhesions. **(A)** Representative images of capillary-like structures in *in vitro* vasculogenic mimicry assay. Mock^S^ and HSF2oe^S^ cells were treated with 10 ng/ml TGF-β_1_ or assay medium (Ctrl) for 24 h prior to initiation of the assay. Following the pre-treatment, 4 × 10^4^ cells were seeded in Matrigel-coated wells of a 96-well plate and treated with 10 ng/ml TGF-β_1_ or assay medium (Ctrl). Each well was imaged at 0 and 6 h with Zeiss Axio Vert. A1 microscope using a 5x objective, NA 0.4. Scale bar 200 µm. **(B)** The capillary-like network formation was analyzed with ImageJ software (version: 1.53f51) using the Angiogenesis Analyzer Toolbox plugin (see Fig. S8). The relative count of each parameter was normalized to the Mock^S^ Ctrl. Results are presented as mean ± SEM, * = p-value < 0.05, ns = not significant. The data represents three biological replicates. **(C)** Immunoblot analysis of HSF2 protein levels in MDA-MB-231 cells transiently transfected with plasmids encoding GFP (Mock^T^) or HSF2 (HSF2oe^T^). Cells were treated with 10 ng/ml TGF-β_1_ or assay medium for 24 h. Arrowhead (◄) denotes exogenous HSF2. HSC70 was used as a loading control. **(D)** Representative images of wound healing assay taken at 0 and 4 h after the scratch. Mock^T^ and HSF2oe^T^ cells were treated with 10 ng/ml TGF-β_1_ or assay medium (Ctrl). The treatments were started 3 h prior to assay initiation to ensure activation of the TGF-β signaling pathway. The rate of wound closure was monitored with live-cell imaging using a Cell-IQ machine with a 5-min frame interval. Scale bar 100 µm. **(E)** Quantitative analysis of wound healing assay. All images were analyzed with ImageJ software (version: 1.53f51) using the MRI wound healing tool. The percentage of wound closure was calculated by normalizing the wound area values of each time point to the initial scratch area. Results were plotted as mean ± SD, * = p-value ≤ 0.05. The data represents three biological replicates. **(F)** Representative images of spheroid-like structure formation using Ultra-Low Attachment (ULA) assay. Mock^T^ and HSF2oe^T^ were incubated in assay medium (Ctrl) or treated with 10 ng/ml TGF-β_1_ alone or in combination with 10 µM TGF-β type I receptor inhibitor (SB431542) for 24 h on ULA plates. Spheroid-like structures were imaged with Zeiss Axio Vert. A1 microscope using a 5x objective, NA 0.4. Scale bar 200 µm. **(G)** Quantitative analysis of spheroid size. The relative area of each spheroid was obtained by analyzing the images with ImageJ software (version: 1.53f51) and the Biotoxin Toolbox plugin. The average area between the biological replicates was calculated, and the relative area for each condition was normalized to the average area of the Mock^T^ Ctrl. The area is shown relative to the Ctrl sample. Results were plotted as mean ± SEM, * = p-value ≤ 0.05. The data represents three biological replicates.

Using a wound healing assay, we investigated if the downregulation of HSF2 in response to TGF-β is required to induce cell migration. MDA-MB-231 cells, with high migratory capacity (Liu et al., 2019), were transfected with either HSF2oe^T^ or Mock^T^ plasmids, and the expression of exogenous HSF2 was verified by immunoblotting (Fig. 5C). TGF-β was added 3 h prior to initiation of the assay to ensure that the cells had activated the TGF-β signaling pathway. Analysis of the relative wound closure revealed that the TGF-β treatment stimulated migration of Mock^T^ cells, whereas HSF2oe^T^ cells displayed a substantial reduction in cell motility (Fig. 5D,E). These results indicate that the loss of HSF2 is required for TGF-β-induced cell migration in breast cancer cells.

Deterioration of cell-cell adhesion contacts is a hallmark of TGF-β-induced EMT (Dongre & Weinberg, 2019). We have previously shown that the loss of HSF2 leads to disrupted cell-cell adhesion (Joutsen et al., 2020; de Thonel et al., 2022), and that HSF2 localizes at cell-cell adhesion sites in human tissues (Joutsen et al., 2024). Prompted by these findings, we examined whether the TGF-β-mediated decrease of HSF2 influenced cell-cell adhesion of breast cancer cells and if the functional consequences could be reversed by overexpression of HSF2. The capacity of MDA-MB-231 Mock^T^ and HSF2oe^T^ cells to maintain cell-cell adhesion contacts was analyzed using Ultra-Low Attachment (ULA) round bottom plates, where a covalently bonded hydrogel surface of the wells stimulates the formation of spheroid-like structures. Transfected cells were treated with TGF-β alone or in combination with the selective TGF-β type I receptor inhibitor, SB431542, for 24 h in ULA plates. Mock^T^ cells formed compact spheroid-like structures under control conditions and the TGF-β treatment diminished this ability (Fig. 5F,G), which is indicative of destabilized cell-cell adhesions. Addition of SB431542 to the TGF-β-treated cells restored their capacity to form spheroid-like structures (Fig. 5F). Intriguingly, HSF2oe^T^ cells formed compact structures even in the presence of TGF-β (Fig. 5F), which denotes preserved cell-cell adhesions. Quantification of the spheroid areas corroborated our results (Fig. 5G), demonstrating that TGF-β-mediated downregulation of HSF2 leads to impaired cell-cell adhesion and forced expression of HSF2 overrides the TGF-β-induced destabilization of cell-cell contacts.

### HSF2 overrides the TGF-β-mediated inhibition of cell proliferation

Under physiological conditions and early phases of tumorigenesis, TGF-β is an important cell cycle regulator and inhibits cell growth by modulating the expression of cyclin-dependent kinases (CDKs), and CDK inhibitors (Principe et al., 2014; Zhang et al., 2017; Chen et al., 2020). Our RNA-seq analysis indicated that the major gene groups that were downregulated by TGF-β signaling but lost their responsiveness upon HSF2 overexpression are related to cell cycle progression (Fig. 2E,F). To gain more insight on how HSF2 affects the TGF-β-mediated regulation of the cell cycle, we expanded our analysis to include more genes important for DNA replication and cell proliferation, such as members of the MCM complex, the GINS gene family and cell cycle-associated kinases. Intriguingly, the expression of the examined genes was decreased by TGF-β, while overexpression of HSF2 dysregulated the TGF-β-dependent transcriptional changes (Fig. 6A). These results are in line with a recently published study reporting that HSF2 regulates the expression of cell cycle-associated genes across multiple cancer cell lines, including those derived from breast, prostate, lung, and colon cancer (Smith et al., 2022).

**Figure 6.**
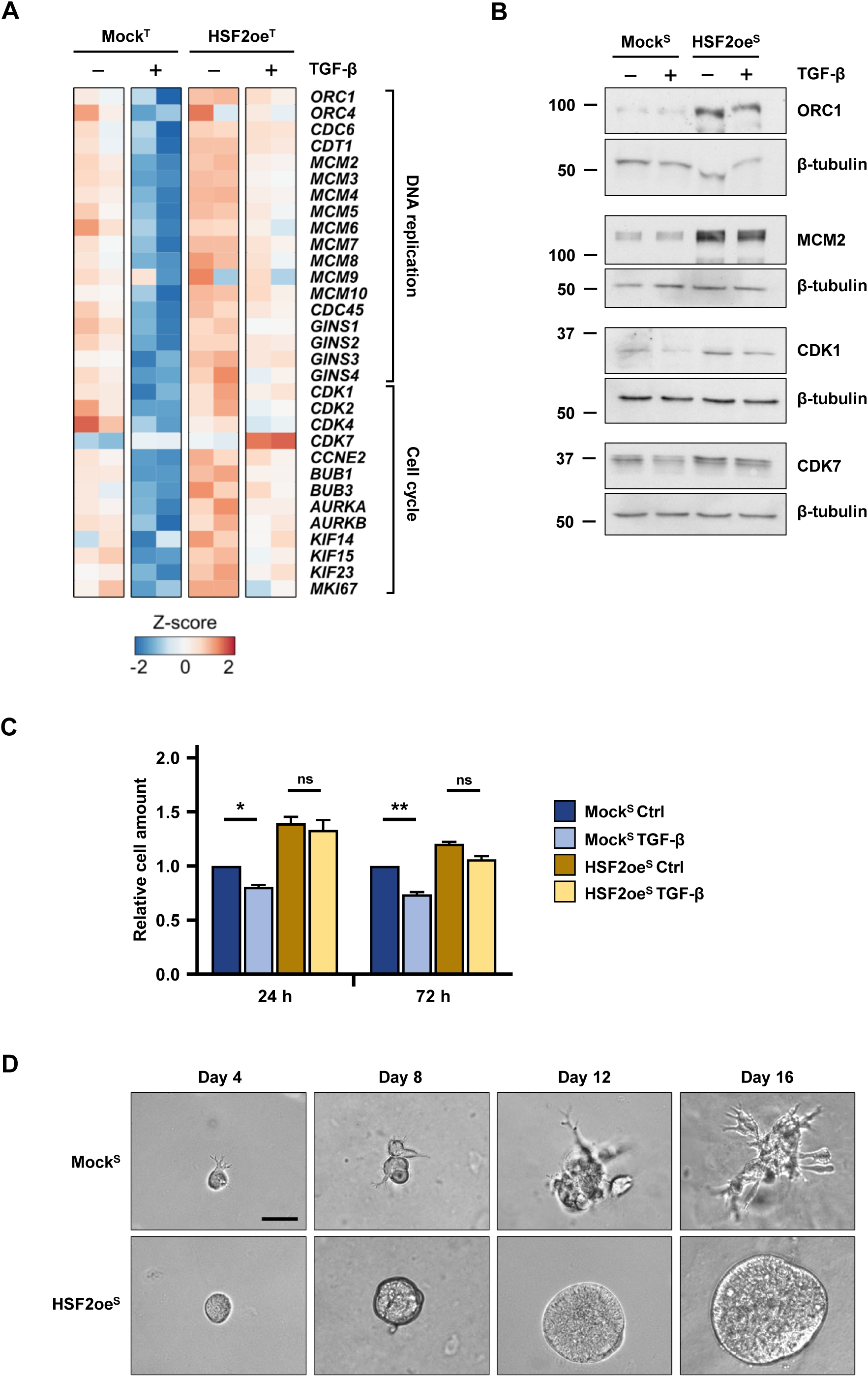
The presence of HSF2 counteracts the TGF-β-mediated inhibition of cell proliferation. **(A)** Heat map illustrating the expression changes of genes associated with DNA replication and cell cycle in HS578T cells transiently transfected with plasmids encoding GFP (Mock^T^) or HSF2 (HSF2oe^T^). Cells were treated with 10 ng/ml TGF-β_1_ or assay medium for 24 h. The Z-score was calculated based on the log₂CPM values for each gene. The double columns for each condition represent the two biological replicates from the RNA-seq. **(B)** Immunoblot analysis of ORC1, MCM2, CDK1, and CDK7 proteins in stable cell lines expressing GFP (Mock^S^) or HSF2 (HSF2oe^S^) treated with 10 ng/ml TGF-β_1_ or assay medium for 72 h. β-tubulin was used as a loading control. **(C)** CCK-8 Cell proliferation analysis of Mock^S^ and HSF2oe^S^ cells treated with 10 ng/ml TGF-β_1_ or assay medium (Ctrl) for 24 h or 72 h. The cell amount was calculated based on the absorbance and shown relative to the Mock^S^ Ctrl sample. Results are presented as mean ± SEM, * = p-value < 0.05, ** = p-value < 0.01, ns = not significant. The data represents three biological replicates. **(D)** Representative images of organotypic 3D cultures. Mock^S^ and HSF2oe^S^ cells were seeded on 7 mg/ml Matrigel at a density of 1 × 10^3^. Formation of organotypic tumoroids was followed and imaged at day 4, 8, 12, and 16 with Zeiss Axio Vert. A1 microscope using a 20x objective, NA 0.4. Scale bar 200 µm. The data represents three biological replicates.

We next analyzed the protein levels of selected targets in Mock^S^ and HSF2oe^S^ cells that were either treated with TGF-β or left untreated for 72 h. Origin recognition complex 1 (ORC1), a factor that promotes initiation of DNA replication by maintaining MCM helicases at the origin sequence (Duncker et al., 2009), increased substantially in cells expressing exogenous HSF2 in comparison to Mock^S^ cells (Fig. 6B; Fig. S6B). Similarly, minichromosome maintenance protein 2 (MCM2), a key component of the pre-replication complex involved in the initiation of eukaryotic genome replication (Forsburg, 2004), was elevated in HSF2oe^S^ cells (Fig. 6B; Fig. S6B). The expression of cyclin-dependent kinases 1 and 7 (CDK1, CDK7), both well-known regulators of cell cycle progression (Fisher, 2005; Enserink et al., 2010), were decreased in TGF-β-treated Mock^S^ cells and their expression remained unaltered in HSF2oe^S^ cells (Fig. 6B; Fig. S6B).

TGF-β signaling affects cell proliferation in normal and tumor cells, and the mode of action can vary depending on the phase in which the cell is in its life cycle or stage of tumorigenesis (Sengupta et al., 2014). To examine the effect of HSF2 on cell proliferation, Mock^S^ and HSF2oe^S^ cells were subjected to TGF-β for either 24 h or 72 h. Using the Cell Counting Kit-8 (CCK-8) assay, we quantified the differences in cell amounts that are indicative of cell proliferation. In line with previous findings demonstrating that Snail inhibits cell cycle progression (Vega et al., 2004), the TGF-β treatment significantly decreased cell proliferation of Mock^S^ cells when compared to untreated cells (Fig. 6C). Interestingly, cell proliferation was markedly higher in HSF2oe^S^ cells, and there was no significant change between the untreated and TGF-β-treated cells (Fig. 6C). Based on these results, we propose that the loss of HSF2 is a requirement for the TGF-β-mediated inhibition of cell proliferation.

To further assess the proliferation and invasion capacity of Mock^S^ and HSF2oe^S^ cells, we analyzed 3D Matrigel cultures where the formation of organotypic tumoroids was followed up to 16 days. Mock^S^ cells started developing invasive structures already at day 4, which gradually increased towards the end point of the assay (Fig. 6D). Remarkably, HSF2oe^S^ cells failed to form invasive structures despite the rapid growth (Fig. 6D), thereby supporting our data on induced proliferation of HSF2oe^S^ cells (Fig. 6C). Upon activation of invasive behavior, malignant cells reduce their proliferation to enable enhanced migration (De Donatis et al., 2010). Thus, our results showed that Mock^S^ cells stopped proliferating to induce cell migration, whereas the rapidly proliferating HSF2oe^S^ cells were inhibited in their ability of becoming invasive.

### HSF2 promotes proliferation and secondary tumor formation in *in vivo* zebrafish xenografts

After performing several cell-based *in vitro* assays, we wanted to know whether the obtained results were applicable also *in vivo*. For this purpose, we examined the capacity of Mock^S^ and HSF2oe^S^ cells to promote tumor growth and invasiveness using the zebrafish xenograft model system (Pekkonen et al., 2018; Asghar et al., 2021). Cells were nanoinjected into the pericardial cavity of 2-day-old zebrafish embryos and the xenografts were imaged at one day post-injection (1dpi) and four days post-injection (4dpi) (Fig. 7A). In comparison to Mock^S^, HSF2-overexpressing cells displayed significantly larger tumor area and impaired invasion capacity (Fig. 7B), reinforcing the anti-invasive effect of HSF2 in breast cancer cells. To more closely examine the proliferation rate of Mock^S^ and HSF2oe^S^ cells, we analyzed the expression of proliferation marker Ki67 in the formed xenograft tumors by immunofluorescence. The cells were again nanoinjected into the pericardial cavity of zebrafish embryos and after 24 h, the animals were fixed and stained with Ki67 (Fig. 7C). Immunofluorescence analysis revealed that the HSF2oe^S^ xenografts displayed clearly more abundant Ki67 expression than the Mock^S^ xenografts (Fig. 7C). Furthermore, analysis of the relative Ki67 area within the tumors demonstrated that a significantly larger proportion of the HSF2oe^S^ cells were Ki67-positive (Fig. 7D), corroborating that HSF2 drives cell proliferation and tumor expansion *in vivo*.

**Figure 7.**
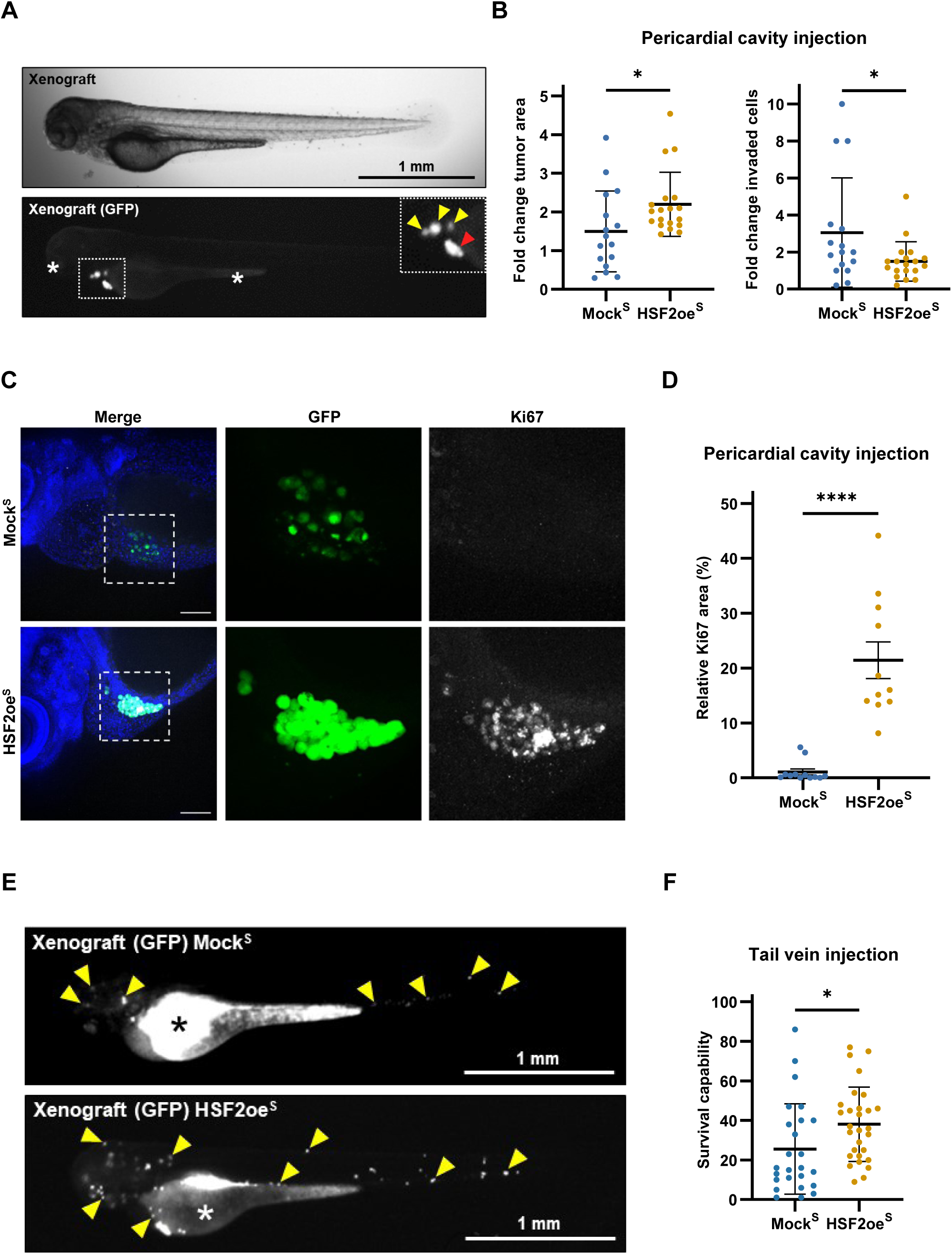
Ectopic expression of HSF2 induces cell proliferation and secondary tumor formation in an *in vivo* zebrafish xenograft model. **(A)** Representative wide-field and GFP images of a HSF2oe^S^ zebrafish xenograft. Mock^S^ or HSF2oe^S^ cells were nanoinjected into the pericardial cavity of 2-day-old zebrafish embryos. Tumor growth and cell invasiveness were followed at one day post-injection (1dpi) and four days post-injection (4dpi). Dotted line indicates area of inset. Red arrowhead in the inset indicates primary tumor and yellow arrowheads indicate disseminated cancer cells. Asterisk (*) denotes non-specific fluorescence. Images were acquired using Nikon Eclipse Ti2 microscope equipped with a Plan UW 2x objective. Scale bar 1 mm. **(B)** Fold change of the tumor area and invaded cells in Mock^S^ and HSF2oe^S^ zebrafish xenografts 1dpi vs 4dpi. Results are presented as mean ± SD, * = p-value < 0.05. For Mock^S^ 16 and for HSF2oe^S^ 19 zebrafish xenografts were analyzed. **(C)** Representative immunofluorescence images of Mock^S^ and HSF2oe^S^ zebrafish xenografts. Cells were stained with CellTracker Green CMFDA and nanoinjected into the pericardial cavity of 2-day-old zebrafish embryos. After 24 h, the animals were fixed and stained with an antibody against the proliferation marker Ki67. DAPI was used as a nuclear marker. Dotted lines indicate areas of insets. Images were acquired using 3i CSU-W1 Spinning Disk confocal microscope with a 20x objective, NA 0.8. Scale bar 100 µm. **(D)** Relative Ki67 area in Mock^S^ and HSF2oe^S^ tumors 24 h after pericardial cavity injection. Results are presented as mean ± SEM, **** = p-value < 0.0001. For Mock^S^ 12 and for HSF2oe^S^ 11 zebrafish xenografts were analyzed. **(E)** Representative GFP images of Mock^S^ and HSF2oe^S^ zebrafish xenografts. Mock^S^ or HSF2oe^S^ cells were stained with CellTracker Green CMFDA and nanoinjected into the tail vein of 2-day-old zebrafish embryos. The survival capability of cells was investigated by following secondary tumor formation 24 h post-injection. Yellow arrowheads indicate examples of secondary tumors. Asterisk (*) denotes non-specific fluorescence. Images were acquired using Nikon Eclipse Ti2 microscope equipped with a Plan UW 2x objective. Scale bar 1 mm. **(F)** The survival capability of Mock^S^ and HSF2oe^S^ cells 24 h after tail vein injection. Results are presented as mean ± SD, * = p-value < 0.05. For Mock^S^ 25 and for HSF2oe^S^ 29 zebrafish xenografts were analyzed.

Metastasizing cancer cells must withstand an extensive amount of environmental stress to be able to form secondary tumors. The ability of malignant cells to resist and adapt to these stressors is a critical determinant underlying their survival capacity and metastatic potential (Fares et al., 2020). Therefore, we investigated the survival capacity of Mock^S^ and HSF2oe^S^ cells by nanoninjecting them directly into the tail vein of zebrafish embryos and by following secondary tumor formation. Direct tail vein injection bypasses the step where cancer cells escape the primary tumor and mimics the state when metastatic cancer cells circulate in the blood flow prone for secondary metastasis. Strikingly, we found that HSF2oe^S^ cells formed significantly more secondary tumors than Mock^S^ cells (Fig. 7E,F), which provides strong evidence for enhanced stress resistance of HSF2-overexpressing cells. Altogether, our results from all cell-based *in vitro* and *in vivo* assays demonstrate that while downregulation of HSF2 is instrumental for the activation of TGF-β-induced EMT and invasiveness, elevated levels of HSF2 are critical for supporting malignant cell proliferation and stress tolerance.

### HSF2 expression and activity increase during breast cancer progression

So far, we have demonstrated that TGF-β-mediated downregulation of HSF2 characterizes the phenotypic switch that occurs during EMT, and that abundant HSF2 expression correlates with induced cell proliferation and stress resistance. To gain knowledge how the acquired results align with human patient material, we analyzed the expression and subcellular localization pattern of HSF2 in healthy breast tissue, pre-invasive DCIS, and IDC (Fig. 8A). HSF1 was included in the analysis as it is strongly linked with the malignant phenotype by promoting cancer-specific transcriptional programs, invasion and metastasis, cell proliferation and survival as well as metabolic reprogramming (Zhao et al., 2009; Santagata et al., 2011; Mendillo et al., 2012; Xi et al., 2012; Santagata et al., 2013; Scherz-Shouval et al., 2014; Carpenter et al., 2017; Jacobs et al., 2024). Fixed frozen tissue sections were stained with antibodies against HSF2 and HSF1. Keratin 8 (KRT8) was used to identify cells of epithelial origin and DAPI as a nuclear marker. Recently, we reported that in formalin-fixed paraffin-embedded benign human breast tissues HSF2 is expressed predominantly in the cytoplasm, whereas HSF1 was detected both in the nucleus and the cytoplasm (Joutsen et al., 2024). Here, we found that both HSF2 and HSF1 localize in the cytoplasm of KRT8-positive breast epithelial cells in healthy breast tissue samples (Fig. 8B,C). In DCIS and IDC, the levels of HSF2 were significantly elevated in comparison to healthy tissue (Fig. 8B,D,E). Curiously, HSF2 displayed a clearly distinct expression pattern, forming hotspots of nuclear HSF2 in DCIS tumors (Fig. 8B,D). This data demonstrates that the subcellular localization of HSF2 converts from cytoplasmic in the healthy cells to nuclear in the pre-invasive cells. The results from several cell-based systems showed that overexpression of HSF2 upregulates cell cycle genes and promotes rapid spheroid and tumor growth in 3D cultures and in zebrafish xenografts, respectively (Figs. 2,6,7). Together with the observed expression pattern of HSF2 in the DCIS samples, we hypothesized that increased nuclear HSF2 is required for the pre-invasive tumor expansion. Our zebrafish tail vein injection data highlighted an eminent survival capability of HSF2oe^S^ cells (Fig. 7C). As the nuclear expression of HSF2 remained elevated in the IDC tumors (Fig. 8B,D), we predicted that high HSF2 in IDC is required to promote cancer cell survival and stress resistance. The expression and nuclear localization of HSF1 were also induced in DCIS and IDC samples when comparing to healthy tissue (Fig. 8C,G,H). In contrast to HSF2, which formed distinct nuclear hotspots in DCIS, HSF1 was homogenously expressed in all studied tissues (Fig. 8C,G). Moreover, while the nuclear localization of HSF2 was dramatically increased already in the pre-invasive state (Fig. 8F), the most abundant nuclear expression of HSF1 was observed in invasive breast cancer (Fig. 8I).

**Figure 8.**
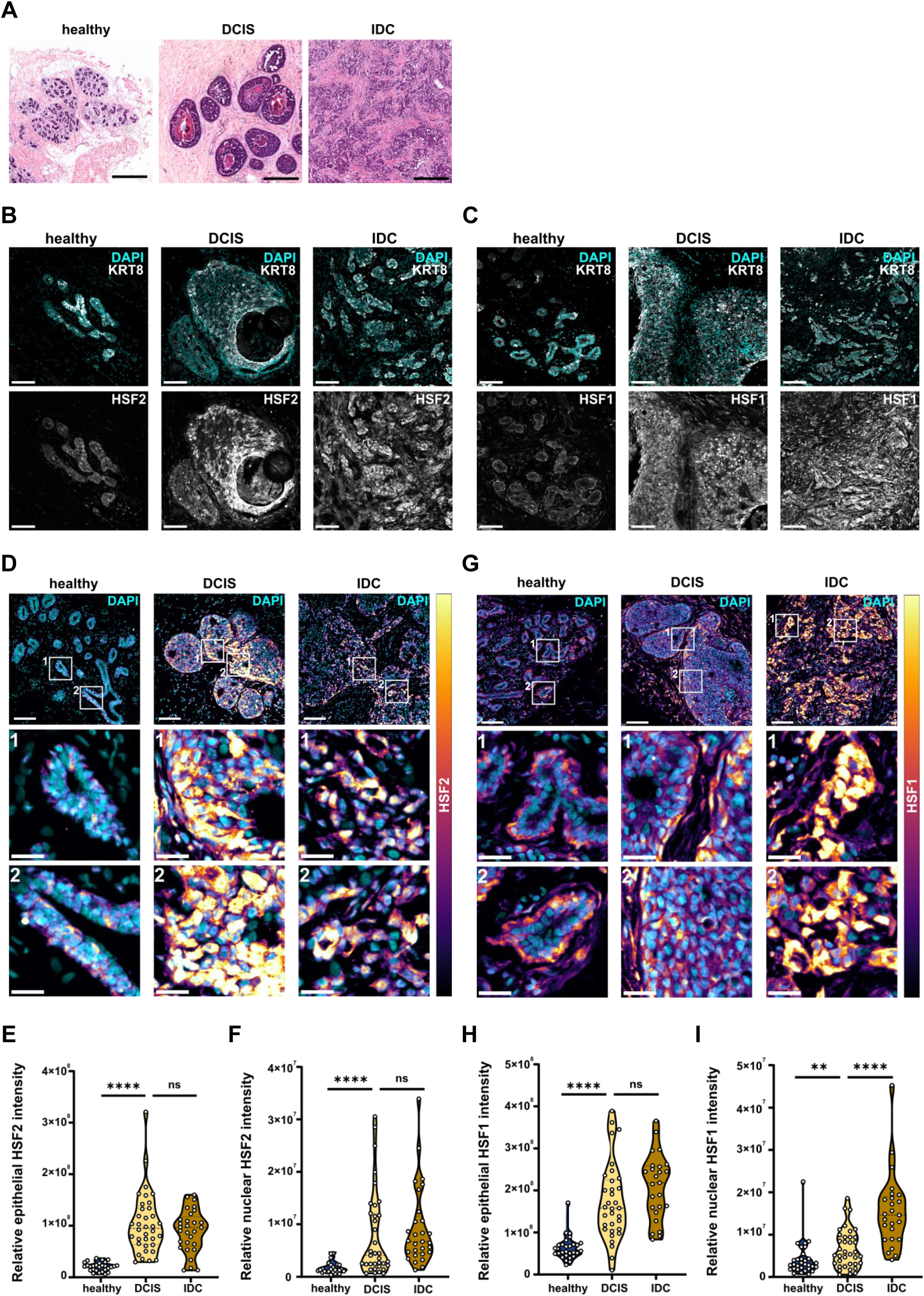
HSF2 expression and activity increase during breast cancer progression. **(A)** Representative hematoxylin-eosin staining of tissue sections from healthy donors, patients with ductal carcinoma *in situ* (DCIS) and patients with invasive ductal carcinoma (IDC). Scale bars 200 µm. **(B)** Representative immunofluorescence images from indicated patient tissue sections stained with keratin 8 (KRT8) and DAPI (top row) and HSF2 (bottom row). n = 6-10 FOVs from 3-4 patients. Scale bars 100 µm. **(C)** Representative immunofluorescence images from indicated patient tissue sections stained with KRT8 and DAPI (top row) and HSF1 (bottom row). n = 6-10 FOVs from 3-4 patients. Scale bars 100 µm. **(D)** Representative immunofluorescence images from indicated patient tissue sections stained with HSF2 and DAPI. HSF2 expression is visualized with an intensity-coded look-up table (high values – yellow). n = 6-10 fields of view (FOVs) from 3-4 patients. Scale bars 100 µm (main) and 25 µm (regions of interest, ROIs) **(E)** Relative epithelial intensity of HSF2 in the patient tissues as determined by keratin 8 (KRT8). Results are presented as violin plots, **** = p-value < 0.0001, ** = p-value < 0.01. n = 6-10 FOVs from 3-4 patients. **(F)** Relative nuclear intensity of HSF2 in the patient tissues as determined by DAPI signal within the epithelial area. Results are presented as violin plots, **** = p-value < 0.0001. n = 6-10 FOVs from 3-4 patients. **(G)** Representative immunofluorescence images from patient tissue sections stained with HSF1 and DAPI. HSF1 expression is visualized with an intensity-coded look-up table (high values – yellow). n = 6-10 FOVs from 3-4 patients. Scale bars 100 µm (main) and 25 µm (ROIs). **(H)** Relative epithelial intensity of HSF1 in the patient tissues as determined by KRT8. Results are presented as violin plots, **** = p-value 0.0001. n = 6-10 FOVs from 3-4 patients. **(I)** Relative nuclear intensity of HSF1 in the patient tissues as determined by DAPI signal within the epithelial area. Results are presented as violin plots, **** = p-value < 0.0001, ** = p-value < 0.01. n = 6-10 FOVs from 3-4 patients.

Finally, we tested our hypothesis that nuclear HSF2 contributes to a hyperplastic cell phenotype in DCIS by analyzing its co-localization pattern with the proliferation marker Ki67 (Fig. 9A). Strikingly, our results revealed that co-expression of HSF2 and Ki67 in the nucleus increased significantly in DCIS and remained elevated in IDC (Fig. 9A,B), suggesting that HSF2 translocates to the nucleus specifically in actively proliferating cells. Notably, no alterations were observed in the co-localization pattern of nuclear HSF1 and Ki67 across the examined tissue samples (Fig. 9C,D). These findings corroborate our hypothesis by demonstrating that HSF2 designates the pre-invasive cells with high proliferative capacity. Collectively, our results from cell-based *in vitro* and *in vivo* assays together with human patient samples, we conclude that unlike HSF1, which supports malignancy in advanced breast cancer, HSF2 acts as a switch between the proliferative and invasive phenotypes at the initial steps of breast cancer progression.

**Figure 9.**
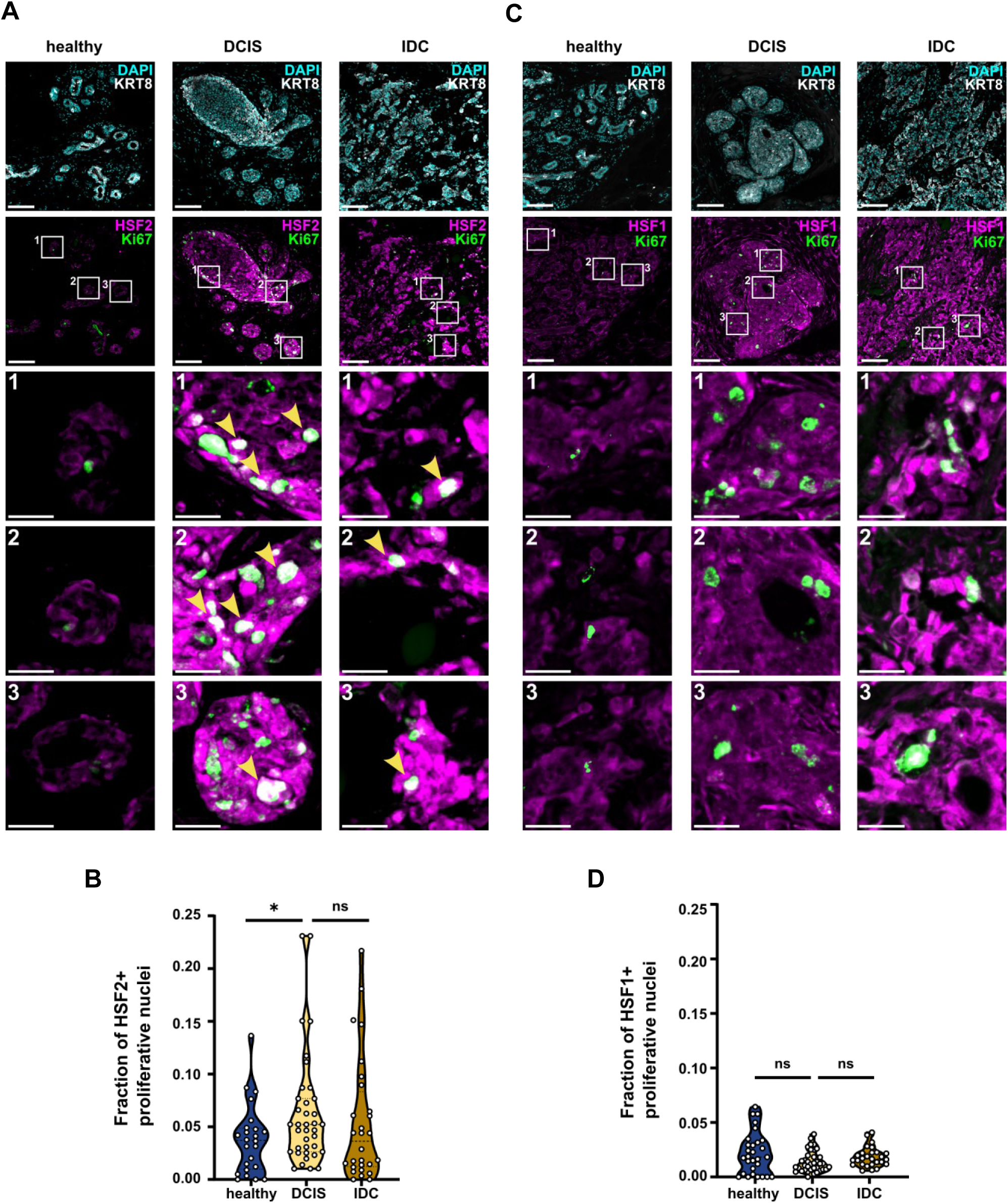
Nuclear co-localization of HSF2 and Ki67 increases in pre-invasive and advanced breast cancer. **(A)** Representative immunofluorescence images from patient tissue sections stained with keratin 8 (KRT8) and DAPI (top row) or HSF2 and Ki67 (middle and bottom rows). Nuclei positive for both Ki67 and HSF2 are visualized as white (indicated by yellow arrowheads). n = 6-10 FOVs from 3-4 patients. Scale bars 100 µm (main) and 25 µm (ROIs). **(B)** Fraction of HSF2-positive (HSF2+) proliferative cells within the epithelium. Results are presented as violin plots, * = p-value < 0.05. n = 6-10 FOVs from 3-4 patients. **(C)** Representative immunofluorescence images of patient tissue sections stained with KRT8 and DAPI (top row) or HSF1 and Ki67 (middle and bottom rows). n = 6-10 FOVs from 3-4 patients. Scale bars 100 µm (main) and 25 µm (ROIs). **(D)** Fraction of HSF1-positive (HSF1+) proliferative cells within the epithelium. Results are presented as violin plots. n = 6-10 FOVs from 3-4 patients.

## DISCUSSION

Despite accumulating evidence linking HSF2 to malignant transformation (Mustafa et al., 2010; Zhong et al., 2016; Björk et al., 2016; Meng et al., 2017; Yang et al., 2019; Smith et al., 2022), its functional role during cancer progression and the signaling pathways regulating HSF2 during tumorigenesis have remained elusive. Here, we show that TGF-β stimulation, mimicking the early steps of EMT-mediated invasion, dramatically downregulates HSF2 expression to enable the activation of TGF-β-induced pro-metastatic gene programs. Remarkably, ectopically expressed HSF2 in human breast cancer cells counteracted the TGF-β-mediated effects on gene expression and cellular properties, including cell proliferation and invasion, in both *in vitro* and *in vivo* assays. We further demonstrate that in patient tissue samples, HSF2 is dynamically expressed during breast cancer progression. Based on our results, we present a model where high levels and nuclear accumulation of HSF2 drive cell proliferation in the pre-invasive DCIS, whereas a temporal downregulation of HSF2 through pro-tumorigenic TGF-β signaling activates EMT, allowing cancer cell invasion (Fig. 10). In the invasive stage, IDC, HSF2 expression is elevated and contributes to pronounced stress tolerance and proliferation of cancer cells (Fig. 10). Collectively, we propose that dynamic regulation of HSF2 steers the balance between cell proliferation and invasion, which is a critical determinant of the invasive transition in cancer progression.

**Figure 10.**
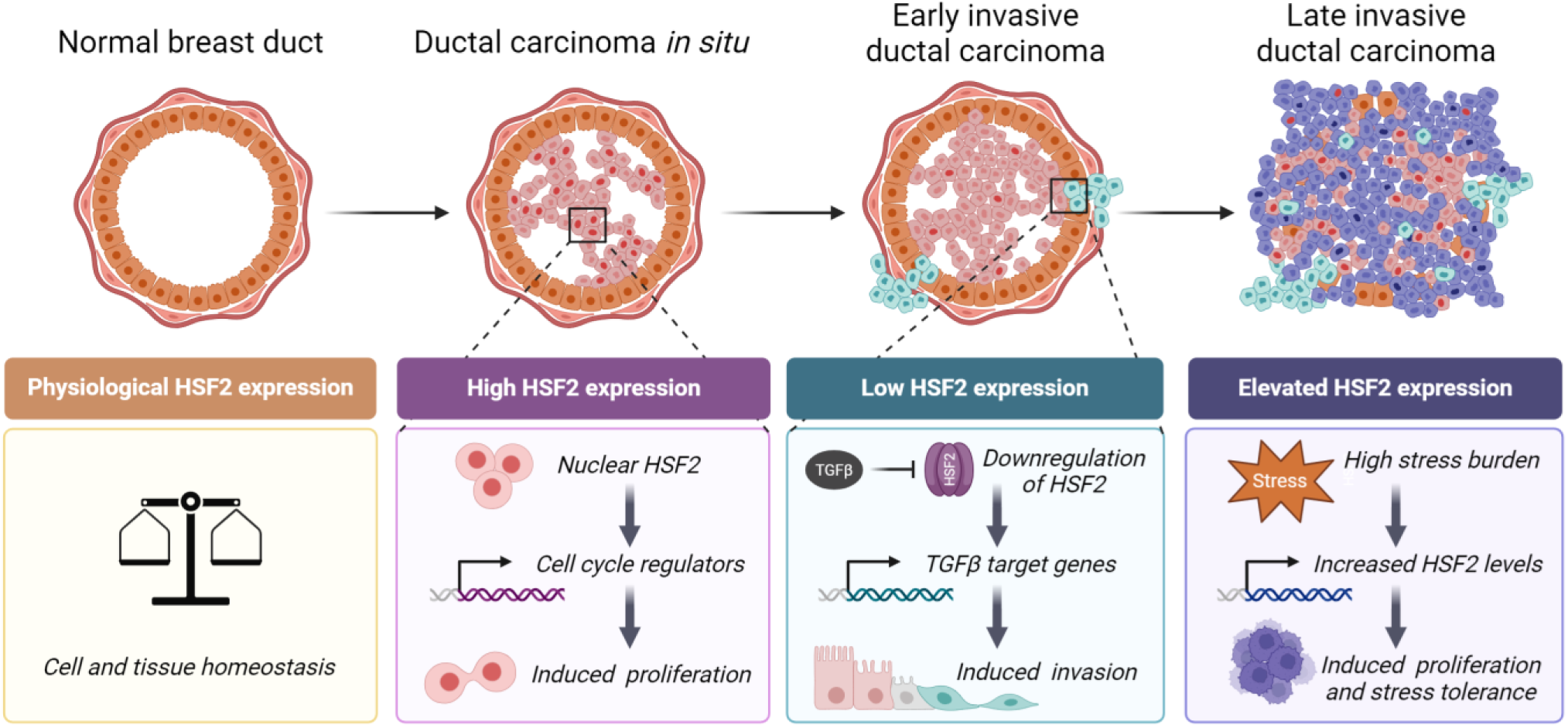
**Dynamic regulation of HSF2 drives breast cancer progression** A schematic model on the dynamic expression of HSF2 during different stages of breast cancer progression. In normal breast epithelium, physiological HSF2 expression contributes to the maintenance of cell and tissue homeostasis. In the pre-invasive stage of breast cancer, ductal carcinoma *in situ*, increased expression, and nuclear localization of HSF2 activate the transcription of genes regulating the cell cycle, leading to induced proliferation of ductal epithelial cells. At the early stage of invasive ductal carcinoma, pro-tumorigenic TGF-β signaling downregulates HSF2 to enable expression of TGF-β target genes and activation of the EMT program in a subset of transformed cells (turquoise). Subsequently, cells undergoing EMT acquire induced migratory abilities and invade through the basement membrane, forming microinvasions around the breast ducts. At the late stage of invasive ductal carcinoma, high stress burden in the tumor microenvironment leads to elevated levels of HSF2 expression and nuclear localization to promote proliferation, stress tolerance and cancer cell survival.

The function of HSF1 is well established across diverse cancers, and it is recognized as a key driver of tumor-supportive transcriptional programs in both cancer cells and the adjacent cancer-associated fibroblasts (CAFs) (Mendillo et al., 2012; Scherz-Shouval et al., 2014; Levi-Galibov et al., 2020; Shaashua et al., 2022). Moreover, due to its prominent disease-supporting role, several inhibitors targeting HSF1 have been developed (Dong et al., 2020; Dong et al., 2022; Pasqua et al., 2023). Interestingly, HSF2 was recently reported to act as a critical supporter of tumor progression by functionally cooperating with HSF1 in several cancer cell lines, including breast cancer cells (Smith et al., 2022). In this study, however, we could not observe any changes in HSF1 expression upon TGF-β treatment, and HSF1 was not directly involved in the TGF-β-mediated regulation of HSF2 levels. The robust downregulation of HSF2 was evident already within 24 h of TGF-β stimulation, mimicking the primary steps of the invasive transition. These findings strongly support our hypothesis that the divergence of HSF1 and HSF2 functions is explicitly important at the initial phases of malignant transformation. TGF-β is one of the main cytokines mediating the interplay between tumor cells and CAFs (Linares et al., 2021), and it is secreted by CAFs in an HSF1-dependent manner (Scherz-Shouval et al., 2014). In colon cancer, HSF1 is upregulated in CAFs where it regulates the expression and secretion of TGF-β and stromal cell-derived factor 1, another important cytokine supporting tumor growth (Scherz-Shouval et al., 2014). In addition, HSF1 contributes to stromal heterogeneity by modulating the ratio of immune-regulatory clusterin-positive CAFs in pancreatic ductal adenocarcinoma with germline mutations in the BRCA gene (Shaashua et al., 2022). Considering the previous reports and the results obtained in this study, it is tempting to speculate that the intercellular signaling between CAFs and cancer cells occurs through crosstalk mediated by HSFs. Accordingly, HSF1 in CAFs stimulates TGF-β secretion allowing adjacent cancer cells to initiate pro-metastatic cell processes due to TGF-β-induced downregulation of HSF2. This would describe an unprecedented non-cell autonomous regulation of HSFs, exemplifying that their impact on cancer progression is more multifaceted than previously anticipated. Thereby, future studies should focus on the mechanisms by which CAF-secreted TGF-β affects HSF2 in malignant cells and elucidate the possible crosstalk of HSF1 and HSF2 in tumorigenesis.

The initiation and early growth of epithelial carcinomas is characterized by the activation of EMT. During EMT, cells lose their epithelial-specific cell-cell junctions and acquire a more motile mesenchymal phenotype with the ability to modify the local microenvironment (Dongre & Weinberg, 2019). TGF-β is one of the key cytokines promoting EMT, and it acts through a selection of transcription factors, which repress the expression of E-cadherin, control the reorganization of cytoskeletal proteins, and enable the production of ECM components (Dongre & Weinberg, 2019; Lee & Massagué, 2022). Here, we observed that HSF2 overrides this regulatory pathway as HSF2-overexpressing cells displayed substantially elevated protein levels of E-cadherin and were able to stabilize cell-cell adhesions even in the presence of TGF-β. In addition, forced expression of HSF2 blocked the TGF-β-mediated upregulation of several key invasion-associated matrix proteins and diminished the ability of *in vitro* vasculogenic mimicry. Interestingly, in a recent report from the Hannon laboratory, vasculogenic mimicry was shown to be promoted by the transcription factor FOXC2 in several solid tumors (Cannell et al., 2023), while we identified HSF2 as a suppressor of vasculogenic mimicry in breast cancer cells. Furthermore, HSF2 overexpression altered the subcellular organization of the vimentin cytoskeleton and decreased cellular stiffness. TGF-β-induced increase in cell stiffness has been demonstrated as a requirement for enhanced invasion in non-small cell lung carcinoma (Gladilin et al., 2019). Indeed, by investigating cell migration in both *in vitro* 3D cell cultures and *in vivo* zebrafish xenografts, we showed that the presence of HSF2 markedly impairs cellular invasiveness. These results indicate that HSF2 prevents epithelial cells from acquiring a mesenchymal phenotype. We have previously reported that the lack of HSF2 leads to downregulation of cadherin-mediated cell-cell adhesions in osteosarcoma cells (Joutsen et al., 2020). More recently, the importance of HSF2 in neuroepithelial integrity was reported using organoids modeling the Rubinstein-Taybi neurodevelopmental disorder, where destabilization of HSF2 caused dysregulation of N-cadherin in neuronal cells (de Thonel et al., 2022). Based on these findings, HSF2 contributes to the maintenance of cell identity. Cells that undergo EMT-linked phenotypic switching are typically considered as cells eventually entering the invasion-metastasis cascade. Once metastasized, cells have the possibility to revert to a more epithelial phenotype by using the reverse process of EMT, *i.e.*, mesenchymal-epithelial transition (Yao et al., 2011). Our data from tail vein-injected zebrafish embryos, where the intravasation step is bypassed, showed that HSF2-expressing cells have a pronounced survival capability and an enhanced ability to form secondary tumors. Hence, it is plausible that HSF2 expression is restored in the established secondary tumor to promote growth and stress tolerance. This work unravels that HSF2 controls the initial phenotypic switch of carcinoma cells, and we propose the TGF-β-mediated loss of HSF2 as a critical mechanism in cancer progression to a metastatic stage.

The dynamic nature of HSF2 has been described in multiple contexts, including cell cycle, spermatogenesis, and embryonal development (Pessa et al., 2024). A striking example is mouse embryogenesis, where HSF2 levels fluctuate dramatically (Mezger et al., 1994; Rallu et al., 1997; Abane & Mezger, 2010). Our findings in cell-based models and in human patient tissue samples portray an equally dynamic expression and activation pattern of HSF2 during breast cancer progression (Fig. 10). Our recent comprehensive analysis of formalin-fixed paraffin-embedded benign human tissues revealed that HSF2 localizes predominantly in the cytoplasm across examined samples, including breast epithelium (Joutsen et al., 2024). Accordingly, HSF2 was found in the cytoplasm of epithelial cells in healthy breast tissue. We demonstrate that HSF2 was significantly upregulated, and its nuclear localization increased in hyperplastic DCIS. Based on our data from transcriptional, protein, and functional analyses, we connect the consequences of HSF2 activation with induced cell proliferation. Ki67 is commonly used as a prognostic marker for invasive breast cancer and its high levels are associated with an elevated risk of DCIS recurrence (Davis et al., 2016; Poulakaki et al., 2018; Wang et al., 2024). However, the drivers of the invasive transition from DCIS to IDC are still poorly characterized and the current biomarkers are unable to reliably distinguish the high-risk lesions (Wang et al., 2024). To this end, this study provides strong advancement in the diagnostics of breast cancer progression. Since the nuclear intensity of HSF2 and its co-localization with Ki67 were strikingly increased in a subpopulation of proliferative DCIS cells, we identify HSF2 as a novel and potent biomarker for high-risk DCIS. Further characterization of the HSF2-high DCIS hotspots, using advanced spatial transcriptional and proteomics approaches in combination with a systematic DCIS patient analysis, is required to assess the prognostic and therapeutic potential of HSF2.

In this work, we found that TGF-β causes a robust downregulation of HSF2. The decrease in HSF2 expression is crucial for TGF-β to exert its impact on cell-cell adhesion, proliferation, cellular elasticity as well as cancer cell migration and invasion. In contrast, when the downregulation of HSF2 was bypassed through its ectopic expression, the activation of TGF-β-induced gene programs and their downstream effects on cellular properties were severely disrupted. Our experimental approach, comprising a wide repertoire of *in vitro* and *in vivo* models as well as analysis of human tissue samples, uncovers a biologically relevant and novel role for HSF2 in breast cancer. Altogether, we propose that dynamic regulation of HSF2 steers the balance between cell proliferation and invasion during breast cancer progression.

## MATERIALS AND METHODS

### Cell culture, treatments, and transient transfections

Cells were grown at 37°C in a humidified 5% CO_2_ atmosphere. Human transformed breast epithelial MDA-MB-231 cells, human embryonic kidney HEK-293 T cells and human dermal fibroblasts (HDFs) were cultured in Dulbecco’s Modified Eagle’s medium (DMEM, Sigma-Aldrich), supplemented with 10% fetal bovine serum (FBS, Serena), 2 mM L-glutamine (Biowest) and 100 µg/ml penicillin-streptomycin (Biowest). Human HS578T transformed breast epithelial cells were cultured in DMEM supplemented with 10% FBS, 2 mM L-glutamine, 100 µg/ml penicillin-streptomycin, and 10 µg/ml insulin (A11382II, Gibco). Human MCF10A breast epithelial cells were cultured in DMEM/F12 (DMEM/Nutrient Mixture F-12, Gibco) supplemented with 5% fetal horse serum (Gibco), 10 µg/ml insulin, 0.5 ng/ml hydrocortisone (Sigma-Aldrich), 20 ng/ml EGF, 100 ng/ml cholera toxin (Sigma-Aldrich), and 100 µg/ml penicillin-streptomycin. All used cells were immortalized cell lines purchased from ATCC and provided with certificates of negative mycoplasma status.

For treatments, the medium was supplemented with 2% serum (referred to as assay medium). Cells were treated with assay medium supplemented with 10 ng/ml transforming growth factor-beta 1 (hereafter referred to as TGF-β, 240-B-002, R&D Systems) for 24 h or 72 h (Peinado et al., 2003; Medici et al., 2006). For the longer treatment, medium was replaced after 48 h. To induce canonical and non-canonical TGF-β signaling, cells were treated with assay medium supplemented with StemXVivo EMT Inducing Media Supplement (EMT supplement, CCM017, R&D Systems). To inhibit TGF-β signaling, cells were treated with 10 µM SB431542 (S4317, Sigma-Aldrich), a TGF-β receptor type 1 inhibitor (Inman et al., 2002).

All transient transfections were performed with the Neon Transfection System (MPK5000, InVitrogen) according to the manufacturer’s instructions. In brief, 1.8 × 10^6^ HS578T or 2.2 × 10^6^ MDA-MB-231 cells were suspended in 100 µl Buffer R containing either 25 µg plasmid and/or 3 µM siRNA. Next, HS578T and MDA-MB-231 cells were subjected to electroporation (1050 V, 20 ms, 3 pulses and 1350 V, 10 ms, 4 pulses, respectively) in the transfection tube containing 3 ml Buffer E2. Cells used for the luciferase assay were transfected with 20 µg of the plasmids encoding luciferase reporter genes and 5 µg of the plasmids encoding β-galactosidase. Transfected cells were allowed to recover for 24 h prior to the treatments.

### Generation of stable cell lines

HEK-293 T cells were cultured in a T25 flask to a confluency of 70-80%. Next day, cells were transfected using Lipofectamine 3000 (Thermo Fisher Scientific) in accordance with the manufacturer’s instructions. Briefly, 14 µl of Lipofectamine reagent was mixed with 500 µl Opti-MEM in a 1.5 ml microcentrifuge tube, and in a separate tube, viral vectors pLP1 (1.7 µg), pLP2 (1.1 µg), PLP/VSVg (1.7 µg), and the expression vector (1.5 µg) were mixed in 500 µl Opti-MEM containing 12 µl p3000 reagent. Expression vectors containing either GFP alone (Mock^S^) or HSF2 and GFP (HSF2oe^S^, connected with a T2A sequence) were designed and purchased from VectorBuilder (Inc). Each microcentrifuge tube was carefully mixed by pipetting and incubated for 15 min at RT. Meanwhile, 4 ml of fresh DMEM was added to each T25 flask and following the incubation period the transfection mix was added dropwise to the cells. Thereafter, cells were incubated at 37°C with 5% CO_2_ overnight. Next day the medium was replaced in each flask. The supernatant containing virus was collected 72 h post-transfection and centrifuged 10 min at 1000 × g. The pool of viral medium was filtered through a 0.45 μm pore size filter, supplemented with 8 µg/ml of polybrene and applied on HS578T cells in a 6-well plate. Next, spinoculation was performed at 450 × g for 30 min at RT. The medium was changed 48 h post-transduction, and antibiotic selection with puromycin was started 72 h post-transduction. Following two weeks of culturing, the negative lentiviral status was confirmed for each cell line using the HIV-1 p24 DuoSet ELISA rom kit (DY7360-05, R&D Systems).

### Plasmid construction

The plasmids used for the luciferase assay were generated by cloning 1 kb of either the human *HSF2* promoter or *MMP9* promoter upstream of the respective TSS into a STARR-seq luciferase validation vector_ORI_empty vector (#99297, Addgene). The DNA from HDFs was isolated by first lysing cells in 3 ml lysis buffer (10 mM Tris-HCl, 400 mM NaCl, 2 mM Na_2_ EDTA [pH 8.2]). Next, 200 µl 10% SDS and 500 µl Proteinase K buffer (1 mg Proteinase K, 1% SDS, 2 mM Na_2_ EDTA [pH 8.2]) was added to the lysis buffer, and the mixture was incubated at 37°C for 24 h. Following this, 1 ml of 5 M NaCl was mixed into the solution containing lysis and Proteinase K buffer. The solution was vortexed and centrifuged at 10,000 × g to remove cell debris. The supernatant containing DNA was mixed with 10 ml absolute ethanol, and the DNA precipitate was isolated by centrifugation. The DNA pellet was washed with absolute ethanol, and residual ethanol was removed from the DNA sample by evaporation at 50°C. The purified DNA pellet was diluted in water. For the luciferase reporter constructs, *SBEluc* and *HSF2luc*, 1 kb promoter sequences (upstream of the TSS) of *MMP9* or *HSF2* were inserted into the linear vector upstream of the luciferase reporter gene using the In-Fusion HD cloning kit (Takara Bio) according to the manufacturer’s instructions. For the *SBEluc* construct, a gene strand containing four SMAD-binding elements (SBEs), purchased from Eurofins Genomics, was inserted in the N-terminus of the *MMP9* promoter to induce SMAD-binding and thereby promoter responsiveness to TGF-β (Itoh et al., 2019), with the In-Fusion HD cloning kit (Takara Bio).

The plasmid encoding exogenous HSF2 was generated by inserting a fragment containing the protein coding sequence of HSF2 and a myc-tag. In addition, a Strep-Tag II, purchased from Eurofins Genomics, was placed downstream of the myc-tag. PCR was used to amplify the whole fragment containing a CMV promoter, HSF2-myc-StrepII, and an SV40 (polyA) tail, which was inserted into a pEGFP-N2 vector (GenBank Accession #U57608). The final construct (HSF2oe) enabled exogenous expression of both HSF2-myc-Strep II and GFP via separate CMV promoters. The pEGFP-N2 plasmid was used as Mock. All primer sequences used for plasmid constructions are listed in Supplementary Table S2.

### Luciferase and β-galactosidase assay

HS578T cells were co-transfected with *Mock*, *SBE9luc* or *HSF2luc* and a plasmid encoding β-galactosidase. Some HS578T cells were additionally transfected with siRNA targeting HSF1 (L-012109-02-0005, Dharmacon). Cells were treated with assay medium supplemented with 10 ng/ml TGF-β or normal assay medium (Ctrl) for 24 h. After treatments, cells were washed with cold PBS, collected by scraping and the pellets were lysed in Passive Lysis Buffer (E153A, Promega). Next, 3 µl of each sample was pipetted as triplicates on a 96-well plate (CulturPlate, 6005680, Perkin Elmer), and Passive Lysis Buffer was used as blank solution. Before measurement, 18 µl Passive Lysis Buffer was added to each well followed by 90 µl luciferase assay substrate solution (Britelite plus Reporter Gene Assay System, 6066766, Perkin Elmer). Luciferase activity was measured with Hidex Sense Microplate Reader (Type 425-301, Hidex), luminescence IR cut-off 1 s. The luciferase activity was normalized by using β-galactosidase as an internal control for the assay. Briefly, cell lysates were incubated in o-nitrophenyl-β-D-galactopyranoside (ONPG) buffer, containing 4 mg/ml ONPG stock solution, Mg^2+^ buffer and 0.1 M sodium phosphate buffer, at 37°C until the color had turned yellow. β-galactosidase activity was determined by measuring the absorbance at 420 nm using Hidex Sense Microplate Reader (Type 425-301, Hidex).

### qRT-PCR

RNA was isolated with RNeasy Mini Kit (74104, QIAGEN) according to the manufacturer’s instructions. The amount of RNA was quantified using NanoDrop 2000 spectrophotometer (Thermo Fisher Scientific), and for each sample, 900 ng of RNA was reverse transcribed using the iScript cDNA Synthesis Kit (#1708891, Bio-Rad). SensiFAST SYBR Hi-ROX kit (Bioline) was used in the qPCR-reactions, which were performed using a QuantStudio 3 Real-Time PCR system (Applied Biosystems, Thermo Fisher Scientific). The housekeeping gene 18S was used for the normalization of all other genes’ expression. Samples were run in triplicates from three biological replicates. Statistical significance between samples was analyzed in GraphPad Prism 8.3.0 (GraphPad Software) using a paired two-tailed student’s t-test. Primers used for qRT-PCR are indicated in Supplementary Table S1.

### Immunoblotting

Cells were washed in cold PBS (BioWest) after which they were lysed and boiled in 3x Laemmli lysis buffer (30% glycerol, 187.5 mM SDS, 3% Tris-HCl, 0.015% bromophenol blue, 3% β-mercaptoethanol), resolved on a self-cast 8% sodium dodecyl sulphate poly-acrylamide gel (SDS-PAGE) or on 7.5% or 4-20% Mini-PROTEAN TGX Stain-Free Precast Gels (Bio-Rad), and transferred onto a Amersham Protran 0.45 nitrocellulose membrane (Cytiva). For detecting HSF2, the membranes were boiled in MQ-H_2_O for 20 min and blocked with 5% milk-PBS-Tween20. Membranes blotted with phospho-specific antibodies were blocked with 2% BSA-TBS-Tween20. The following antibodies were used for immunoblotting with a 1:1000 dilution: anti-HSF2 (HPA031455, Sigma-Aldrich), anti-HSF1 (ADI-SPA-901, Enzo Life Sciences), anti-HSC70 (ADI-SPA-815, Enzo Life Sciences), anti-β-Tubulin (T8328, Sigma-Aldrich), anti-Snail (C15D3, Cell Signaling Technology), anti-SMAD2/3 (3102, Cell Signaling Technology), anti-pSMAD2 (138D4, Cell Signaling Technology), anti-ORC1 (7A7, BioNordika), anti-MCM2 (D7611, BioNordika), anti-ITGA1 (22146-1-AP, Proteintech), anti-COL3A1 (22734-1-AP, Proteintech), anti-MMP2 (10373-2-AP, Proteintech), anti-CDK1 (A17, ab18, Abcam), anti-CDK-7 (sc-7344, Santa Cruz Biotechnology), and anti-E-cadherin (24E10, Cell Signaling Technology). Horseradish peroxidase-conjugated secondary antibodies were purchased from Promega and GE Healthcare and used with a 1:10,000 dilution. Immunoblots were quantified with FIJI software using the gel analysis tool and the measured values were normalized to respective loading controls. All immunoblots represent three biological replicates unless otherwise indicated.

### Immunofluorescence from cultured cells

For immunofluorescence, 2 × 10^5^ GFP^S^ and 4 × 10^5^ HSF2oe^S^ HS578T cells were seeded on MatTek plates (P35GC-1.5-14-C, MatTek Corporation) 48 h prior to fixation. After 24 h, medium was changed to assay medium only for control samples or to assay supplemented with 10 ng/ml TGF-β for treatments. Following a 24-h treatment, cells were fixed with 4% paraformaldehyde for 10 min, washed three times with PBS, permeabilized with 0.5% Triton X-100 and 3 mM EDTA in PBS for 30 min, and washed again three times with PBS. The cells were then blocked with 10% FBS in PBS for 1 h at RT and incubated with primary antibodies overnight at 4°C. The following primary antibodies were diluted in 10% FBS-PBS: 1:200 anti-COL3A1 (22734-1-AP, Proteintech), 1:100 anti-ITGB1 (12G10, Abcam) and 1:100 anti-vimentin (V6389, Sigma-Aldrich). After primary antibody incubation, cells were washed three times with PBS and incubated with secondary antibodies for 1 h at RT. The following secondary antibodies were diluted in 10% FBS-PBS; 1:500 Alexa Fluor 546 goat anti-rabbit (A11035, Life Technologies), 1:500 Alexa Fluor 555 donkey anti-mouse (A31570, Life Technologies) and 1:500 Alexa Fluor 633 goat anti-mouse (A21052, Invitrogen). For staining F-actin, 1:200 Phalloidin-Atto 647N (65906, Sigma-Aldrich) was used during secondary antibody incubation. Next, the cells were washed once with PBS, incubated with 300 nm DAPI in PBS for 5 min, washed again once with PBS, and covered with VECTASHIELD mounting medium (H-100, Vector Laboratories). Images were taken with a 3i CSU-W1 spinning disk confocal microscope using a 20x objective (Intelligent Imaging Innovations). Acquired images are shown as maximum intensity projections.

### RNA-sequencing

HS578T cells were transiently transfected with plasmids encoding GFP (Mock^T^) or HSF2 (HSF2oe^T^). After a 24-h recovery, the cells were incubated in assay medium supplemented with 10 ng/ml TGF-β or assay medium (Ctrl) for 24 h. Cells were collected, and RNA was purified from the samples using the AllPrep DNA/RNA/miRNA Universal Kit (80224, QIAGEN) according to the manufacturer’s instructions. The amount of RNA was quantified using NanoDrop 2000 spectrophotometer (Thermo Fisher Scientific). Prior to sequencing, the RNA library was prepared according to Illumina stranded mRNA preparation guide (1000000124518). Briefly, poly-A containing mRNA molecules were isolated using poly-T oligo magnetic beads and fragmented with divalent cations under elevated temperatures. The first-strand cDNA synthesis was performed wherein RNA fragments were copied using reverse transcriptase and random primers. In the second-strand cDNA synthesis, dUTP replaced dTTP to achieve strand specificity. Unique dual indexing adapters were ligated to each sample and the quality and concentration of cDNA samples were analyzed with Advanced Analytical Fragment Analyzer and Bioanalyzer 2100 (Agilent, Santa Clara, CA, USA) and Qubit Fluorometric Quantitation (Life Technologies). Samples were sequenced with NovaSeq 6000 S1 v1.5. All the experimental steps after the RNA extraction were conducted at the Finnish Functional Genomics Centre core facility, Turku, Finland. RNA-sequencing was performed from two independent sample series.

### Analysis of RNA-sequencing data

FastQC v0.11.9 was used to confirm the quality of the raw reads and the paired-end reads were aligned to the human genome (primary assembly GRCh38.p13, GENCODE) with STAR version 2.7.9a (Dobin et al., 2013), using the default settings. The number of read pairs mapped to each genomic feature in release 38 of the GENCODE annotation was determined by featureCounts from subread v2.0.1 (Liao et al., 2014). Only read pairs with both ends aligned were counted. Differential gene expression analysis was performed using the Bioconductor R package edgeR (version 3.34.1) (Robinson et al., 2010). Lowly expressed genes were filtered out (using filterByExpr defaults) and the samples were normalized using the trimmed mean of M-values (TMM) method. The threshold for differentially expressed genes was set to FDR < 0.05 and the gene expression data was visualized as an MA plot, produced by the Bioconductor R package Glimma v2.2.0 (Su et al., 2017). Z-score transformed log_2_CPM values were used by the CRAN R package pheatmap v1.0.12 to produce the heat maps. The GO term analysis was performed with topGO v2.44.0.

### Ultra-low attachment assay

MDA-MB-231 cells transiently transfected with plasmids encoding GFP (Mock^T^) or HSF2 (HSF2oe^T^) were collected in assay medium 24 h post-transfection. After counting, 2.7 × 10^3^ cells were placed in each well of a 96-well Ultra-Low Attachment (ULA) plate (#7007, Corning). The total volume in each well was 200 µl. Cells were imaged with Zeiss Axio Vert. A1 microscope (NA 0.4) after 24 h and using a 5x objective. All images were analyzed with ImageJ software (version: 1.53f51) using the BioVoxxel Toolbox plugin (Brocher, 2023). Briefly, the images were converted from 16 bit to 8 bit, a gaussian blur filter with a sigma (radius): 13.00 was applied to reduce background. The perimeter of the spheroids was segmented with the thresholding algorithm of ImageJ choosing the black background option, and the extended particle analyzer option of the BioVoxxel Toolbox plugin was run with default settings.

### Wound healing assay

MDA-MB-231 cells transiently transfected with plasmids encoding GFP (Mock^T^) or HSF2 (HSF2oe^T^) were collected in DMEM medium after a 24 h recovery, and 0.5 × 10^6^ cells were placed in a 12-well plate. After 21 h, a pre-treatment was started by switching to assay medium supplemented with 10 ng/ml TGF-β or normal assay medium for the control cells. Following the 3-h pre-treatment, a scratch was made into the confluent cell sheet by using a 10 µl pipette tip. The rate of wound closure was followed by live cell imaging with a frame interval of 5 min for 24 h using Cell-IQ (Chip-Man Technologies). All images were analyzed with the ImageJ software (version: 1.53f51) using the MRI wound healing tool from https://github.com/MontpellierRessourcesImagerie/imagej_macros_and_scripts with the following settings: method = variance, variance filter radius = 10, threshold = 50, radius open = 4, min. size = 10,000. After running the MRI wound healing tool, we filtered the results to keep the biggest area of each image in the time frame. The wound closure percentage compared to the initial wound area (0 h measurement = 0%) was calculated for all time points. Finally, to correct differences in the time of image acquisition and reduce the segmentation-associated variability, we interpolated the values of wound area with a second-degree polynomial equation.

### Cell Counting Kit-8

The proliferation capacity of Mock^S^ and HSF2oe^S^ cells was assessed using Cell Counting Kit-8 (CCK-8) according to the manufacturer’s instructions (Dojindo, CK04). In brief, cells were seeded at a density of 2 × 10^3^ cells/well in a 96-well plate and let to attach for 24 h before treating with assay medium supplemented with TGF-β (10 ng/ml) for 24 and 72 h. For the longer treatment, fresh medium was changed after 48 h. Following treatments, 10 μl of CCK-8 solution was added to each well of the plate and incubated for 2 h. The absorbance was measured at 450 nm with the Hidex Sense Microplate Reader (Type 425-301, Hidex).

### Atomic force microscopy

The elastic modulus of GFP^S^ and HSF2oe^S^ HS578T cells (Young’s modulus, *E*) was measured using a JPK NanoWizard Atomic Force Microscope (AFM) with a CellHesion module (JPK Instruments) mounted on a Carl Zeiss confocal microscope (Zeiss LSM510) and silicon nitride cantilevers (spring constant: 0.06 Nm^-1^, spherical 4.5 µm tip diameter, Novascan Technologies). The spring constant of the cantilever was calibrated in fluid by using the thermal noise method (Hutter & Bechhoefer, 1993) and a glass substrate was used as an infinitely stiff reference material to determine the deflection sensitivity. The Hertz model of impact was used to determine the elastic properties of cells (Hertz, 1881) with a calibrated force of 4 nN. For assessing the elastic modulus by AFM, 5 × 10^5^ GFP^S^ and 1 × 10^6^ HSF2oe^S^ HS578T cells were seeded on MatTek plates (P35GC-1.5-14-C, MatTek Corporation) 48 h prior to force measurements. Cells were indented in assay medium supplemented with 10 ng/ml TGF-β_1_ or assay medium only at RT. Prior to indentation experiments, a CCD camera mounted on the AFM was used to image the grid of the cell culture dish. The force measurements were performed at four different locations on the culture dish. For each selected location, nine indentation measurements distributed in a 3 × 3 point grid (150 µm × 150 µm^2^) were completed and 5 consecutive measurements were done at each of the nine positions. The elastic modulus for each force curve was calculated using a JPK data processing software (JPK DP version 4.2) predicting a Hertz model of impact. In total, 414 processed indentation curves were used for Mock^S^ Ctrl, 524 for Mock^S^ TGF-β, 400 for HSF2oe^S^ Ctrl, and 443 for HSF2oe^S^ TGF-β. The processed data was summarized using GraphPad Prism 8.3.0 and the bin sizes were determined to set the total number of bins to 24. Fitted curves were represented as a Gaussian distribution or as a sum of two Gaussians. Statistical analysis was done in GraphPad Prism 8.3.0 using a nested two-tailed t-test.

### *In vitro* vasculogenic mimicry assay

Analysis of *in vitro* vasculogenic mimicry assay was used to examine the ability of Mock^S^ and HSF2oe^S^ to form cellular networks in 3D. The cells were pre-treated with TGF-β for 24 h. The wells of a 96-well plate were coated with 50 µl 7 mg/ml Corning Matrigel LDEV-Free Growth Factor Reduced Basement Membrane Matrix (REF 354230, LOT 0266001, Corning) and incubated for 30 min at 37°C to solidify the coating. Next, cells were counted, centrifuged, and resuspended in assay medium only (Ctrl) or in assay medium supplemented with 10 ng/µl TGF-β. A cell suspension of 100 µl containing 4 × 10^4^ cells was added to each Matrigel-coated well. The plate was incubated at 37°C with 5% CO_2_ and each well was imaged after 0 h and 6 h with a Zeiss Axio Vert. A1 microscope (NA 0.4) using a 5x objective. The formation of capillary-like structures was analyzed and quantified from the 6 h images with the ImageJ Software (version: 1.53f51) using the Angiogenesis Analyzer toolset (invert filter, HUVEC Phase Contrast analysis) (Carpentier et al., 2020).

### Organotypic 3D cultures

Organotypic 3D cultures were performed using µ-Slide 15 Well 3D plates (#81506, Ibidi). In brief, 10 µl 7 mg/ml Matrigel was added to the bottom of each well and incubated at 37°C for 1 h. After incubation, 1 × 10^3^ GFP^S^ or HSF2^S^ cells were seeded in the wells in a total medium volume of 20 µl. The slide was then incubated at 37°C for 60 min to allow cell attachment to substrate. Next, the medium was aspirated and 20 µl 7 mg/ml Matrigel was added on top of the cells. After a 30-min incubation, 30 µl medium or medium supplemented with 10 ng/ml TGF-β was added to each well. The medium was changed every 2 days and organotypic tumoroid formation was followed by imaging the wells at day 4, 8, 12, and 16 with Zeiss Axio Vert. A1 microscope using a 20x objective, NA 0.4.

### Zebrafish xenografts

Experiments were performed as previously described in Pekkonen et al., 2018 under the license ESAVI/31414/2020 (granted by Project Authorization Board of Regional State Administrative Agency for Southern Finland) according to the regulations of the Finnish Act on Animal Experimentation (62/2006). Casper zebrafish strain (White et al., 2008) adults were placed in mating tanks and let to spawn overnight. The following day, embryos were collected and cultured in E3 medium (5 mM NaCl, 0.17 mM KCl, 0.33 mM CaCl_2_ 0.33 mM MgSO_4_) supplemented with 0.2 mM phenylthiourea (PTU, Sigma-Aldrich) at 28.5°C. Pronase E was added to culture dishes after 24 h to aid hatching. After 48 h of cultivation, embryos were anesthetized with 200 mg/l MS-222 (Sigma-Aldrich) and placed on agarose for cell transplantation. GFP^S^ and HSF2oe^S^ HS578T cells were trypsinized, washed twice with PBS and resuspended in PBS at a concentration of 1 × 10^6^/10 µl. For immunofluorescence staining and tail vein injection, cells were detached in 5 ml trypsin supplemented with 5 µM CellTracker Green CMFDA (Thermo Fisher Scientific), incubated for at least 5 min, washed twice with PBS and resuspended at a concentration of 1 × 10^6^/10 µl. Nanoinjection into the pericardial cavity and tail vein of embryos was performed using the Nanoject II (Drummond Scientific) and in-house produced capillary glass needles. For each nanoinjection, 4.6 nl of cell suspension was used. Following transplantation, embryos were removed from agarose, placed back on culture dishes, and cultured in E3-PTU at 33°C overnight.

One day post-injection (1dpi), successfully microinjected embryos were selected for experiments, anesthetized as previously described, and moved into a 96-well-plate (one embryo per well). Embryos were imaged using a Nikon Eclipse Ti2 microscope equipped with a Plan UW 2x objective. Imaging was repeated at four days post-injection (4dpi). The GFP intensity of the tumor was measured with FIJI software after background subtraction (rolling ball radius=25, ImageJ version 1.49k) (Schindelin et al., 2012; Schneider et al., 2012) with manual correction. The relative growth of the primary tumor was calculated by dividing the GFP intensity at 4dpi with the GFP intensity at 1dpi. Invading cells in the lens of embryos were not counted due to frequent autofluorescence. The number of invaded cells was counted manually at 1dpi and 4dpi and the fold change was calculated by dividing the obtained value at 4dpi with that of 1dpi. Data was analyzed in GraphPad Prism 8.3.0. using unpaired two-tailed t-test for statistical analyses and results were plotted as mean ± SD. For Mock^S^ 16 and for HSF2oe^S^ 19 zebrafish xenografts were analyzed. Significantly malformed, dead, poorly oriented embryos, wells with two embryos and wells where no GFP signal was detected were excluded from the analysis. Samples were not blinded for imaging or subsequent analyses.

For immunofluorescence staining, zebrafish xenografts were fixed 24 h post-injection in 4% PFA-PBS-0.2% Tween-20 for 2 h at RT. After fixation, animals were washed 4 x 5 min in PBS-0.2% Tween-20, permeabilized in PBS-2% Triton X-100 for 3 h at RT and blocked in PBS-0.2% Triton X-100-5% FBS on rotation at 4 °C overnight. The following day, primary antibody anti-Ki67 (8D5, Cell Signaling Technologies) was diluted in blocking solution 1:800 and the zebrafish were incubated in the primary antibody solution on rotation at 4 °C overnight. The animals were then washed 6 x 30 min in PBS-0.2% Tween-20 at RT, after which secondary antibody solution containing 1:400 Alexa Fluor 633 goat anti-mouse (A21052, Invitrogen) and 300 nM DAPI in blocking solution was added and incubated similarly as the primary antibody. The next day, animals were washed in PBS-0.2% Tween-20 6 x 30 min at RT and mounted on glass bottom dishes using low-melting point agarose. Fixed and stained zebrafish embryos were imaged using 3i CWU Spinning Disk confocal microscope with a 20x objective (Intelligent Imaging Innovations). Acquired images were analyzed with FIJI software. The area of GFP and Ki67 signals were segmented by manual thresholding. The relative area of Ki67 in the tumors was then calculated by dividing the measured area of Ki67 with the area of GFP. Data was analyzed in GraphPad Prism 8.3.0 using unpaired two-tailed t-test for statistical analyses and results were plotted as mean ± SEM. For Mock^S^ 12 and for HSF2oe^S^ 11 zebrafish xenografts were analyzed. Samples were not blinded for imaging or subsequent analyses.

For tail vein injection, successfully microinjected embryos were selected for experiments 24 h post-injection, anesthetized as previously described, and moved into a 96-well-plate (one embryo per well). Embryos were imaged using a Nikon Eclipse Ti2 microscope equipped with a Plan UW 2x objective. The number of formed secondary tumors was calculated manually with FIJI software. Data was analyzed in GraphPad Prism 8.3.0. using unpaired two-tailed t-test for statistical analyses and results were plotted as mean ± SD. For Mock^S^ 25 and for HSF2oe^S^ 29 zebrafish xenografts were analyzed. Samples were blinded for image analysis.

### Human patient tissue samples

Human breast tissue samples were obtained from patients undergoing breast surgery at the Department of Plastic and General Surgery at Turku University Hospital (Turku, Finland). All tissues were voluntarily donated upon written informed consent (Ethical approval ETKM 23/2018) (Table S3). Healthy breast tissue was obtained from female patients undergoing breast reduction mammoplasty surgery. Breast tumor tissue was collected from patients undergoing mastectomy due to pre-invasive (4 samples) or invasive (3 samples) breast carcinoma and were taken by a pathologist after diagnostic sample preparation. All tissues were surgically dissected and processed for fixation. Fixed frozen tissue sections were prepared from donor tissue as previously described (https://doi.org/10.1101/2023.03.12.532249). For immunofluorescence labeling, the frozen sections were first thawed for approximately 30 min at RT, and then blocked and permeabilized in permeabilization buffer (2% BSA, 0.1% Triton X-100 in PBS) for 30 min at RT. Primary antibodies were incubated in 2% BSA in PBS overnight at 4 °C [anti-keratin 8 1:1000 (Hybridoma Bank, TROMA-1); anti-HSF2 1:50 (HPA031455, Sigma-Aldrich); anti-HSF1 1:500 (PA3-017, Invitrogen); anti-Ki67 1:800 (8D5, Cell Signaling Technology)]. After 3 x 10 min washes with PBS, the sections were incubated with the secondary antibodies, diluted in 2% BSA-PBS for 1 h at RT. [anti-rat 488 secondary antibody (H+L) 1:400 (A21208 Thermo Scientific, Alexa Fluor 488); anti-rabbit 568 secondary antibody (H+L) 1:400 (A10042 Thermo Scientific, Alexa Fluor 568); anti-mouse 647 secondary antibody (H+L) 1:400 (A31571 Thermo Scientific, Alexa Fluor 647)]. Finally, the sections were washed 3 x 10 min with PBS, 1:1000 DAPI (D1306 Life Technologies) in the second wash, and then 5 min with MQ-H_2_O. Lastly, the sections were mounted under a #1.5 glass coverslip with Mowiol® (475904 Calbiochem) supplemented with 2.5% 1,4-diazabicyclo[2.2.2]octane (DABCO, D27802 Sigma-Aldrich).

Samples were imaged with the 3i Marianas CSU-W1 spinning disk (50 µm pinholes) confocal microscope using SlideBook 6 software. The objective used was a 20x Zeiss Plan-Apochromat objective (NA 0.8). Signal was detected with the following solid-state lasers and emission filters; 405nm laser, 445/45nm; 488nm laser, 525/30nm; 561nm laser, 617/73nm; 640nm laser, 692/40nm). 3D images were acquired by taking Z-stacks with a 1 µm step size using the Photometrics Prime BSI sCMOS camera with a pixel size of 6.5 μm. Images were 16bit with pixel dimensions 2048×2048.

Image analysis was conducted with ImageJ (version 1.52p). Images of tissue sections were projected as maximum intensity projections. The relative intensity of HSF1 and HSF2 were measured in the area of interest by segmenting the relevant area (keratin 8 for epithelium, DAPI for nuclei) and normalized by subtracting the background intensity. For proliferation analysis, Ki67 signal was segmented, and overlaid on epithelial nuclei to distinguish the fraction of proliferative nuclei. To determine HSF1/2-positive proliferative nuclei, nuclear HSF1/2 signal was segmented and added on the Ki67 mask to create a double positive mask. This was then overlaid on epithelial nuclei to distinguish the fraction of HSF1/2-positive proliferative nuclei.

### Statistical analysis

For qRT-PCR, luciferase reporter assay, ULA assay, CCK-8, *in vitro* vasculogenic mimicry assay, immunofluorescence analysis of zebrafish xenografts, and supplementary RNA-seq data, the statistical significance was analyzed using paired two-tailed student’s t-test and the data was plotted as mean ± SEM (standard error of the mean). For AFM, the statistical significance was analyzed using a nested unpaired two-tailed t-test and the peak values were shown as mean ± SD (standard deviation). The fitted curves represent a Gaussian distribution or a sum of two Gaussians (GraphPad Prism 8.3.0). For the prolonged (1dpi vs 4dpi) xenograft assay and tail vein-injected zebrafish, immunoblotting, and wound healing assay, the statistical significance was tested using an unpaired two-tailed t-test and the data was plotted as mean ± SD. For human patient tissues, data was analyzed in GraphPad Prism 10.1.2. using Mann-Whitney test for statistical analyses, and results were plotted as box plots showing the range of data points (whiskers) and the median value (line).

### Visualization of data

GraphPad Prism 8.3.0 Software (GraphPad Software).

Venn diagrams were generated with BioVenn web application http://www.cmbi.ru.nl/cdd/biovenn/ Schematic illustrations were created with BioRender.com.

## ACKNOWLEDGEMENTS

We thank the patients that voluntarily donated their tissue to this study and the clinical staff of Turku University Hospital. We thank all members of the Sistonen laboratory for their valuable comments and critical review of the manuscript. Dr. Marek Budzynski is acknowledged for his assistance with cloning of the *HSF2luc* plasmid. We thank Dr. Marc L. Mendillo for kindly providing us with the MDA-MB-231 HSF1-KO cell line. Dr. Markus Peurla is acknowledged for his assistance with the image analysis pipelines. We thank Dr. Ilkka Paatero, the Head of Zebrafish Facility at Turku Bioscience Centre, for his assistance with zebrafish xenograft experiments, and Dr. Camilo Guzman and Euro-BioImaging Finland for the assistance with atomic force microscopy. The Finnish Functional Genomics Centre is acknowledged for library preparation and sequencing of RNA-seq samples. The Finnish Functional Genomics Centre is supported by the University of Turku, Åbo Akademi University, and Biocenter Finland. The human tissue samples were prepared at Histocore (Institute of Biomedicine, University of Turku) and microscopy analyses were conducted at The Cell Imaging and Cytometry Core facility (Turku Bioscience, University of Turku and Åbo Akademi University, and Biocenter Finland).

## AUTHOR CONTRIBUTIONS

J.C. Pessa, M.C. Puustinen, P. Hartiala, E. Peuhu, J. Joutsen, and L. Sistonen designed the research; J.C. Pessa, O. Paavolainen, M.C. Puustinen, S. Pihlström, S. Gramolelli, and P. Boström performed the experiments; J.C. Pessa, O. Paavolainen, M.C. Puustinen, H.S.E. Hästbacka, A.J. Da Silva, E. Peuhu, and J. Joutsen analyzed the data; J.C. Pessa, J. Joutsen and L. Sistonen wrote the manuscript with all authors providing feedback.

## FUNDING

This study has been funded by Research Council of Finland (S.G., E.P., L.S.); Sigrid Jusélius Foundation (E.P., L.S.); Cancer Foundation Finland (E.P., L.S.); Jane and Aatos Erkko Foundation (L.S.); Åbo Akademi University Center of Excellence “CellMech” (L.S.); Åbo Akademi University funded Doctoral Position (J.C.P.); Turku Doctoral Program in Molecular Medicine (O.P.) Instrumentarium Science Foundation (M.C.P.); Swedish Cultural Foundation (M.C.P.); Cancer Foundation Southwestern Finland (M.C.P.); Åbo Akademi University Research Foundation (M.C.P.); Turku Doctoral Network in Molecular Biosciences (M.C.P., A.J.D.S.); K. Albin Johansson’s Foundation (M.C.P., A.J.D.S.); Magnus Ehrnrooth Foundation (A.J.D.S.); Otto A. Malm Foundation (A.J.D.S.); The Medical Research Foundation Liv och Hälsa (A.J.D.S.); Finnish Cultural Foundation (A.J.D.S., S.G.); The Finnish Cultural Foundation, Lapland regional fund (J.J.); Lapland hospital district research grant (J.J.); The Hospital District of Southwestern Finland (E.P.).

## COMPETING INTERESTS

Authors declare no competing interests.

## DATA AND MATERIALS AVAILABILITY

The original data is available at Gene Expression Omnibus (GEO) database under accession number GSE211020.

## Notes

### Competing Interest Statement

The authors have declared no competing interest.

## REFERENCES

Abane, R., & Mezger, V. (2010). Roles of heat shock factors in gametogenesis and development. The FEBS journal, 277(20), 4150–4172. 10.1111/j.1742-4658.2010.07830.x

Asghar, M. Y., Lassila, T., Paatero, I., Nguyen, V. D., Kronqvist, P., Zhang, J., Slita, A., Löf, C., Zhou, Y., Rosenholm, J., & Törnquist, K. (2021). Stromal interaction molecule 1 (STIM1) knock down attenuates invasion and proliferation and enhances the expression of thyroid-specific proteins in human follicular thyroid cancer cells. Cellular and molecular life sciences : CMLS, 78(15), 5827–5846. 10.1007/s00018-021-03880-0

Björk, J. K., Åkerfelt, M., Joutsen, J., Puustinen, M. C., Cheng, F., Sistonen, L., & Nees, M. (2016). Heat-shock factor 2 is a suppressor of prostate cancer invasion. Oncogene, 35(14), 1770–1784. 10.1038/onc.2015.241

Brocher, J. (2023) biovoxxel/BioVoxxel-Toolbox: BioVoxxel Toolbox v2.6.0. Zenodo. 10.5281/zenodo.8214743

Cain, S. A., Mularczyk, E. J., Singh, M., Massam-Wu, T., & Kielty, C. M. (2016). ADAMTS-10 and -6 differentially regulate cell-cell junctions and focal adhesions. Scientific reports, 6, 35956. 10.1038/srep35956

Cannell, I. G., Sawicka, K., Pearsall, I., Wild, S. A., Deighton, L., Pearsall, S. M., Lerda, G., Joud, F., Khan, S., Bruna, A., Simpson, K. L., Mulvey, C. M., Nugent, F., Qosaj, F., Bressan, D., CRUK IMAXT Grand Challenge Team, Dive, C., Caldas, C., & Hannon, G. J. (2023). FOXC2 promotes vasculogenic mimicry and resistance to anti-angiogenic therapy. Cell reports, 42(8), 112791. 10.1016/j.celrep.2023.112791

Carpenter, R. L., Sirkisoon, S., Zhu, D., Rimkus, T., Harrison, A., Anderson, A., Paw, I., Qasem, S., Xing, F., Liu, Y., Chan, M., Metheny-Barlow, L., Pasche, B. C., Debinski, W., Watabe, K., & Lo, H. W. (2017). Combined inhibition of AKT and HSF1 suppresses breast cancer stem cells and tumor growth. Oncotarget, 8(43), 73947–73963. 10.18632/oncotarget.18166

Carpentier, G., Berndt, S., Ferratge, S., Rasband, W., Cuendet, M., Uzan, G., & Albanese, P. (2020). Angiogenesis Analyzer for ImageJ - A comparative morphometric analysis of "Endothelial Tube Formation Assay" and "Fibrin Bead Assay". Scientific reports, 10(1), 11568. 10.1038/s41598-020-67289-8

Chen, Y., Di, C., Zhang, X., Wang, J., Wang, F., Yan, J. F., Xu, C., Zhang, J., Zhang, Q., Li, H., Yang, H., & Zhang, H. (2020). Transforming growth factor β signaling pathway: A promising therapeutic target for cancer. Journal of cellular physiology, 235(3), 1903–1914. 10.1002/jcp.29108

Davis, J. E., Nemesure, B., Mehmood, S., Nayi, V., Burke, S., Brzostek, S. R., & Singh, M. (2016). Her2 and Ki67 Biomarkers Predict Recurrence of Ductal Carcinoma in Situ. Applied immunohistochemistry & molecular morphology : AIMM, 24(1), 20–25. 10.1097/PAI.0000000000000223

De Donatis, A., Ranaldi, F., & Cirri, P. (2010). Reciprocal control of cell proliferation and migration. Cell communication and signaling, 8, 20. 10.1186/1478-811X-8-20

de Thonel, A., Ahlskog, J. K., Daupin, K., Dubreuil, V., Berthelet, J., Chaput, C., Pires, G., Leonetti, C., Abane, R., Barris, L. C., Leray, I., Aalto, A. L., Naceri, S., Cordonnier, M., Benasolo, C., Sanial, M., Duchateau, A., Vihervaara, A., Puustinen, M. C., Miozzo, F., … Mezger, V. (2022). CBP-HSF2 structural and functional interplay in Rubinstein-Taybi neurodevelopmental disorder. Nature communications, 13(1), 7002. 10.1038/s41467-022-34476-2

Derynck, R., Turley, S. J., & Akhurst, R. J. (2021). TGFβ biology in cancer progression and immunotherapy. Nature reviews. Clinical oncology, 18(1), 9–34. 10.1038/s41571-020-0403-1

Deville, S. S., & Cordes, N. (2019). The Extracellular, Cellular, and Nuclear Stiffness, a Trinity in the Cancer Resistome-A Review. Frontiers in oncology, 9, 1376. 10.3389/fonc.2019.01376

Dobin, A., Davis, C. A., Schlesinger, F., Drenkow, J., Zaleski, C., Jha, S., Batut, P., Chaisson, M., & Gingeras, T. R. (2013). STAR: ultrafast universal RNA-seq aligner. Bioinformatics (Oxford, England), 29(1), 15–21. 10.1093/bioinformatics/bts635

Dong, B., Jaeger, A. M., Hughes, P. F., Loiselle, D. R., Hauck, J. S., Fu, Y., Haystead, T. A., Huang, J., & Thiele, D. J. (2020). Targeting therapy-resistant prostate cancer via a direct inhibitor of the human heat shock transcription factor 1. Science translational medicine, 12(574), eabb5647. 10.1126/scitranslmed.abb5647

Dong, Q., Xiu, Y., Wang, Y., Hodgson, C., Borcherding, N., Jordan, C., Buchanan, J., Taylor, E., Wagner, B., Leidinger, M., Holman, C., Thiele, D. J., O’Brien, S., Xue, H. H., Zhao, J., Li, Q., Meyerson, H., Boyce, B. F., & Zhao, C. (2022). HSF1 is a driver of leukemia stem cell self-renewal in acute myeloid leukemia. Nature communications, 13(1), 6107. 10.1038/s41467-022-33861-1

Dongre, A., & Weinberg, R. A. (2019). New insights into the mechanisms of epithelial-mesenchymal transition and implications for cancer. Nature reviews. Molecular cell biology, 20(2), 69–84. 10.1038/s41580-018-0080-4

Duncker, B. P., Chesnokov, I. N., & McConkey, B. J. (2009). The origin recognition complex protein family. Genome biology, 10(3), 214. 10.1186/gb-2009-10-3-214

Enserink, J. M., & Kolodner, R. D. (2010). An overview of Cdk1-controlled targets and processes. Cell division, 5, 11. 10.1186/1747-1028-5-11

Fares, J., Fares, M. Y., Khachfe, H. H., Salhab, H. A., & Fares, Y. (2020). Molecular principles of metastasis: a hallmark of cancer revisited. Signal transduction and targeted therapy, 5(1), 28. 10.1038/s41392-020-0134-x

Feng, X. H., & Derynck, R. (2005). Specificity and versatility in tgf-beta signaling through Smads. Annual review of cell and developmental biology, 21, 659–693. 10.1146/annurev.cellbio.21.022404.142018

Fisher R. P. (2005). Secrets of a double agent: CDK7 in cell-cycle control and transcription. Journal of cell science, 118(Pt 22), 5171–5180. 10.1242/jcs.02718

Forsburg S. L. (2004). Eukaryotic MCM proteins: beyond replication initiation. Microbiology and molecular biology reviews, 68(1), 109–131. 10.1128/MMBR.68.1.109-131.2004

Gharibi, A., La Kim, S., Molnar, J., Brambilla, D., Adamian, Y., Hoover, M., Hong, J., Lin, J., Wolfenden, L., & Kelber, J. A. (2017). ITGA1 is a pre-malignant biomarker that promotes therapy resistance and metastatic potential in pancreatic cancer. Scientific reports, 7(1), 10060. 10.1038/s41598-017-09946-z

Gladilin, E., Ohse, S., Boerries, M., Busch, H., Xu, C., Schneider, M., Meister, M., & Eils, R. (2019). TGFβ-induced cytoskeletal remodeling mediates elevation of cell stiffness and invasiveness in NSCLC. Scientific reports, 9(1), 7667. 10.1038/s41598-019-43409-x

Gomez-Pastor, R., Burchfiel, E. T., & Thiele, D. J. (2018). Regulation of heat shock transcription factors and their roles in physiology and disease. Nature reviews. Molecular cell biology, 19(1), 4–19. 10.1038/nrm.2017.73

Hertz, H. (1881). On the contact of elastic solids. Journal fur die Reine und Angewandte Mathematik, 92: 156–171.

Hutter, J. L., & Bechhoefer, J. (1993). Calibration of atomic-force microscope tips. Review of Scientific Instruments, 64: 1868–1873.

Inman, G. J., Nicolás, F. J., Callahan, J. F., Harling, J. D., Gaster, L. M., Reith, A. D., Laping, N. J., & Hill, C. S. (2002). SB-431542 is a potent and specific inhibitor of transforming growth factor-beta superfamily type I activin receptor-like kinase (ALK) receptors ALK4, ALK5, and ALK7. Molecular pharmacology, *62*(1), 65–74. 10.1124/mol.62.1.65

Itoh, Y., Koinuma, D., Omata, C., Ogami, T., Motizuki, M., Yaguchi, S. I., Itoh, T., Miyake, K., Tsutsumi, S., Aburatani, H., Saitoh, M., Miyazono, K., & Miyazawa, K. (2019). A comparative analysis of Smad-responsive motifs identifies multiple regulatory inputs for TGF-β transcriptional activation. The Journal of biological chemistry, 294(42), 15466–15479. 10.1074/jbc.RA119.009877

Jacobs, C., Shah, S., Lu, W. C., Ray, H., Wang, J., Hockaden, N., Sandusky, G., Nephew, K. P., Lu, X., Cao, S., & Carpenter, R. L. (2024). HSF1 Inhibits Antitumor Immune Activity in Breast Cancer by Suppressing CCL5 to Block CD8+ T-cell Recruitment. Cancer research, 84(2), 276–290. 10.1158/0008-5472.CAN-23-0902

Jordà, M., Olmeda, D., Vinyals, A., Valero, E., Cubillo, E., Llorens, A., Cano, A., & Fabra, A. (2005). Upregulation of MMP-9 in MDCK epithelial cell line in response to expression of the Snail transcription factor. Journal of cell science, 118(Pt 15), 3371–3385. 10.1242/jcs.02465

Joutsen, J., & Sistonen, L. (2019). Tailoring of Proteostasis Networks with Heat Shock Factors. Cold Spring Harbor perspectives in biology, 11(4), a034066. 10.1101/cshperspect.a034066

Joutsen, J., Da Silva, A. J., Luoto, J. C., Budzynski, M. A., Nylund, A. S., de Thonel, A., Concordet, J. P., Mezger, V., Sabéran-Djoneidi, D., Henriksson, E., & Sistonen, L. (2020). Heat Shock Factor 2 Protects against Proteotoxicity by Maintaining Cell-Cell Adhesion. Cell reports, 30(2), 583–597.e6. 10.1016/j.celrep.2019.12.037

Joutsen, J., Pessa, J. C., Jokelainen, O., Sironen, R., Hartikainen, J. M., & Sistonen, L. (2024). Comprehensive analysis of human tissues reveals unique expression and localization patterns of HSF1 and HSF2. Cell stress & chaperones, 29(2), 235–271. 10.1016/j.cstres.2024.03.001

Kanchanawong, P., & Calderwood, D. A. (2023). Organization, dynamics and mechanoregulation of integrin-mediated cell-ECM adhesions. Nature reviews. Molecular cell biology, 24(2), 142–161. 10.1038/s41580-022-00531-5

Kashani, A. S., & Packirisamy, M. (2020). Cancer cells optimize elasticity for efficient migration. Royal Society open science, 7(10), 200747. 10.1098/rsos.200747

Kim, B. N., Ahn, D. H., Kang, N., Yeo, C. D., Kim, Y. K., Lee, K. Y., Kim, T. J., Lee, S. H., Park, M. S., Yim, H. W., Park, J. Y., Park, C. K., & Kim, S. J. (2020). TGF-β induced EMT and stemness characteristics are associated with epigenetic regulation in lung cancer. Scientific reports, 10(1), 10597. 10.1038/s41598-020-67325-7

Kmiecik, S. W., & Mayer, M. P. (2022). Molecular mechanisms of heat shock factor 1 regulation. Trends in biochemical sciences, 47(3), 218–234. 10.1016/j.tibs.2021.10.004

Kuivaniemi, H., & Tromp, G. (2019). Type III collagen (COL3A1): Gene and protein structure, tissue distribution, and associated diseases. Gene, 707, 151–171. 10.1016/j.gene.2019.05.003

Kutz, W. E., Wang, L. W., Bader, H. L., Majors, A. K., Iwata, K., Traboulsi, E. I., Sakai, L. Y., Keene, D. R., & Apte, S. S. (2011). ADAMTS10 protein interacts with fibrillin-1 and promotes its deposition in extracellular matrix of cultured fibroblasts. The Journal of biological chemistry, 286(19), 17156–17167. 10.1074/jbc.M111.231571

Lee, J. H., & Massagué, J. (2022). TGF-β in developmental and fibrogenic EMTs. Seminars in cancer biology, 86(Pt 2), 136–145. 10.1016/j.semcancer.2022.09.004

Lei M. (2005). The MCM complex: its role in DNA replication and implications for cancer therapy. Current cancer drug targets, 5(5), 365–380. 10.2174/1568009054629654

Levi-Galibov, O., Lavon, H., Wassermann-Dozorets, R., Pevsner-Fischer, M., Mayer, S., Wershof, E., Stein, Y., Brown, L. E., Zhang, W., Friedman, G., Nevo, R., Golani, O., Katz, L. H., Yaeger, R., Laish, I., Porco, J. A., Sahai, E., Shouval, D. S., Kelsen, D., & Scherz-Shouval, R. (2020). Heat Shock Factor 1-dependent extracellular matrix remodeling mediates the transition from chronic intestinal inflammation to colon cancer. Nature communications, 11(1), 6245. 10.1038/s41467-020-20054-x

Li, J., Chauve, L., Phelps, G., Brielmann, R. M., & Morimoto, R. I. (2016). E2F coregulates an essential HSF developmental program that is distinct from the heat-shock response. Genes & development, 30(18), 2062–2075. 10.1101/gad.283317.116

Liao, Y., Smyth, G. K., & Shi, W. (2014). featureCounts: an efficient general purpose program for assigning sequence reads to genomic features. *Bioinformatics (Oxford*, England), 30(7), 923–930. 10.1093/bioinformatics/btt656

Linares, J., Marín-Jiménez, J. A., Badia-Ramentol, J., & Calon, A. (2021). Determinants and Functions of CAFs Secretome During Cancer Progression and Therapy. Frontiers in cell and developmental biology, 8, 621070. 10.3389/fcell.2020.621070

Liu, Y. L., Chou, C. K., Kim, M., Vasisht, R., Kuo, Y. A., Ang, P., Liu, C., Perillo, E. P., Chen, Y. A., Blocher, K., Horng, H., Chen, Y. I., Nguyen, D. T., Yankeelov, T. E., Hung, M. C., Dunn, A. K., & Yeh, H. C. (2019). Assessing metastatic potential of breast cancer cells based on EGFR dynamics. Scientific reports, 9(1), 3395. 10.1038/s41598-018-37625-0

Lo, P. K., Zhang, Y., Yao, Y., Wolfson, B., Yu, J., Han, S. Y., Duru, N., & Zhou, Q. (2017). Tumor-associated myoepithelial cells promote the invasive progression of ductal carcinoma *in situ* through activation of TGFβ signaling. The Journal of biological chemistry, 292(27), 11466–11484. 10.1074/jbc.M117.775080

Massagué J. (2012). TGFβ signalling in context. Nature reviews. Molecular cell biology, 13(10), 616–630. 10.1038/nrm3434

Massett, M. P., Bywaters, B. C., Gibbs, H. C., Trzeciakowski, J. P., Padgham, S., Chen, J., Rivera, G., Yeh, A. T., Milewicz, D. M., & Trache, A. (2020). Loss of smooth muscle α-actin effects on mechanosensing and cell-matrix adhesions. *Experimental biology and medicine (Maywood*, N.J.), 245(4), 374–384. 10.1177/1535370220903012

Mayer M. P. (2024). Hsf1 and Hsf2 in normal, healthy human tissues: Immunohistochemistry provokes new questions. Cell stress & chaperones, 29(3), 437–439. Advance online publication. 10.1016/j.cstres.2024.04.004

Medici, D., Hay, E. D., & Goodenough, D. A. (2006). Cooperation between snail and LEF-1 transcription factors is essential for TGF-beta1-induced epithelial-mesenchymal transition. Molecular biology of the cell, 17(4), 1871–1879. 10.1091/mbc.e05-08-0767

Mendillo, M. L., Santagata, S., Koeva, M., Bell, G. W., Hu, R., Tamimi, R. M., Fraenkel, E., Ince, T. A., Whitesell, L., & Lindquist, S. (2012). HSF1 drives a transcriptional program distinct from heat shock to support highly malignant human cancers. Cell, 150(3), 549–562. 10.1016/j.cell.2012.06.031

Meng, X., Chen, X., Lu, P., Ma, W., Yue, D., Song, L., & Fan, Q. (2017). miR-202 Promotes Cell Apoptosis in Esophageal Squamous Cell Carcinoma by Targeting HSF2. Oncology research, 25(2), 215–223. 10.3727/096504016X14732772150541

Mezger, V., Rallu, M., Morimoto, R. I., Morange, M., & Renard, J. P. (1994). Heat shock factor 2-like activity in mouse blastocysts. Developmental biology, 166(2), 819–822. 10.1006/dbio.1994.1361

Moreno-Bueno, G., Cubillo, E., Sarrió, D., Peinado, H., Rodríguez-Pinilla, S. M., Villa, S., Bolós, V., Jordá, M., Fabra, A., Portillo, F., Palacios, J., & Cano, A. (2006). Genetic profiling of epithelial cells expressing E-cadherin repressors reveals a distinct role for Snail, Slug, and E47 factors in epithelial-mesenchymal transition. Cancer research, *66*(19), 9543–9556. 10.1158/0008-5472.CAN-06-0479

Mukherjee, P., Winter, S. L., & Alexandrow, M. G. (2010). Cell cycle arrest by transforming growth factor beta1 near G1/S is mediated by acute abrogation of prereplication complex activation involving an Rb-MCM interaction. Molecular and cellular biology, 30(3), 845– 856. 10.1128/MCB.01152-09

Mustafa, D. A., Sieuwerts, A. M., Zheng, P. P., & Kros, J. M. (2010). Overexpression of Colligin 2 in Glioma Vasculature is Associated with Overexpression of Heat Shock Factor 2. Gene regulation and systems biology, 4, 103–107. 10.4137/GRSB.S4546

Paavolainen, O., Peurla, M., Koskinen, L. M., Pohjankukka, J., Saberi, K., Tammelin, E., Sulander, S., Valkonen, M., Mourao, L., Boström, P., Brück, N., Ruusuvuori, P., Scheele, C., Hartiala, P., & Peuhu, E. (2024). Volumetric analysis of the terminal ductal lobular unit architecture and cell phenotypes in the human breast. bioRxiv. 10.1101/2023.03.12.532249

Pasqua, A. E., Sharp, S. Y., Chessum, N. E. A., Hayes, A., Pellegrino, L., Tucker, M. J., Miah, A., Wilding, B., Evans, L. E., Rye, C. S., Mok, N. Y., Liu, M., Henley, A. T., Gowan, S., De Billy, E., Te Poele, R., Powers, M., Eccles, S. A., Clarke, P. A., Raynaud, F. I., … Cheeseman, M. D. (2023). HSF1 Pathway Inhibitor Clinical Candidate (CCT361814/NXP800) Developed from a Phenotypic Screen as a Potential Treatment for Refractory Ovarian Cancer and Other Malignancies. Journal of medicinal chemistry, 66(8), 5907–5936. 10.1021/acs.jmedchem.3c00156

Peinado, H., Quintanilla, M., & Cano, A. (2003). Transforming growth factor beta-1 induces snail transcription factor in epithelial cell lines: mechanisms for epithelial mesenchymal transitions. The Journal of biological chemistry, 278(23), 21113–21123. 10.1074/jbc.M211304200

Pekkonen, P., Alve, S., Balistreri, G., Gramolelli, S., Tatti-Bugaeva, O., Paatero, I., Niiranen, O., Tuohinto, K., Perälä, N., Taiwo, A., Zinovkina, N., Repo, P., Icay, K., Ivaska, J., Saharinen, P., Hautaniemi, S., Lehti, K., & Ojala, P. M. (2018). Lymphatic endothelium stimulates melanoma metastasis and invasion via MMP14-dependent Notch3 and β1-integrin activation. eLife, 7, e32490. 10.7554/eLife.32490

Pessa, J. C., Joutsen, J., & Sistonen, L. (2024). Transcriptional reprogramming at the intersection of the heat shock response and proteostasis. Molecular cell, 84(1), 80–93. 10.1016/j.molcel.2023.11.024

Petojevic, T., Pesavento, J. J., Costa, A., Liang, J., Wang, Z., Berger, J. M., & Botchan, M. R. (2015). Cdc45 (cell division cycle protein 45) guards the gate of the Eukaryote Replisome helicase stabilizing leading strand engagement. Proceedings of the National Academy of Sciences of the United States of America, 112(3), E249–E258. 10.1073/pnas.1422003112

Poulakaki, N., Makris, G. M., Papanota, A. M., Marineli, F., Marinelis, A., Battista, M. J., Boehm, D., Psyrri, A., & Sergentanis, T. N. (2018). Ki-67 Expression as a Factor Predicting Recurrence of Ductal Carcinoma In Situ of the Breast: A Systematic Review and Meta-Analysis. Clinical breast cancer, 18(2), 157–167.e6. 10.1016/j.clbc.2017.12.007

Principe, D. R., Doll, J. A., Bauer, J., Jung, B., Munshi, H. G., Bartholin, L., Pasche, B., Lee, C., & Grippo, P. J. (2014). TGF-β: duality of function between tumor prevention and carcinogenesis. Journal of the National Cancer Institute, 106(2), djt369. 10.1093/jnci/djt369

Puustinen, M. C., & Sistonen, L. (2020). Molecular Mechanisms of Heat Shock Factors in Cancer. Cells, 9(5), 1202. 10.3390/cells9051202

Quintero-Fabián, S., Arreola, R., Becerril-Villanueva, E., Torres-Romero, J. C., Arana-Argáez, V., Lara-Riegos, J., Ramírez-Camacho, M. A., & Alvarez-Sánchez, M. E. (2019). Role of Matrix Metalloproteinases in Angiogenesis and Cancer. Frontiers in oncology, 9, 1370. 10.3389/fonc.2019.01370

Rallu, M., Loones, M., Lallemand, Y., Morimoto, R., Morange, M., & Mezger, V. (1997). Function and regulation of heat shock factor 2 during mouse embryogenesis. Proceedings of the National Academy of Sciences of the United States of America, 94(6), 2392–2397. 10.1073/pnas.94.6.2392

Ridge, K. M., Eriksson, J. E., Pekny, M., & Goldman, R. D. (2022). Roles of vimentin in health and disease. Genes & development, 36(7-8), 391–407. 10.1101/gad.349358.122

Riva, L., Koeva, M., Yildirim, F., Pirhaji, L., Dinesh, D., Mazor, T., Duennwald, M. L., & Fraenkel, E. (2012). Poly-glutamine expanded huntingtin dramatically alters the genome wide binding of HSF1. Journal of Huntington’s disease, 1(1), 33–45.

Robinson, M. D., McCarthy, D. J., & Smyth, G. K. (2010). edgeR: a Bioconductor package for differential expression analysis of digital gene expression data. *Bioinformatics (Oxford*, England), 26(1), 139–140. 10.1093/bioinformatics/btp616

Roos-Mattjus, P., & Sistonen, L. (2022). Interplay between mammalian heat shock factors 1 and 2 in physiology and pathology. The FEBS journal, 289(24), 7710–7725. 10.1111/febs.16178

Sala, A. J., & Morimoto, R. I. (2022). Protecting the future: balancing proteostasis for reproduction. Trends in cell biology, 32(3), 202–215. 10.1016/j.tcb.2021.09.009

Santagata, S., Hu, R., Lin, N. U., Mendillo, M. L., Collins, L. C., Hankinson, S. E., Schnitt, S. J., Whitesell, L., Tamimi, R. M., Lindquist, S., & Ince, T. A. (2011). High levels of nuclear heat-shock factor 1 (HSF1) are associated with poor prognosis in breast cancer. Proceedings of the National Academy of Sciences of the United States of America, 108(45), 18378–18383. 10.1073/pnas.1115031108

Santagata, S., Mendillo, M. L., Tang, Y. C., Subramanian, A., Perley, C. C., Roche, S. P., Wong, B., Narayan, R., Kwon, H., Koeva, M., Amon, A., Golub, T. R., Porco, J. A., Jr, Whitesell, L., & Lindquist, S. (2013). Tight coordination of protein translation and HSF1 activation supports the anabolic malignant state. Science (New York, N.Y.), 341(6143), 1238303. 10.1126/science.1238303

Santopolo, S., Riccio, A., Rossi, A., & Santoro, M. G. (2021). The proteostasis guardian HSF1 directs the transcription of its paralog and interactor HSF2 during proteasome dysfunction. Cellular and molecular life sciences, 78(3), 1113–1129. 10.1007/s00018-020-03568-x

Sasaki, H., Sato, T., Yamauchi, N., Okamoto, T., Kobayashi, D., Iyama, S., Kato, J., Matsunaga, T., Takimoto, R., Takayama, T., Kogawa, K., Watanabe, N., & Niitsu, Y. (2002). Induction of heat shock protein 47 synthesis by TGF-beta and IL-1 beta via enhancement of the heat shock element binding activity of heat shock transcription factor 1. Journal of immunology (Baltimore, Md. : 1950), *168*(10), 5178–5183. 10.4049/jimmunol.168.10.5178

Scherz-Shouval, R., Santagata, S., Mendillo, M. L., Sholl, L. M., Ben-Aharon, I., Beck, A. H., Dias-Santagata, D., Koeva, M., Stemmer, S. M., Whitesell, L., & Lindquist, S. (2014). The reprogramming of tumor stroma by HSF1 is a potent enabler of malignancy. Cell, 158(3), 564–578. 10.1016/j.cell.2014.05.045

Schindelin, J., Arganda-Carreras, I., Frise, E., Kaynig, V., Longair, M., Pietzsch, T., Preibisch, S., Rueden, C., Saalfeld, S., Schmid, B., Tinevez, J. Y., White, D. J., Hartenstein, V., Eliceiri, K., Tomancak, P., & Cardona, A. (2012). Fiji: an open-source platform for biological-image analysis. *Nature methods*, *9*(7), 676–682. 10.1038/nmeth.2019

Schneider, C. A., Rasband, W. S., & Eliceiri, K. W. (2012). NIH Image to ImageJ: 25 years of image analysis. Nature methods, 9(7), 671–675. 10.1038/nmeth.2089

Sengupta, S., Jana, S., & Bhattacharyya, A. (2014). TGF-β-Smad2 dependent activation of CDC 25A plays an important role in cell proliferation through NFAT activation in metastatic breast cancer cells. Cellular signalling, 26(2), 240–252. 10.1016/j.cellsig.2013.11.013

Shaashua, L., Ben-Shmuel, A., Pevsner-Fischer, M., Friedman, G., Levi-Galibov, O., Nandakumar, S., Barki, D., Nevo, R., Brown, L. E., Zhang, W., Stein, Y., Lior, C., Kim, H. S., Bojmar, L., Jarnagin, W. R., Lecomte, N., Mayer, S., Stok, R., Bishara, H., Hamodi, R., … Scherz-Shouval, R. (2022). BRCA mutational status shapes the stromal microenvironment of pancreatic cancer linking clusterin expression in cancer associated fibroblasts with HSF1 signaling. Nature communications, 13(1), 6513. 10.1038/s41467-022-34081-3

Shinde, A. V., Humeres, C., & Frangogiannis, N. G. (2017). The role of α-smooth muscle actin in fibroblast-mediated matrix contraction and remodeling. Biochimica et biophysica acta. Molecular basis of disease, 1863(1), 298–309. 10.1016/j.bbadis.2016.11.006

Smith, R. S., Takagishi, S. R., Amici, D. R., Metz, K., Gayatri, S., Alasady, M. J., Wu, Y., Brockway, S., Taiberg, S. L., Khalatyan, N., Taipale, M., Santagata, S., Whitesell, L., Lindquist, S., Savas, J. N., & Mendillo, M. L. (2022). HSF2 cooperates with HSF1 to drive a transcriptional program critical for the malignant state. Science advances, 8(11), eabj6526. 10.1126/sciadv.abj6526

Strzalka, W., & Ziemienowicz, A. (2011). Proliferating cell nuclear antigen (PCNA): a key factor in DNA replication and cell cycle regulation. Annals of botany, 107(7), 1127–1140. 10.1093/aob/mcq243

Su, S., Law, C. W., Ah-Cann, C., Asselin-Labat, M.-L., Blewitt, M. E., & Ritchie, M. E. (2017). Glimma: interactive graphics for gene expression analysis. *Bioinformatics (Oxford*, England), 33(13), 2050–2052. 10.1093/bioinformatics/btx094

Svitkina T. (2018). The Actin Cytoskeleton and Actin-Based Motility. Cold Spring Harbor perspectives in biology, 10(1), a018267. 10.1101/cshperspect.a018267

Vega, S., Morales, A. V., Ocaña, O. H., Valdés, F., Fabregat, I., & Nieto, M. A. (2004). Snail blocks the cell cycle and confers resistance to cell death. Genes & development, 18(10), 1131–1143. 10.1101/gad.294104

Wang, J., Li, B., Luo, M., Huang, J., Zhang, K., Zheng, S., Zhang, S., & Zhou, J. (2024). Progression from ductal carcinoma in situ to invasive breast cancer: molecular features and clinical significance. Signal transduction and targeted therapy, 9(1), 83. 10.1038/s41392-024-01779-3

White, R. M., Sessa, A., Burke, C., Bowman, T., LeBlanc, J., Ceol, C., Bourque, C., Dovey, M., Goessling, W., Burns, C. E., & Zon, L. I. (2008). Transparent adult zebrafish as a tool for in vivo transplantation analysis. Cell stem cell, 2(2), 183–189. 10.1016/j.stem.2007.11.002

Wilson, G. M., Dinh, P., Pathmanathan, N., & Graham, J. D. (2022). Ductal Carcinoma in Situ: Molecular Changes Accompanying Disease Progression. Journal of mammary gland biology and neoplasia, 27(1), 101–131. 10.1007/s10911-022-09517-7

Xi, C., Hu, Y., Buckhaults, P., Moskophidis, D., & Mivechi, N. F. (2012). Heat shock factor Hsf1 cooperates with ErbB2 (Her2/Neu) protein to promote mammary tumorigenesis and metastasis. The Journal of biological chemistry, 287(42), 35646–35657. 10.1074/jbc.M112.377481

Xu, J., Lamouille, S., & Derynck, R. (2009). TGF-beta-induced epithelial to mesenchymal transition. Cell research, 19(2), 156–172. 10.1038/cr.2009.5

Yang, L. N., Ning, Z. Y., Wang, L., Yan, X., & Meng, Z. Q. (2019). HSF2 regulates aerobic glycolysis by suppression of FBP1 in hepatocellular carcinoma. American journal of cancer research, 9(8), 1607–1621.

Yao, D., Dai, C., & Peng, S. (2011). Mechanism of the mesenchymal-epithelial transition and its relationship with metastatic tumor formation. Molecular cancer research, 9(12), 1608–1620. 10.1158/1541-7786.MCR-10-0568

Zhang, Y., Alexander, P. B., & Wang, X. F. (2017). TGF-β Family Signaling in the Control of Cell Proliferation and Survival. Cold Spring Harbor perspectives in biology, 9(4), a022145. 10.1101/cshperspect.a022145

Zhao, Y. H., Zhou, M., Liu, H., Ding, Y., Khong, H. T., Yu, D., Fodstad, O., & Tan, M. (2009). Upregulation of lactate dehydrogenase A by ErbB2 through heat shock factor 1 promotes breast cancer cell glycolysis and growth. Oncogene, 28(42), 3689–3701. 10.1038/onc.2009.229

Zhong, Y. H., Cheng, H. Z., Peng, H., Tang, S. C., & Wang, P. (2016). Heat shock factor 2 is associated with the occurrence of lung cancer by enhancing the expression of heat shock proteins. Oncology letters, 12(6), 5106–5112. 10.3892/ol.2016.5368

