## Supplementary figures 1-8 and supplementary tables 1-3 for "Dynamic HSF2 regulation drives breast cancer progression by steering the balance between proliferation and invasion"

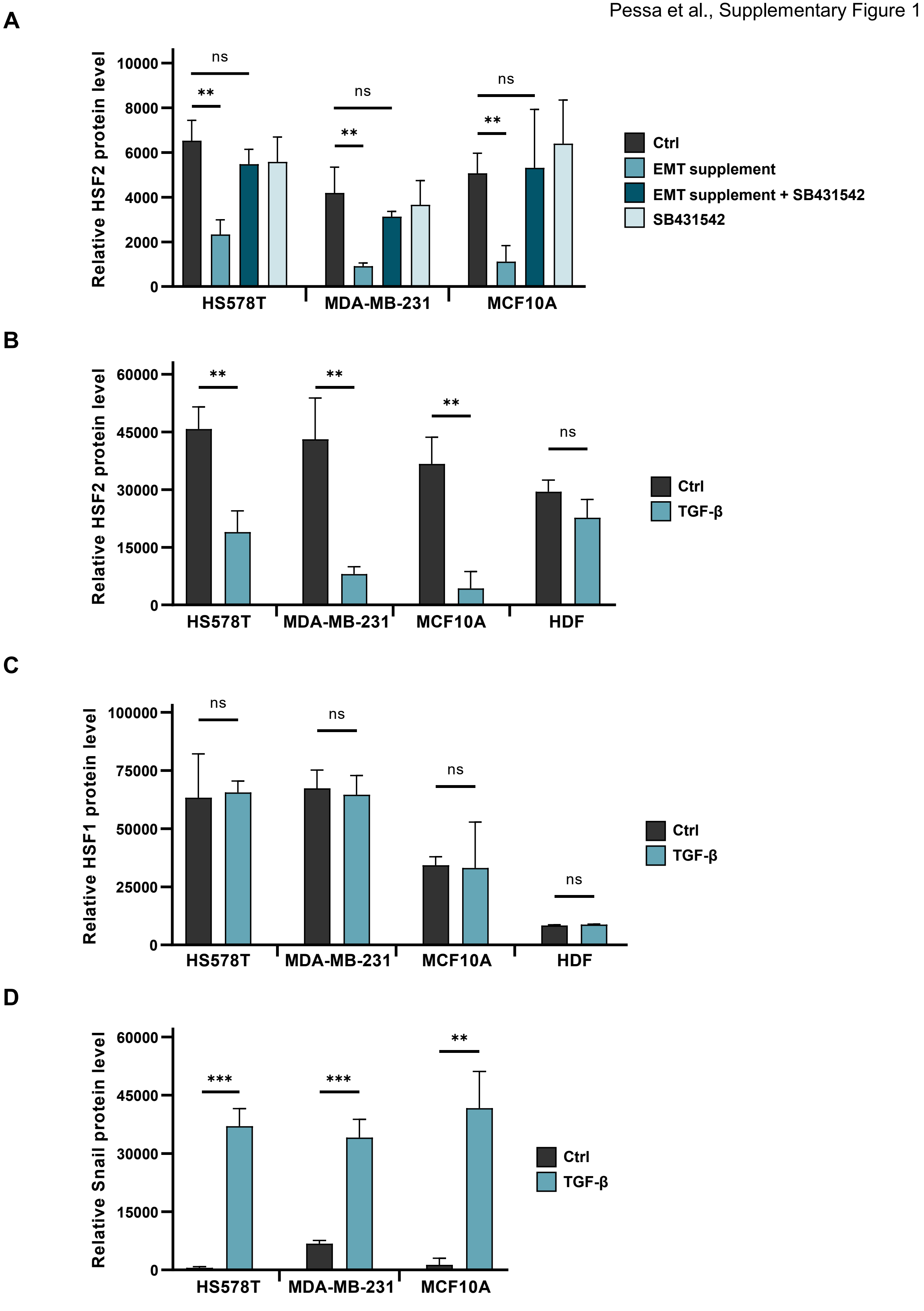
A
Relative HSF2 protein level
10000
8000
6000
4000
2000
0
ns
**
ns
**
ns
**
HS578T
MDA-MB-231
MCF10A
Ctrl
EMT supplement
EMT supplement + SB431542
SB431542
B
Relative HSF2 protein level
60000
45000
30000
15000
0
**
**
**
ns
HS578T
MDA-MB-231
MCF10A
HDF
Ctrl
TGF-β
C
Relative HSF1 protein level
100000
75000
50000
25000
0
ns
ns
ns
ns
HS578T
MDA-MB-231
MCF10A
HDF
Ctrl
TGF-β
D
Relative Snail protein level
60000
45000
30000
15000
0
***
***
**
HS578T
MDA-MB-231
MCF10A
Ctrl
TGF-β

A

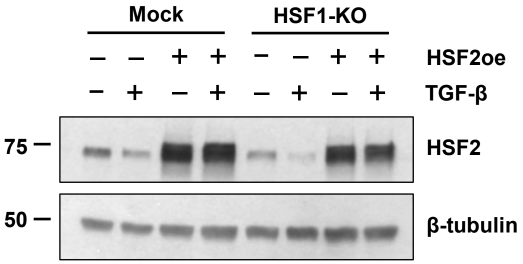
Mock
HSF1-KO
– – + + – – + +
HSF2oe
– + – + – + – +
TGF-β
75
HSF2
50
β-tubulin

B

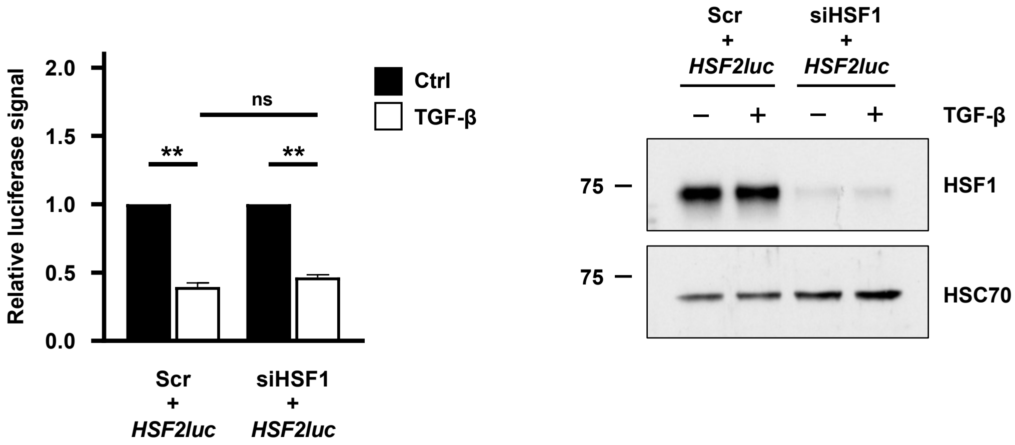
Relative luciferase signal
2.0
1.5
1.0
0.5
0.0
ns
**
**
Ctrl
TGF-β
Scr + HSF2luc
siHSF1 + HSF2luc
Scr + HSF2luc
siHSF1 + HSF2luc
– + – +
TGF-β
75
HSF1
75
HSC70

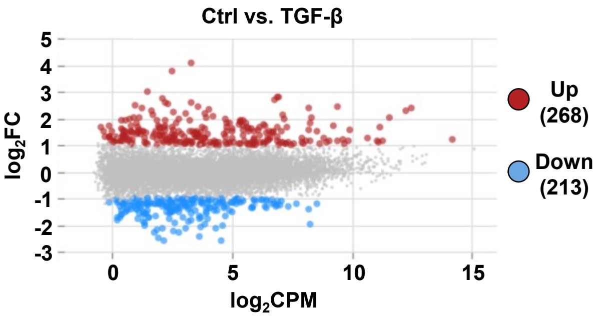
Ctrl vs. TGF-β
log2FC
log2CPM
5
4
3
2
1
0
-1
-2
-3
0
5
10
15
Up
(268)
Down
(213)

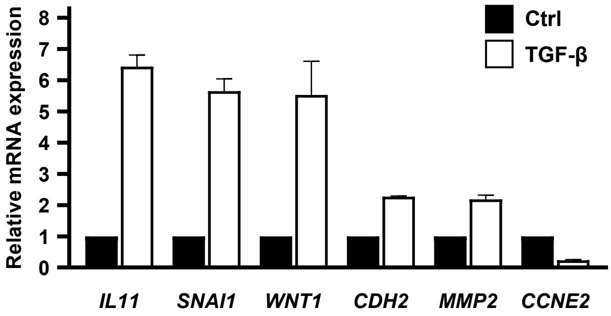
Relative mRNA expression
8
7
6
5
4
3
2
1
0
Ctrl
TGF-β
IL11
SNAI1
WNT1
CDH2
MMP2
CCNE2

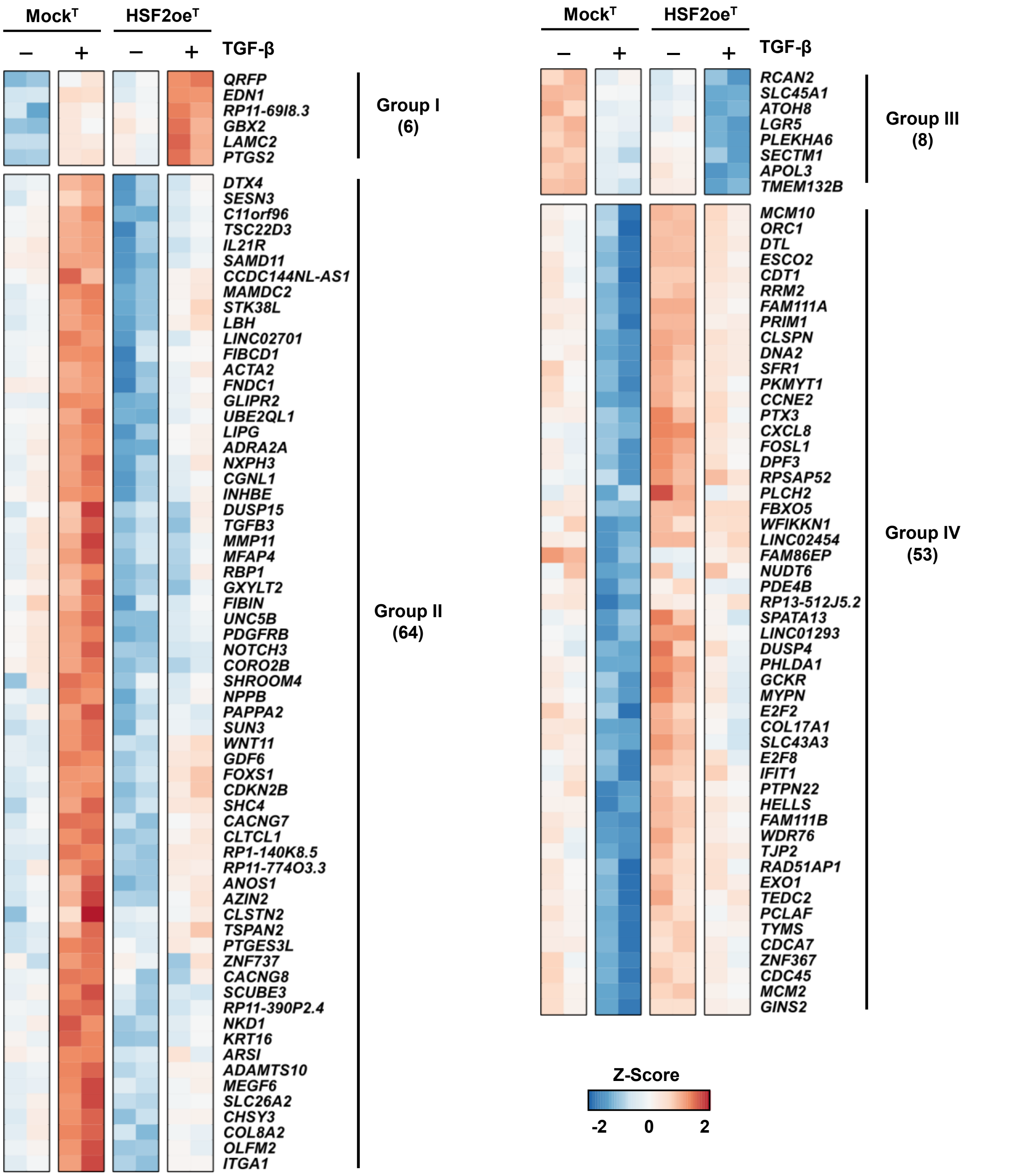

Mock$^T$
HSF2oe$^T$
– + – + TGF-β
QRFP
EDN1
RP11-69I8.3
GBX2
LAMC2
PTGS2
Group I (6)
DTX4
SESN3
C11orf96
TSC22D3
IL21R
SAMD11
CCDC144NL-AS1
MAMDC2
STK38L
LBH
LINC02701
FIBCD1
ACTA2
FNDC1
GLIPR2
UBE2QL1
LIPG
ADRA2A
NXPH3
CGNL1
INHBE
DUSP15
TGFB3
MMP11
MFAP4
RBP1
GXYLT2
FIBIN
UNC5B
PDGFRB
NOTCH3
CORO2B
SHROOM4
NPPB
PAPPA2
SUN3
WNT11
GDF6
FOXS1
CDKN2B
SHC4
CACNG7
CLTCL1
RP1-140K8.5
RP11-774O3.3
ANOS1
AZIN2
CLSTN2
TSPAN2
PTGES3L
ZNF737
CACNG8
SCUBE3
RP11-390P2.4
NKD1
KRT16
ARSI
ADAMTS10
MEGF6
SLC26A2
CHSY3
COL8A2
OLFM2
ITGA1
Group II (64)
Mock$^T$
HSF2oe$^T$
– + – + TGF-β
RCAN2
SLC45A1
ATOH8
LGR5
PLEKHA6
SECTM1
APOL3
TMEM132B
Group III (8)
MCM10
ORC1
DTL
ESCO2
CDT1
RRM2
FAM111A
PRIM1
CLSPN
DNA2
SFR1
PKMYT1
CCNE2
PTX3
CXCL8
FOSL1
DPF3
RPSAP52
PLCH2
FBXO5
WFIKKN1
LINC02454
FAM86EP
NUDT6
PDE4B
RP13-512J5.2
SPATA13
LINC01293
DUSP4
PHLDA1
GCKR
MYPN
E2F2
COL17A1
SLC43A3
E2F8
IFIT1
PTPN22
HELLS
FAM111B
WDR76
TJP2
RAD51AP1
EXO1
TEDC2
PCLAF
TYMS
CDCA7
ZNF367
CDC45
MCM2
GINS2
Group IV (53)
Z-Score
-2 0 2

**A**

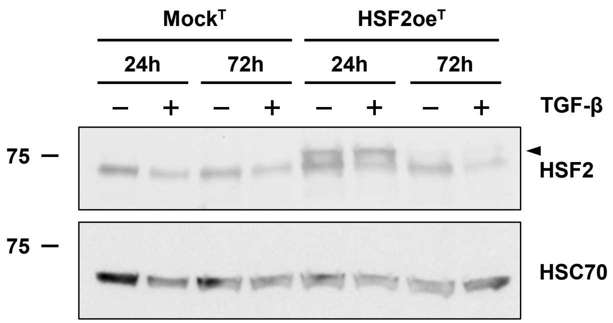
Mock^T
HSF2oe^T
24h
72h
24h
72h
– + – + – + – +
TGF-β
75
HSF2
75
HSC70

**B**

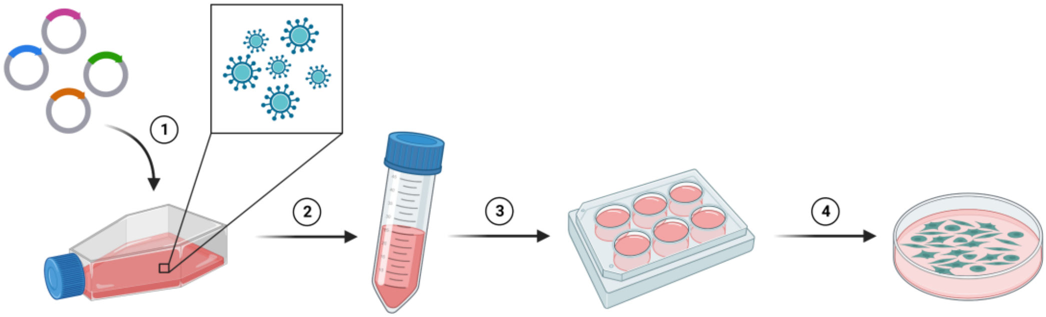
1
2
3
4

**C**

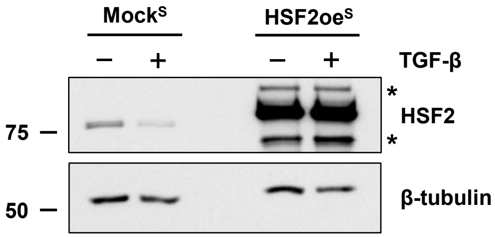
Mock^S
HSF2oe^S
– + – +
TGF-β
*
HSF2
*
75
β-tubulin
50

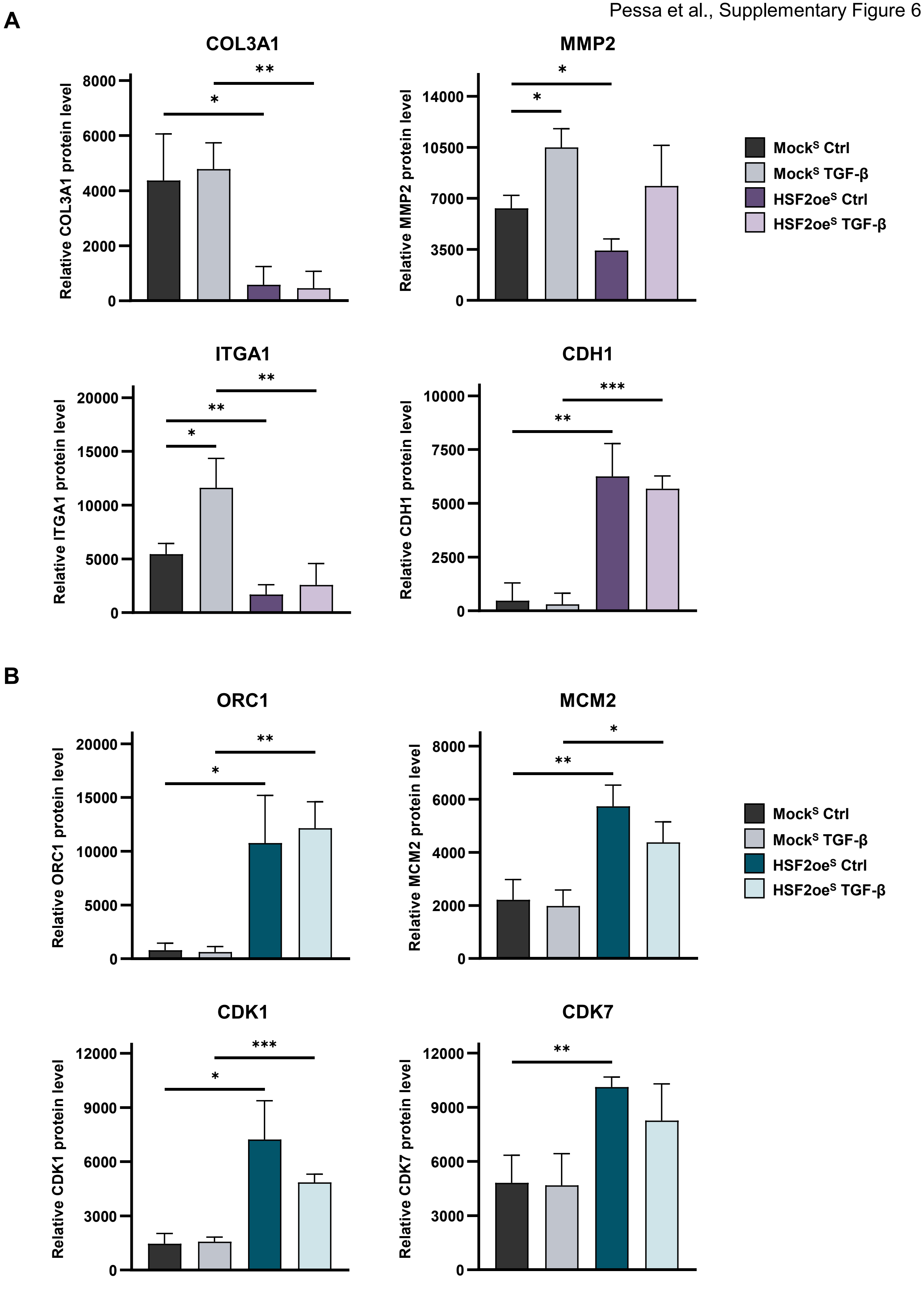
A
COL3A1
Relative COL3A1 protein level
0
2000
4000
6000
8000
*
**
MMP2
Relative MMP2 protein level
0
3500
7000
10500
14000
*
*
Mock^S Ctrl
Mock^S TGF-β
HSF2oe^S Ctrl
HSF2oe^S TGF-β
ITGA1
Relative ITGA1 protein level
0
5000
10000
15000
20000
*
**
**
CDH1
Relative CDH1 protein level
0
2500
5000
7500
10000
**
***
B
ORC1
Relative ORC1 protein level
0
5000
10000
15000
20000
*
**
MCM2
Relative MCM2 protein level
0
2000
4000
6000
8000
**
*
Mock^S Ctrl
Mock^S TGF-β
HSF2oe^S Ctrl
HSF2oe^S TGF-β
CDK1
Relative CDK1 protein level
0
3000
6000
9000
12000
*
***
CDK7
Relative CDK7 protein level
0
3000
6000
9000
12000
**

A

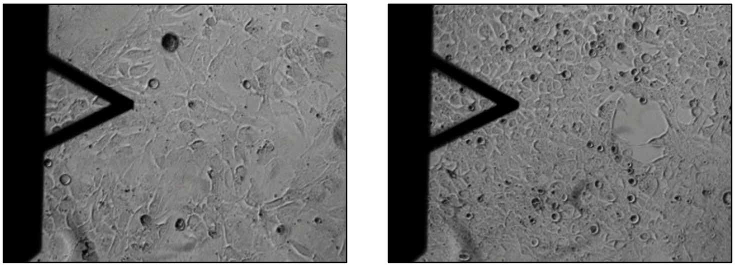

B

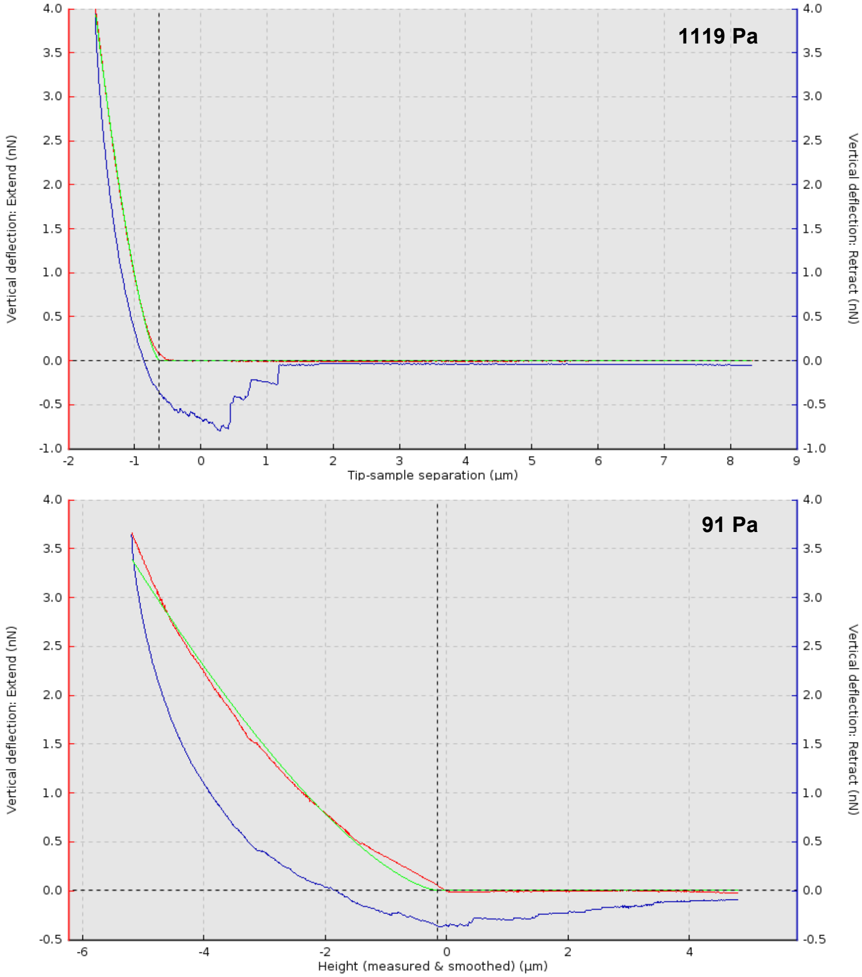
1119 Pa
Vertical deflection: Extend (nN)
Vertical deflection: Retract (nN)
Tip-sample separation (µm)
91 Pa
Vertical deflection: Extend (nN)
Vertical deflection: Retract (nN)
Height (measured & smoothed) (µm)

C

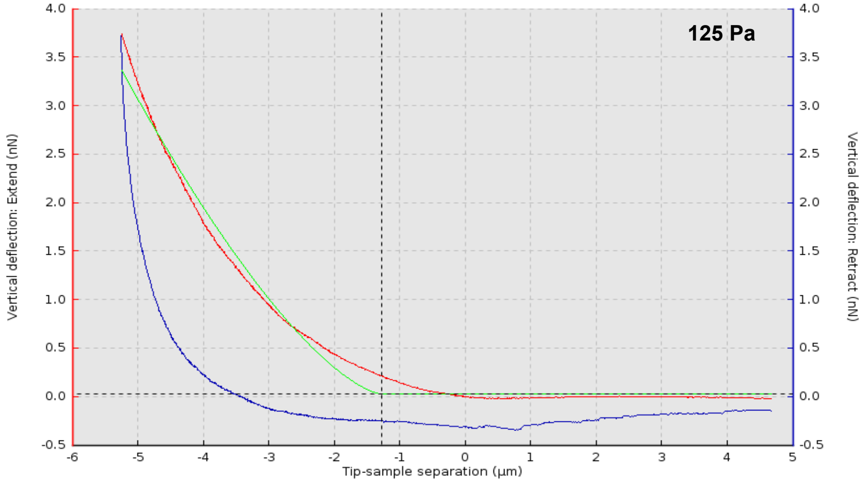
125 Pa
Vertical deflection: Extend (nN)
Vertical deflection: Retract (nN)
Tip-sample separation (µm)

**A**

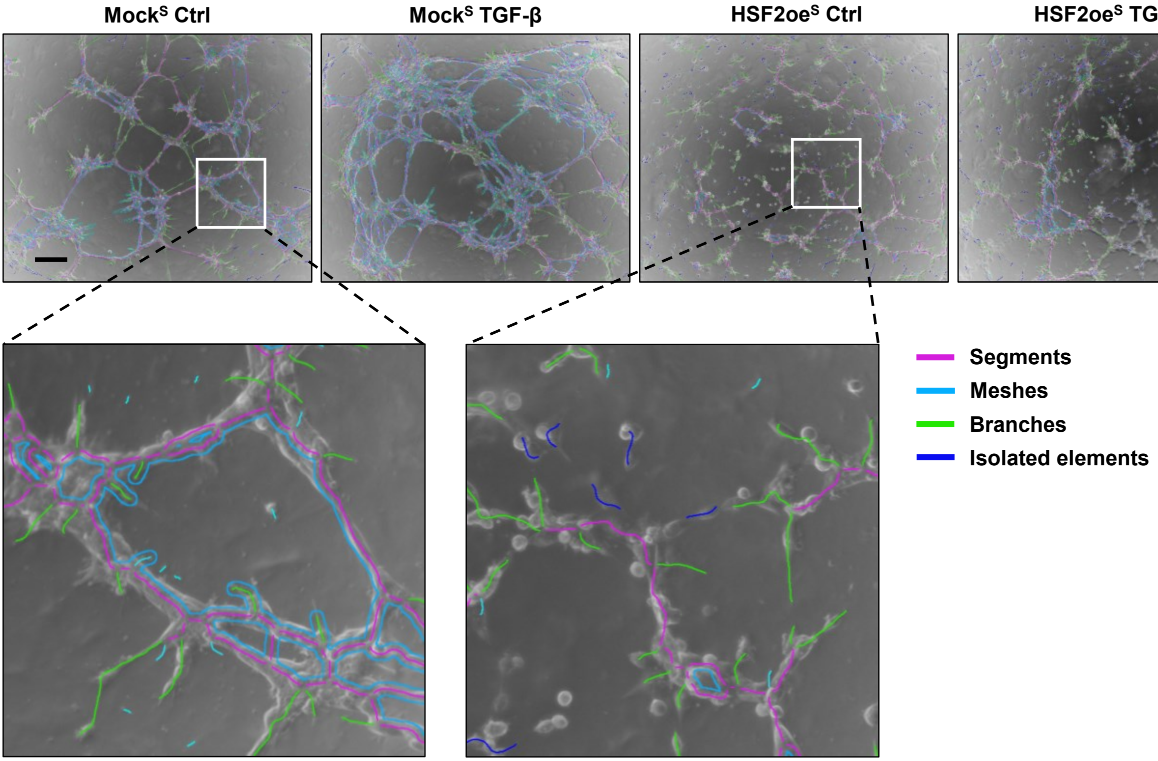


**B**

| Mock$^S$ – | Mock$^S$ + | HSF2oe$^S$ – | HSF2oe$^S$ + | TGF-β |
|---|---|---|---|---|
| 1.00 | 1.34 | 2.20 | 1.53 | Number of isolated segments |
| 1.00 | 0.94 | 1.34 | 1.77 | Total isolated branches length |
| 1.00 | 1.41 | 0.33 | 0.35 | Total mesh area |
| 1.00 | 1.78 | 1.39 | 1.05 | Number of meshes |
| 1.00 | 1.54 | 1.16 | 0.81 | Number of segments |
| 1.00 | 1.54 | 1.21 | 0.91 | Number of nodes |
| 1.00 | 1.62 | 1.22 | 0.91 | Number of master segments |
| 1.00 | 1.11 | 1.25 | 1.08 | Number of extremities |
| 1.00 | 1.35 | 1.11 | 0.97 | Number of pieces |
| 1.00 | 1.34 | 1.14 | 0.89 | Branching interval |
| 1.00 | 1.43 | 1.21 | 1.15 | Number of branches |
| 1.00 | 1.38 | 1.27 | 1.01 | Number of junctions |
| 1.00 | 1.38 | 1.18 | 1.04 | Total length |
| 1.00 | 1.03 | 1.00 | 0.79 | Total branches length |
| 1.00 | 1.00 | 0.99 | 1.00 | Mesh index |
| 1.00 | 1.32 | 1.03 | 0.72 | Total branching length |
| 1.00 | 1.49 | 0.84 | 0.75 | Total master segments length |
| 1.00 | 1.42 | 0.86 | 0.78 | Mean mesh size |
| 1.00 | 1.29 | 0.90 | 0.81 | Number of master junctions |
| 1.00 | 1.34 | 0.80 | 0.83 | Total segments length |

0 1 2

| Target | Sequence |
|---|---|
| **HSF1** | tcgaacacgtggaagctgt |
| | caagctgtggaccctcgt |
| **HSF2** | ggaggaaacccacactaacg |
| | atcgttgctcatccaagacc |
| **CDH2** | agcttctcacggccatacacc |
| | gtgcatgaaggacagcctct |
| **MMP13** | ccagtctccgaggagaaaca |
| | aaaaacagctccgcatcaac |
| **BMP6** | acatggtcatgagctttgtga |
| | actctttgtggtgtcgctga |
| **MMP9** | ttgacagcgacaagaagtgg |
| | gccattcacgtcgtccttat |
| **18S** | gcaattattccccatgaacg |
| | gggacttaatcaacgcaagc |

| *SBEluc* |
|---|
| **1) Amplification of genomic region with MMP9 promoter**<br>tggcttccttactgacggtg<br>tcaaagcctccacaagaccc |
| **2) MMP2 promoter fragment with overhangs**<br>ttggcatcttccatggtggcgactgcagctgctgttgtggggg<br>ctgacgctcagtggaacgaatgagccgagatcacgccac |
| **3) Vector linearization for MMP9 insertion**<br>gccaccatggaagatgccaaaa<br>ttcgttccactgagcgtcagac |
| **4) 4xSBE primer sequence**<br>cgaggtgggaggattgcttggtctagactgccgtctagacttagtacgtctagactgccgtctagacttcgttccactgagcgtcagac<br>(Plasmid #16495, Addgene) |
| **5) Vector linearization for 4xSBE insertion**<br>ttcgttccactgagcgtcagac<br>caagcaatcctcccacct |
| **HSF2oe$^{T}$** |
| **1) HSF2-Myc fragment with overhangs**<br>ctagagtcgcggccgctttacagatcctcttctgagatgagt<br>ctcagatctcgagctcaagcatgaagcagagttcgaacgtg |
| **2) EGFP-N2 vector linearization for HSF2-Myc insertion**<br>gcttgagctcgagatctgagt<br>taaagcggccgcgactcta |
| **3) Strep II primer sequence**<br>tcatctcagaagaggatctgggcagtggagggggatcctggagccacccccagttcgagaagtaaagcggccgcgactctagatc |
| **4) EGFP-N2 vector linearization for Strep II insertion**<br>taaagcggccgcgactcta<br>cagatcctcttctgagatgagt |
| **5) CMV-HSF2-Myc-Strep II-SV40poly(A)tail fragment with overhangs**<br>ccgtaagttatgtaacgcggacgccttaagatacattgatgag<br>cctttttacggttcctggccccgcgttacataacttacgg |
| **6) EGFP-N2 vector linearization for CMV-HSF2-myc-StrepII-SVpoly40(A)tail fragment insertion**<br>ccgcgttacataacttacgg<br>ggccaggaaccgtaaaaaggc |

| Donor | Age (years) | Procedure | Grade |
|---|---|---|---|
| Healthy_1 | 43 | Reduction mammoplasty | N/A |
| Healthy_2 | 18 | Reduction mammoplasty | N/A |
| Healthy_3 | 35 | Reduction mammoplasty | N/A |
| DCIS_1 | 41 | Mastectomy | DCIS grade 2 |
| DCIS_2 | 54 | Mastectomy | DCIS grade 1-2 |
| DCIS_3 | 64 | Mastectomy | DCIS grade 2 |
| DCIS_4 | 64 | Mastectomy | DCIS grade 2 |
| IDC_1 | 85 | Mastectomy | IDC grade 1-2 |
| IDC_2 | 77 | Mastectomy | IDC grade 2 |
| IDC_3 | 88 | Tumor resection | IDC grade 2 |

### SUPPLEMENTARY FIGURE LEGENDS

**Supplementary figure 1. HSF2 is downregulated in response to TGF-β signaling**

**(A)** Quantification of the relative HSF2 protein level from immunoblot analysis in Fig. 1A. Results were normalized to the loading control and presented as mean ± SD, ** = p-value < 0.01, ns = not significant.

**(B)** Quantification of the relative HSF2 protein level from immunoblot analysis in Fig. 1B. Results were normalized to the loading control and presented as mean ± SD, ** = p-value < 0.01, ns = not significant.

**(C)** Quantification of the relative HSF1 protein level from immunoblot analysis in Fig. 1B. Results were normalized to the loading control and presented as mean ± SD, ns = not significant.

**(D)** Quantification of the relative Snail protein level from immunoblot analysis in Fig. 1D. Results were normalized to the loading control and presented as mean ± SD, ** = p-value < 0.01, *** = p-value < 0.001.

**Supplementary figure 2. HSF1 is not directly involved in the TGF-β-mediated downregulation of HSF2**

**(A)** Immunoblot analysis of HSF2 protein levels in an MDA-MB-231 control cell line (Mock) and in a stable HSF1 knock-out cell line (HSF1-KO) (Smith et al., 2022). Cell lines were transiently transfected with plasmids encoding ectopic HSF2 and treated with 10 ng/ml TGF-$\beta_1$ or assay medium for 24 h. β-tubulin was used as a loading control.

**(B)** Analysis of the luciferase reporter gene activity in HS578T cells co-transfected with β-galactosidase, *HSF2luc* and with scrambled siRNA (Scr) or siRNA targeting HSF1 (siHSF1). At 24 h post-transfection, cells were treated with 10 ng/ml TGF-$\beta_1$ or assay medium (Ctrl) for 24 h. The luciferase signal was normalized to β-galactosidase activity and is shown relative to the Ctrl sample. Results were plotted as mean ± SEM, ** = p-value $\leq$ 0.01, ns = not significant. All data represents three biological replicates. Silencing of HSF1 was verified by immunoblotting and HSC70 was used as a loading control.

**Supplementary figure 3. Activation of the TGF-β signaling pathway induces expression of well-known TGF-β target genes**

RNA-seq analysis of gene expression profile in HS578T cells treated with 10 ng/ml TGF-$\beta_1$ or assay medium (Ctrl) for 24 h. In the MA plot, the total number of upregulated (red) and downregulated (blue) genes is indicated in brackets. Genes were considered differentially expressed if the $\log_2$FC was at least ± 1. Relative mRNA expression of a subset of well-known TGF-β-responsive genes: *IL11*, *SNAI1*, *WNT1*, *CDH2*, *MMP2*, and *CCNE2*. The data is presented as mean ± SEM and shown relative to the Ctrl sample.

**Supplementary figure 4. HSF2 disrupts the expression of genes involved in TGF-β-induced cellular responses**

Heat map of the differentially expressed genes in Groups I, II, III, and IV (Fig. 2D). Group I: genes upregulated by both TGF-β and exogenous HSF2, Group II: genes upregulated by TGF-β but impaired response in the presence of exogenous HSF2, Group III: genes downregulated by both TGF-β and exogenous HSF2, and Group IV: genes downregulated by TGF-β and impaired response in the presence of exogenous HSF2. The number of genes in each group is indicated in brackets. The Z-score was calculated based on the $log_2$CPM values for each gene. The double columns for each condition represent the two biological replicates from the RNA-seq.

**Supplementary figure 5. Expression of ectopic HSF2 is sustained in TGF-β-treated stable cell lines**

**(A)** Immunoblot analysis of HSF2 protein levels in HS578T cells transiently transfected with plasmids encoding GFP ($Mock^T$) or HSF2 ($HSF2oe^T$) and treated with 10 ng/ml TGF-$\beta_1$ or assay medium for 24 or 72 h. HSC70 was used as a loading control. Arrowhead (◀) denotes exogenous HSF2.

**(B)** Schematic illustration of the generation of stable cell lines expressing GFP ($Mock^S$) and HSF2 ($HSF2oe^S$). **1)** HEK-293T cells were transfected with plasmids encoding GFP or HSF2, and three separate plasmids encoding fundamental parts of the lentivirus. Post-transfection, cells produce functionally active virus particles. **2)** Viral medium was collected and filter-purified. **3)** Viral medium was applied to HS578T cells. **4)** Post-transduction, cells were cultured under puromycin selection. For details, see Materials and Methods.

**(C)** Immunoblot analysis of HSF2 protein levels in $Mock^S$ and $HSF2oe^S$ cells treated with 10 ng/ml TGF-$\beta_1$ or assay medium for 72 h. β-tubulin was used as a loading control. Asterisk (*) denotes unspecific bands.

**Supplementary figure 6. Ectopic HSF2 suppresses the expression of ECM-related proteins and induces the expression of cell cycle regulators**

**(A)** Quantification of the relative COL3A1, MMP2, ITGA1, and CDH1 protein levels from immunoblot analysis in Fig. 3B. Results were normalized to the loading control and presented as mean ± SD, * = $p$-value $< 0.05$, ** = $p$-value $< 0.01$.

**(B)** Quantification of the relative ORC1, MCM2, CDK1, and CDK7 protein levels from immunoblot analysis in Fig. 6B. Results were normalized to the loading control and presented as mean ± SD, * = $p$-value $< 0.05$, ** = $p$-value $< 0.01$, *** = $p$-value $< 0.001$.

**Supplementary figure 7. Forced expression of HSF2 reduces cellular stiffness**

**(A)** Representative images of selected positions for AFM indentation measurements ($Mock^S$ left, $HSF2oe^S$ right). A CCD camera mounted on the AFM was used for image acquisition.

**(B)** Representative force curves from AFM indentation measurements of $Mock^S$ cells. Due to a wide distribution in measured stiffnesses, a stiff (1119 Pa) and a soft (91 Pa) force curve is shown. Elastic modulus for each force curve was calculated using a JPK data processing software (JPK DP version 4.2) predicting a Hertz model of impact.

**(C)** A representative force curve from AFM indentation measurement of HSF2oe$^S$ cells (125 Pa). Elastic modulus for the force curve was calculated using a JPK data processing software (JPK DP version 4.2) predicting a Hertz model of impact.

**Supplementary figure 8. Analysis of capillary-like network formation in *in vitro* vasculogenic mimicry assay**

**(A)** Representative images of capillary-like network segmentation. Mock$^S$ and HSF2oe$^S$ cells were treated with 10 ng/ml TGF-$\beta_1$ or assay medium (Ctrl) for 24 h prior to initiation of the assay. Following the pre-treatment, $4 \times 10^4$ cells were seeded on Matrigel and treated with 10 ng/ml TGF-$\beta_1$ or assay medium (Ctrl). Images were taken at 6 h with Zeiss Axio Vert. A1 microscope using a 5x objective, NA 0.4. Scale bar 200 µm. The capillary-like structures were analyzed and quantified with ImageJ Software (version: 1.53f51) using the Angiogenesis Analyzer plugin toolset and the "Analyze HUVEC phase contrast" command. Insets of Mock$^S$ Ctrl and HSF2oe$^S$ Ctrl (white rectangles) denote selected parameters of vascular structure segmentation: blue = meshes, magenta = segments, and green = branches, which are quantified in Fig. 5B. Dark blue indicates isolated elements that are short fragments not attached to other structures.

**(B)** Heat map of the parameters from quantification of capillary-like networks. The value for each parameter was normalized relative to the value of Mock$^S$ Ctrl. The data represents three biological replicates.

**Supplementary table 1. Primers used for qRT-PCR (5' – 3').**

**Supplementary table 2. Primers used for plasmid construction (5' – 3').**

**Supplementary table 3. Characteristics of patient tissue samples.** Ductal carcinoma *in situ* (DCIS), invasive ductal carcinoma (IDC).
